# Evolution of protective human antibodies against *Plasmodium falciparum* circumsporozoite protein repeat motifs

**DOI:** 10.1101/798769

**Authors:** Rajagopal Murugan, Stephen W. Scally, Giulia Costa, Ghulam Mustafa, Elaine Thai, Tizian Decker, Alexandre Bosch, Katherine Prieto, Elena A. Levashina, Jean-Philippe Julien, Hedda Wardemann

## Abstract

Circumsporozoite protein of the human malaria parasite *Plasmodium falciparum* (PfCSP) is the main target of antibodies that prevent the infection and disease. Protective antibodies recognize the central PfCSP domain, but our understanding of how parasite inhibition is associated with recognition of this domain and with the evolution of potent antibodies remains scattered. Here, we characterized the epitope specificity of 200 human monoclonal PfCSP antibodies. We show that the majority of PfCSP antibodies bind to NANP and NANP-like motifs with different preferences and define the molecular basis for recognition. Epitope cross-reactivity evolved with increasing antibody affinity around a conserved (N/D)P-NANP-N(V/A) core. High affinity to this motif, but not binding to NANP-like motifs, was associated with parasite inhibition and protection. Thus, NANP drives the development of potent PfCSP antibodies independently of their cross-reactivity profile, a finding with direct implications for the design of a second-generation PfCSP-based malaria vaccine.

**HIGHLIGHTS:**

- The majority of human PfCSP antibodies recognize multiple epitopes
- NANP affinity maturation drives the evolution of cross-reactive PfCSP antibodies
- Preferential PfCSP antibody binding to a conserved core motif
- High affinity not epitope specificity is associated with PfCSP antibody potency

## INTRODUCTION

*Plasmodium falciparum* (Pf) is a unicellular apicomplexan parasite that causes malaria, a life-threatening vector-borne disease. Pf sporozoites, the parasite stage that is transmitted to humans by infectious *Anopheles* mosquitoes, are densely covered by circumsporozoite protein (PfCSP). PfCSP plays an essential role in parasite development in the mosquito vector and establishment of the infection in the human host (Coppi et al., 2011; Frevert et al., 1993; Ménard et al., 1997). It consists of an amino (N) terminal domain, a disordered central region made up of only five amino acids (aa; asparagine (N), alanine (A), valine (V), aspartate (D), proline (P)) arranged in repeating NANP and a few interspersed NVDP sequence motifs, as well as a carboxy (C) terminal domain that anchors the protein to the sporozoite surface by a GPI anchor (Figure 1A, (Dame et al., 1984; Enea et al., 1984)). In contrast to the C terminus that harbors substantial sequence diversity, the repeating motifs in the central domain are fully conserved in their aa sequences and show variability only in their numbers (Bailey et al., 2012; Escalante et al., 2002).

**Figure 1.**
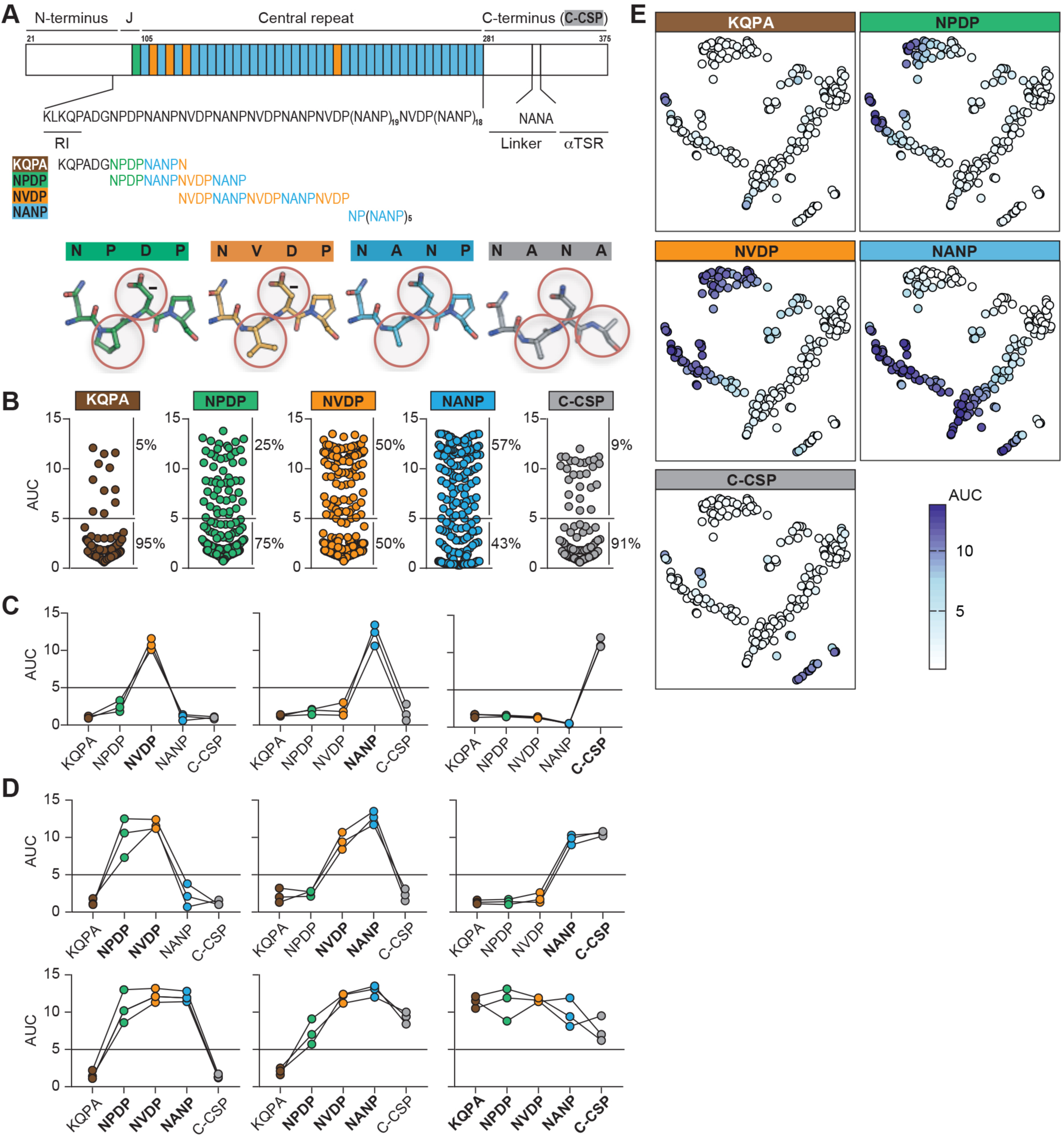
Cross-reactivity of human PfCSP-reactive antibodies. (A) Schematic representation of PfCSP (NF54) comprising the N-terminus, central repeat, and C-terminal (C-CSP) domain. The amino acid sequence downstream of the N-terminal domain including the conserved region 1 (RI, (Dame et al., 1984)), the N-terminal junction, and the central repeat domain is indicated, as well as the NANA epitope in the linker region upstream of the aTSR domain in C-CSP. Amino acid sequences of overlapping peptides in the N-terminal junction containing known epitopes of protective antibodies (Kisalu et al., 2018; Tan et al., 2018), designated as KQPA (brown), NPDP (green), NVDP (orange), and of a NANP 5.5-mer peptide (designated as NANP, blue) are indicated and color-coded. NPDP, NANP, NVDP and NANA motifs are highlighted and depicted as stick structures. (B) Binding strength of anti-PfCSP antibodies (n=200; Table S1; (Murugan et al., 2018)) to the indicated overlapping peptides and C-CSP as in (A) is shown as calculated area under the curve (AUC) values based on ELISA measurements at different antibody concentrations (Figure S1). The frequency of reactive and non-reactive antibodies is indicated. (C and D) Binding profile to the indicated PfCSP peptides and C-CSP as in (B) of representative antibodies with specificity for NVDP, NANP, or C-CSP (C) and cross-reactive antibodies with different binding profiles (D). (E) t-SNE clustering-based illustration of the ELISA binding strength to the indicated peptides as determined by AUC for all PfCSP-reactive antibodies (n=200). Data in B-E shows mean values from three independent experiments. Horizontal lines in B-D indicate the reactivity threshold. See also Figure S1 and Table S1.

The repeating NANP motif represents the major PfCSP B cell epitope on sporozoites and induces dominant serum antibody responses (Dame et al., 1984; Enea et al., 1984; Zavala et al., 1983). Based on historic observations that antibodies against the central repeat inhibit Pf *in vitro* and can protect from *Plasmodium* infection in animal models, a PfCSP-based malaria vaccine (RTS,S) has been developed (Cohen et al., 2010; Nardin et al., 1982; Potocnjak et al., 1980; Yoshida et al., 1980). RTS,S contains a truncated version of PfCSP composed of 18.5 NANP repeats and the PfCSP C-terminal domain. To increase immunogenicity, the truncated PfCSP protein is fused to Hepatitis B surface antigen (HBsAg) and co-expressed with free HBsAg, assembling into virus-like particles. A large pediatric Phase III clinical trial with RTS,S in AS01, a potent liposome-based adjuvant, has been performed at eleven malaria endemic sites in seven African countries (The RTS,S Clinical Trials Partnerships 2011, 2012, 2015). Although RTS,S-induced anti-NANP antibody titers associated with clinical protection, the overall efficacy of the vaccine even after a booster immunization was below 40% within two years and therefore, needs to be improved (Olotu et al., 2016).

A handful of studies have recently identified potent human monoclonal PfCSP antibodies against the central region, but not the N-or C-terminal domains (Foquet et al., 2014; Murugan et al., 2018; Oyen et al., 2017; Scally et al., 2018; Triller et al., 2017). In addition, protective B cell epitopes in the short amino acid (aa) junction that links the PfCSP N terminus with the NANP-, NVDP-repeat region and contains a single highly similar NPDP sequence motif were reported (Figure 1A; (Kisalu et al., 2018; Tan et al., 2018)). Data from a small number of recombinant antibodies with reactivity to these epitopes suggest that recognition of the N-terminal junction might be associated with more potent parasite inhibition than binding to the repeat region, a finding of great importance for the design of an improved second generation PfCSP-based vaccine (Kisalu et al., 2018; Tan et al., 2018). However, all antibodies with reactivity to the N-terminal junction showed at least some degree of NANP cross-reactivity and recognized more than one epitope on full-length PfCSP. Thus, whether the N-terminal junction represents a better antibody target than the NANP repeat, what the molecular basis is for antibody cross-reactivity with different PfCSP epitopes, and how antibodies against the junction evolve remains elusive. Here, we addressed these questions by performing extensive structural and functional analyses on a large panel of human PfCSP-reactive monoclonal antibodies (mAbs). The data identify a conserved recognition mode of the most potent Pf-inhibitory antibodies around NANP motifs and demonstrate that cross-reactivity with the N-terminal junction is associated with affinity maturation to NANP. Independently of the cross-reactivity profile, we show that high PfCSP affinity is a prerequisite for antibody potency but not sufficient to predict the parasite-inhibitory capacity.

## RESULTS

### The majority of PfCSP antibodies are cross-reactive with several epitopes

To identify antibodies with reactivity to the different subdomains, we analyzed a large panel of 200 recombinant PfCSP-reactive human mAbs (Murugan et al., 2018) for binding to the junction, central repeat and C-terminal region of PfCSP by ELISA (Figure 1; Table S1). The antibodies were derived from PfCSP-reactive memory B cells of malaria-naïve volunteers after repeated immunization with live sporozoites under chemoprophylaxis (PfSPZ-CVac; (Mordmüller et al., 2017)). Binding was determined to three overlapping peptides in the N-terminal junction containing highly similar NPDP, NVDP, and NANP aa motifs, to a NP(NANP)_5_ peptide representative of the central repeat, and to the complete C-terminus (C-CSP) with a unique NANA sequence (Figures 1A and S1). The N-terminal junction peptides, here abbreviated as KQPA, NPDP, and NVDP according to the first four aa of each peptide, covered aa 95-109 (KQPADGNPDPNANPN), aa 101-116 (NPDPNANPNVDPNANP), and aa 109-125 (NVDPNANPNVDVNANPNVDP), respectively. Nearly 80% of the antibodies (155/200) bound at least one of the peptides above our cut-off with an ELISA area under the curve (AUC) of >5. The vast majority bound NANP (57%), as previously described (Murugan et al., 2018) and NVDP (50%). Antibodies with reactivity to NPDP (25%) or C-CSP (9%) were less abundant, and only a few recognized KQPA (5%), representing the most N-terminal part of the junction (Figure 1B). Several of the 155 antibodies recognized only the NANP repeat (26%), the NVDP peptide (11%), or the C terminus (3%) but not KQPA, or NPDP and were therefore defined as epitope-specific (Figure 1C). In contrast, 92/155 antibodies (59%) recognized two or more PfCSP peptides with distinct binding profiles and were therefore referred to as epitope cross-reactive (Figure 1D). The vast majority (76%) of these cross-binders interacted strongly with the NANP repeat, whereas 24% lacked NANP-reactivity and instead showed preferential binding to NVDP and NPDP. Presumably due to the low degree of sequence similarity between the C terminus and the N-terminal junction, C-CSP binding was lower in C-CSP-reactive antibodies with cross-reactivity to the repeat and the N-terminal junction peptides compared to C-CSP-reactive antibodies that only cross-reacted with NANP or compared to C-CSP specific antibodies. A t-distributed stochastic neighbor embedding (t-SNE) analysis visualizes the binding profile of all antibodies to the individual peptides and to C-CSP (Figure 1E). In summary, we identified a high number of cross-reactive human PfCSP antibodies with a wide spectrum of antigen-binding profiles to the N-terminal junction, the central repeat and C-CSP.

### Cross-reactivity with the N-terminal junction correlates with antibody affinity to NANP

To delineate the link between cross-reactivity and binding strength, we measured the affinity of antibodies with ELISA-reactivity to the different PfCSP peptides and C-CSP by surface plasmon resonance (SPR; Figures 2 and S2). The data confirmed the high degree of antibody cross-reactivity observed in the ELISA. Anti-NVDP antibodies with cross-reactivity to the overlapping NPDP peptide showed, on average, significantly higher NVDP affinities compared to NVDP-specific antibodies (Figure 2A). Similarly, anti-NANP antibodies with cross-reactivity to the N-terminal junction had higher NANP affinities than NANP-specific antibodies (Figure 2B). Thus, NANP affinity increased with cross-reactivity to the N-terminal junction. The gain in NANP affinity was paralleled by a comparable increase in affinity to NVDP and NPDP (Figure 2B). Highest mean affinities to NANP, NVDP and NPDP were observed in cross-reactive antibodies that bound all four peptides, although their KQPA affinity was overall low compared to the other peptides. In contrast, strong binding to C-CSP was not associated with cross-reactivity to the repeat and N-terminal junction peptides (Figure 2C). In summary, high affinity to NANP and NVDP correlated with antibody cross-reactivity to the PfCSP N-terminal junction but not with C-CSP binding.

**Figure 2.**
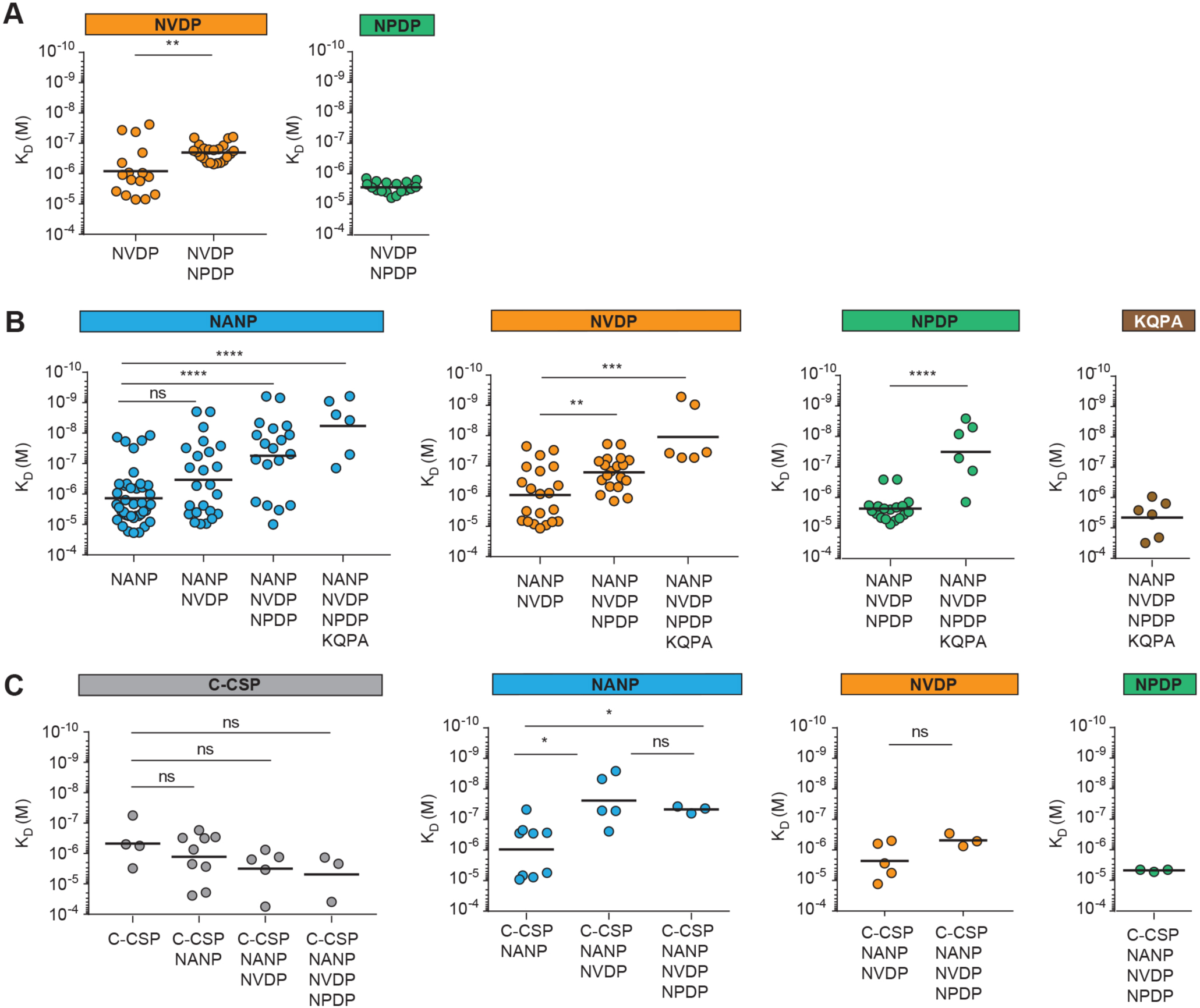
Cross-reactivity of NANP antibodies with the N-terminal junction but not C-CSP is associated with high affinity. (A-C) Affinity of epitope-specific compared to cross-reactive antibodies as determined by SPR. (A) NVDP affinity (orange; left) of NVDP-specific antibodies (n=16) and of NVDP, NPDP cross-reactive antibodies (n=22), and NPDP affinity of NVDP, NPDP cross-binders (green; right). (B) NANP (blue; left), NVDP (orange; center left), NPDP (green; center right) and KQPA (brown; right) affinities of NANP-specific antibodies (n=41) and of NANP binders with cross-reactivity to NVDP (n=24), NVDP and NPDP (n=19), or NVDP, NPDP and KQPA (n=6). (C) C-CSP (grey; left), NANP (blue; center left), NVDP (orange; center right) and NPDP (green; right) affinities of C-CSP-specific antibodies (n=4) and C-CSP binders with cross-reactivity to NANP (n=9), NANP and NVDP (n=5), or NANP, NVDP and NPDP (n=3). (A-C) Black horizontal lines indicate geometric means. P-values were calculated by Mann-Whitney test. *P < 0.05; **P < 0.01; ***P < 0.001; ****P < 0.0001; ns indicates statistically non-significant differences. See also Figure S2.

### Structural basis for C-CSP cross-reactivity of *IGHV3-33*-encoded NANP-reactive antibodies

Repertoire analyses showed that NANP-specific and, even more frequently, NANP-cross-reactive antibodies were encoded by *IGHV3-33* genes, particularly in association with *IGKV1-5* (Figures 3A and S3A). This observation agrees with the fact that high-affinity anti-NANP antibodies are frequently encoded by this gene combination (Murugan et al., 2018) and our observation that cross-reactivity with the N-terminal junction was associated with higher NANP affinity (Figure 2). Furthermore, the targeted cloning of surface immunoglobulin (Ig) negative plasmablast antibodies from the same donors identified several high-binding PfCSP antibodies exclusively encoded by *IGHV3-33*, *IGKV1-5* genes, demonstrating that this combination is strongly enriched among PfCSP-reactive B cells (Figure S3B; Table S2 (Murugan et al., 2018)). C-CSP cross-reactive antibodies were also encoded by *IGHV3-33* genes but in combination with different light chains, whereas NVDP-specific and NVDP, NPDP cross-reactive antibodies used a variety of *IGHV* gene segments other than *IGHV3-33* (Figures 3A and S3A).

**Figure 3.**
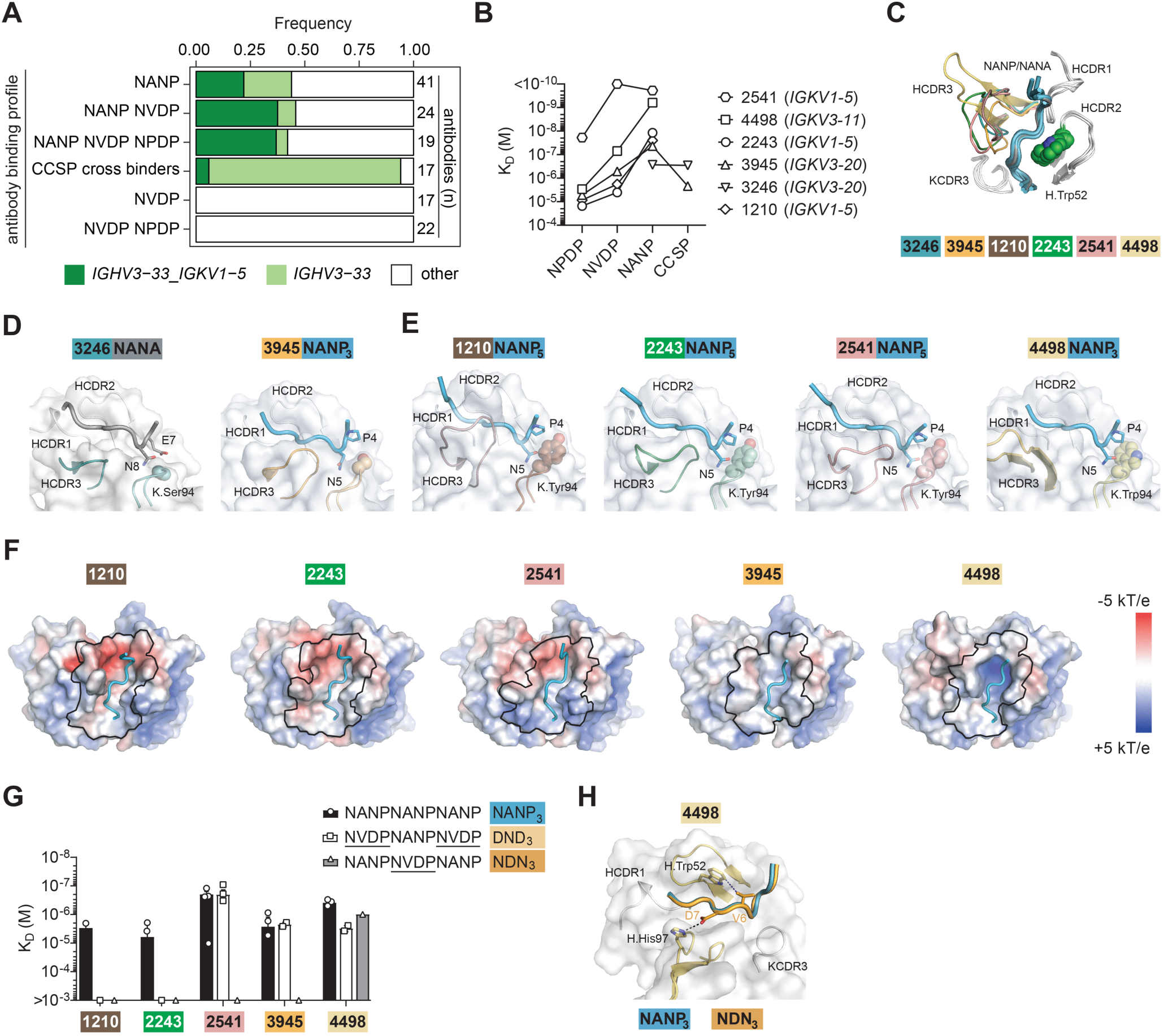
PfCSP peptide recognition by *IGHV3-33-*encoded PfCSP antibodies. (A) Frequency of antibodies with the indicated reactivity profiles encoded by *IGHV3-33* paired with either *IGKV1-5* (dark green) or other light chain V-gene segments (light green), or not encoded by *IGHV3-33* (other; white). n indicates the number of antibodies. (B) Affinity profiles of the indicated *IGHV3-33*-encoded monoclonal antibodies (mAbs) to NPDP, NVDP, NANP, and C-CSP as measured by SPR. The corresponding *IGKV* genes are annotated. (C) Superposition of the NANP_3_, NANP_5_ (blue) and NANA (grey) peptides in co-complexes with the indicated mAbs compared to mAb 1210 with NANP5 (Imkeller et al., 2018). Only the HCDR1, HCDR2, HCDR3, and KCDR3 are shown. HCDR3 regions are color-coded: 3246 (teal), 3945 (orange), 1210 (brown), 2243 (green), 2541 (pink), and 4498 (yellow). The germline-encoded H.Trp52 residues in HCDR2 are highlighted. (D) Binding of the C-CSP-reactive *IGHV3-33*-encoded mAbs 3246 and 3945 to NANA and NANP_3_ peptides, respectively. HCDR1 and HCDR2 (grey) are indicated. HCDR3 and KCDR3 of are highlighted in teal (mAb 3246) and orange (mAb 3945). The small serine residues at position 94 in KCDR3, which can accommodate Glu7 of the NANA-containing peptide and Pro4 in NANP_3_, are highlighted. (E) Binding of the non C-CSP-reactive *IGHV3-33*-encoded mAbs 1210 (Imkeller et al., 2018), 2243 and 2541 to NANP_5_, and 4498 to NANP_3_. HCDR1 and HCDR2 (grey) are indicated. HCDR3 and KCDR3 are highlighted in green (2243), brown (1210), pink (2541), and yellow (4498). The large tyrosine (mAbs 2243, 1210, 2541) and tryptophan (mAb 4498) residues at position 94 in KCDR3 are indicated. (F) Electrostatic surface potential of mAbs 1210 (Imkeller et al., 2018), 2243, 2541, 3945 and 4498 bound to NANP peptides. Electrostatic calculations were performed using APBS ((Baker et al., 2001); scale: −5 kT/e (red) to +5 kT/e (blue)). The paratope of each mAb is outlined in black. (G) Bar graphs indicate affinities of Fab fragments from the indicated antibodies for NANP_3_ and for the two NVDP-containing peptides DND_3_ (NVDPNANPNVDP) and NDN_3_ (NANPNVDPNANP) measured by isothermal titration calorimetry. Symbols indicate independent measurements. Means and SEM of three independent measurements are shown. (H) Binding of mAb 4498 to NDN_3_ (orange). HCDR1 and KCDR3 (grey) are indicated. HCDR1 and HCDR2 (yellow) are highlighted. Van der Waals interactions between HCDR2 Trp52 and Val6 of the NDN_3_ peptide are indicated by the deep blue dashed line, while the H-bond formed by HCDR3 His97 with Asp7 of the NDN_3_ peptide is shown by the black dashed line. The NANP_3_ peptide (blue) as bound by mAb 4498 (C-E) is shown for comparison. See also Figure S3 and Tables S3 and S4.

To better understand how cross-reactive antibodies recognize their epitopes, we selected five cross-reactive *IGHV3-33*-encoded NANP-binding antibodies (mAbs 2541, 4498, 2243, 3945 and 3246) with distinct binding profiles (Figures 3B and S3C) for structural studies. mAb 2541 bound NANP and NVDP with high affinity and showed cross-reactivity with NPDP. mAb 4498, and mAbs 2243 and 3945 with overall weaker affinity, preferentially bound NANP and, at lower levels, NVDP and NPDP. mAb 3945 additionally bound C-CSP. In contrast, mAb 3246 recognized exclusively NANP and C-CSP, but not NVDP or NPDP. Specifically, we determined the crystal structure of the Fab fragment of mAb 3246 in complex with a 14-aa peptide (PNRNVDENANANSA) derived from the C-CSP linker region comprising the unique NANA motif (Figure 1A) to a resolution of 2.40 Å (Table S3). We also determined the crystal structures of Fab fragments of mAbs 3945 and 4498 in complex with a NANP trimer peptide (NANP_3_), and of mAbs 2243 and 2541 in complex with a NANP pentamer peptide (NANP_5_) to a resolution of 1.40 Å, 2.90 Å, 2.00 Å, and 2.55 Å, respectively (Table S3). For comparison, we used the reported structure of NANP_5_ in complex with the *IGHV3-33-, IGKV1-5*-encoded mAb 1210 with a similar binding profile to mAb 2243 (Figure 3B; (Imkeller et al., 2018)). The six antibodies bound the respective PfCSP peptides in nearly identical inverted S-shape folds, and had highly similar HCDR1, HCDR2, and KCDR3 antibody conformations (Figure 3C). Importantly, the *IGHV3-33* germline-encoded H.Trp52 in the HCDR2, a key residue for binding to NANP as previously reported for 1210 and other antibodies (Imkeller et al., 2018; Oyen et al., 2017; Tan et al., 2018), was critically involved in the binding of mAbs 3945, 2243, 2541, and 4498 to the NANP peptides, and of mAb 3246 to its core epitope (DENANANS) in the NANA peptide (Figures 3C and S3D; Table S4). Notably, mAbs 2541 and 4498 seemed optimally disposed to bind NANP. Aromatic residues in their HCDR3s packed against Pro of the repeat and, with H.Trp52, formed an aromatic cage that likely contributes to their high NANP affinity (Figure S3D). Similarly, a vicinal disulphide bond in mAb 2243 between HCDR3 residues H.Cys100D and H.Cys100E mimicked this stacking effect, contacting Pro of the repeat (Figure S3D). In total, 15% (10/66) of *IGHV3-33* antibodies contained two neighbouring cysteines in their HCDR3 compared to 0.01% (2/134) in all other PfCSP-reactive antibodies, likely contributing to the strong association between *IGHV3-33* gene usage and NANP-reactivity.

The small K.Ser94 side chain at the tip of KCDR3 in mAb 3246 accommodated binding to the NANA peptide (Figure 3D). mAb 3945 also had a *IGKV3-20*-encoded K.Ser94 and was cross-reactive with the NANA peptide (Figures 3B and 3D). In contrast, the four non-C-CSP cross-reactive antibodies, mAbs 1210, 2243, 2541, and 4498 harboured large *IGKV1-5*-encoded Tyr or *IGKV3-11*-encoded Trp side chains at the same position (Figure 3E). We propose that these bulky KCDR3 residues explain the lack of C-CSP cross-reactivity for these antibodies due to steric constraints with a glutamic acid in the NANA peptide instead of a proline in the NANP peptide N-terminal of the core motif (Figures 3D and 3E).

Importantly, the *IGHV3-33*-encoded mAbs generated a spectrum of electrostatic potential in their paratopes (Figure 3F). Differences in the electrostatic potential of the antibodies correlated with their propensity to bind NVDP-containing epitopes (Figure 3G). The more electronegative paratopes of mAbs 1210 and 2243 bound NANP_3_ but disfavoured binding to 3-mer peptides containing NVDP motifs in isothermal titration calorimetry (ITC) experiments, likely due to the electrostatic repulsion (Figure 3G). In contrast, the less electronegative paratope of mAb 2541 and the polar paratopes of mAbs 3945 and 4498 were more amenable to binding NVDP-containing peptides. mAbs 2541 and 3945 bound the 3-mer peptides only when the NVDP motifs were flanking the NANP motif (DND_3_, NVDPNANPNVDP) but not when a single NVDP motif was surrounded by two NANP motifs (NDN_3_, NANPNVDPNANP); whereas mAb 4498 interacted with both peptides, although with lower affinity compared to NANP_3_. A 1.70 Å co-complex structure of mAb 4498 with NDN_3_ revealed that the peptide adopted the same conformation as NANP_3_ and occupied the same position in the mAb 4498 paratope, illustrating how a single antibody can accommodate different epitopes (Figure 3H). Specifically, H.Trp52 in HCDR2 and H.His97 in HCDR3 engaged in direct contacts with Val and Asp in the central NVDP motif, respectively. Thus, despite the highly similar binding mode of *IGHV3-33*-encoded antibodies, fine differences in KCDR3, HCDR3 and paratope electrostatic potential explain the specificity in their cross-reactivity profiles and binding affinities for the different epitopes.

### NANP motifs are the core epitope of PfCSP antibodies with cross-reactivity to the N-terminal junction

We also identified PfCSP cross-reactive antibodies that were not encoded by *IGHV3-33* genes (Table S1). To understand how a non-*IGHV3-33*-encoded antibody binds to the NANP repeat and the N-terminal junction peptides, we investigated mAb 4493, an *IGHV3-49-, IGKV3-20*-encoded antibody with high affinity to NANP and the junctional peptides (Figures 4A and S4A). Specifically, we determined the crystal structures of the mAb 4493 Fab fragment in complex with KQPA (1.93 Å), NPDP (2.4 Å), the two shortened NVDP peptides, NDN_3_ (2.1 Å) and DND_3_ (2.02 Å), and with NANP_3_ (2.15 Å) (Figures 4B-4F; Table S5). mAb 4493 contacted all peptides via its HCDR3 and KCDR3, while its KCDR2 made additional contacts with NPDP, DND_3_ and NANP_3_ peptides. In contrast to the conformation adopted in complex with *IGHV3-33*-encoded antibodies, the 4493-bound peptides adopted a U-shaped conformation centered around the *IGHV3-49* germline-encoded amino acid H.Arg52, which alone formed up to five H-bonds with the peptides. The preferential binding of mAb 4493 to NPDP (Figure 4A) was associated with more extensive H-bond interactions compared to the other peptides (11 H-bonds with NPDP compared to 8 or 9 with DND_3_ and NANP_3_, respectively; Figures 4A-4F; Table S6).

**Figure 4:**
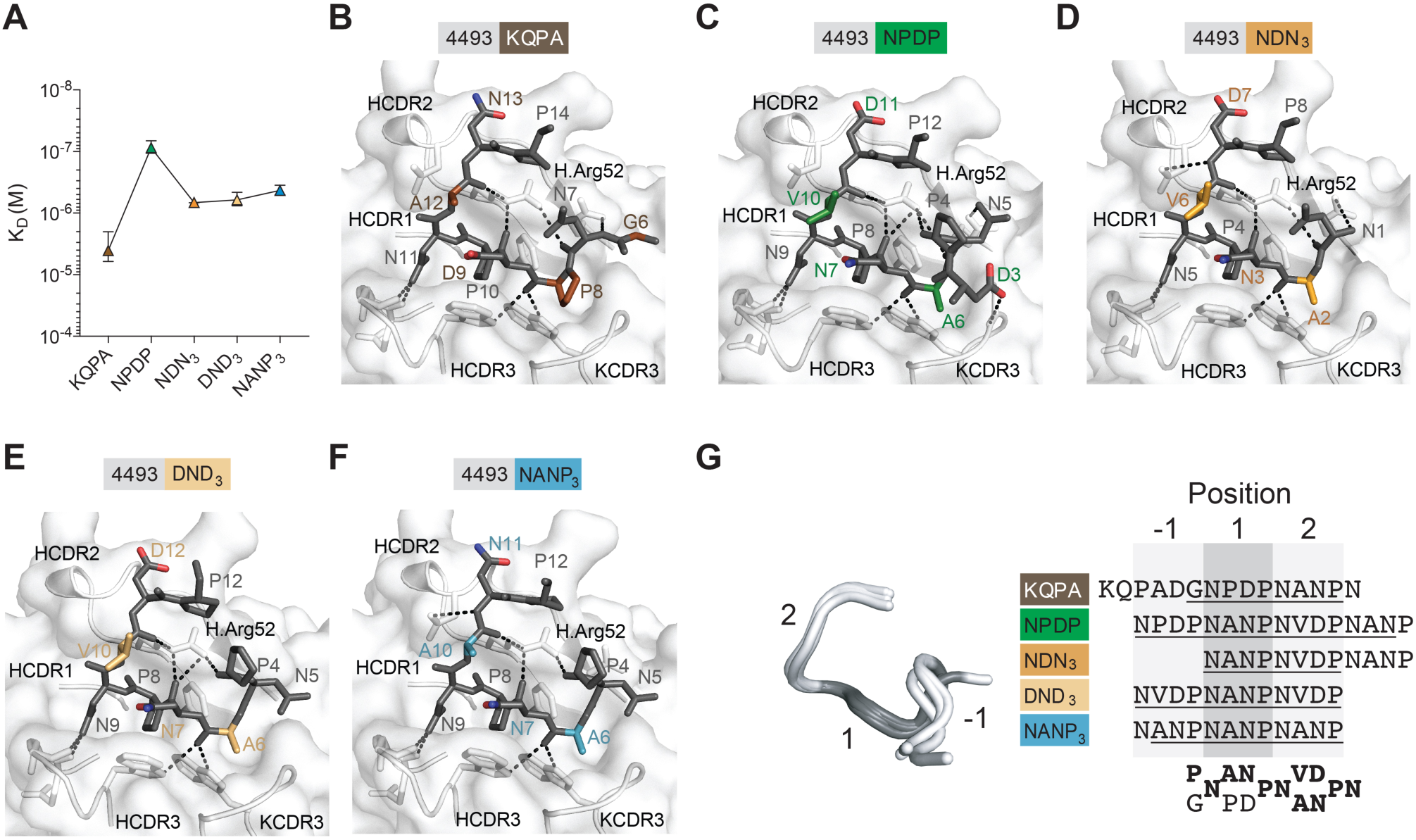
Preferential recognition of different PfCSP peptides around NANP core epitopes by the non-*IGHV3-33*-encoded cross-reactive mAb 4493. (A) Affinity of mAb 4493 to the indicated peptides as measured by isothermal titration calorimetry (ITC). (B-F) mAb 4493 in co-complex with the peptides KQPA (B), NPDP (C), NDN_3_ (D), DND_3_ (E) and NANP (F). Peptides are shown in dark grey, except for atomic positions that vary between peptides, which are colored to highlight how the paratope accommodates these subtle differences. H-bonds between mAb 4493 and the respective peptides are shown as black dashed lines. HCDR1-3 and KCDR3 are indicated. HArg52 in HCDR2 is labeled. (G) Overlay of the peptides as observed in the antibody co-complex structures (B-F). Three distinct paratope positions (−1, 1 and 2) are indicated. Positioning of the 4-aa motifs in the indicated peptides at position −1, 1, and 2 of the mAb 4493 paratope as observed in the co-complex structures is shown (B-F). Amino acid residues resolved in the X-ray crystal structures are underlined. The preferred amino acid residues at positions −1, 1, and 2 are indicated. See also Figure S4 and Tables S5 and S6.

Strikingly, mAb 4493 bound all peptides in largely superimposable conformations (rmsd < 0.2 Å) (Figure 4G). The mAb 4493 paratope engaged with three consecutive 4-aa motifs at three distinct positions, here referred to as position −1, 1, and 2. In five of the six structures, position 1 was occupied by NANP, illustrating a strong preference of the core paratope for this motif (Figure 4G and Table 1). Indeed, although the NPDP motif could be accommodated at position 1 as observed in the co-complex of mAb 4493 with the KQPA peptide, this binding mode, which placed the NANP motif in position 2, was associated with overall weaker affinity (K_D_ = 4 μM) compared to all other peptides (K_D_ < 676 nM) including the high-affinity NPDP peptide interaction with NPDP, NANP, and NVDP motifs at positions −1, 1, and 2, respectively (K_D_ = 87 nM). Interaction with NPDP at position 1 and NANP at position 2 has previously been observed for mAb MGG4, an *IGHV3-33*-like antibody, in a co-complex with an NPDPNANP peptide, although the structure of this antibody with other peptides has not been reported ((Tan et al., 2018) Table 1). The NVDP motif was not favorably accommodated by 4493 at position 1 due to the Val side chain being too bulky for the paratope, whereas Ala/Pro residues of NANP and NPDP motifs are optimally buried by the KCDR3. In contrast to the strong preference for the NANP motif at position 1, mAb 4493 could accommodate NPDP, NANP, and NVDP equally well at position −1, and NANP or NVDP at position 2. Likely due to the strong preference for binding to NANP at position 1, position 2 was more frequently occupied by NVDP, thereby placing NPDP into position −1 according to the natural order of the NPDP-, NANP-, NVDP-motifs in full-length PfCSP (Figure 1A). Our structures indicate that electrostatic potential differences between Asn and Asp in the PfCSP repeat do not impact the cross-reactivity of mAb 4493. Strikingly, the antigen recognition mode of mAb 4493 was highly similar to CIS43 (Figure S4B), a potent cross-reactive *IGHV1-3*-, *IGKV4-1*-encoded PfCSP antibody with preference for binding to the N-terminal junction that had been induced by immunization with irradiated sporozoites (Kisalu et al., 2018). Despite differences in the binding orientation between the two antibodies, CIS43 also recognizes junctional peptides in a U-shaped conformation through interactions of the core paratope centered on the NANP motif (Table 1 and Figure S4C). The same preference for NANP at position 1 that was observed for mAbs 4493 and CIS43 was also seen in X-ray and cryo-EM co-complexes of two other human anti-PfCSP antibodies, mAb CIS42 (*IGHV7-4-1*, *IGLV2-23*-encoded; induced by irradiated sporozoite immunization; (Kisalu et al., 2018)) and mAb 311 (*IGHV3-33*-, *IGLV1-40*-encoded; induced by RTS,S vaccination (Oyen et al., 2017)), respectively, with different binding modes, epitope-reactivity profiles, and Ig gene usage (Table 1, Figures S4D and S4E). The core position 1 in the paratopes of these non-*IGHV3-33-* and *IGHV3-33-*encoded antibodies was optimally suited to accommodate NANP, whereas variability was observed at position −1 and to lesser extent position 2. Together, the data define that binding of cross-reactive antibodies to PfCSP is overall preferentially centered on interactions of the paratope with NANP flanked by N terminal NP/DP and C terminal NA/NV residues in a conserved NP/DP-NANP-NA/NV motif independently of their gene usage or binding profile (Table 1).

**Table 1.**
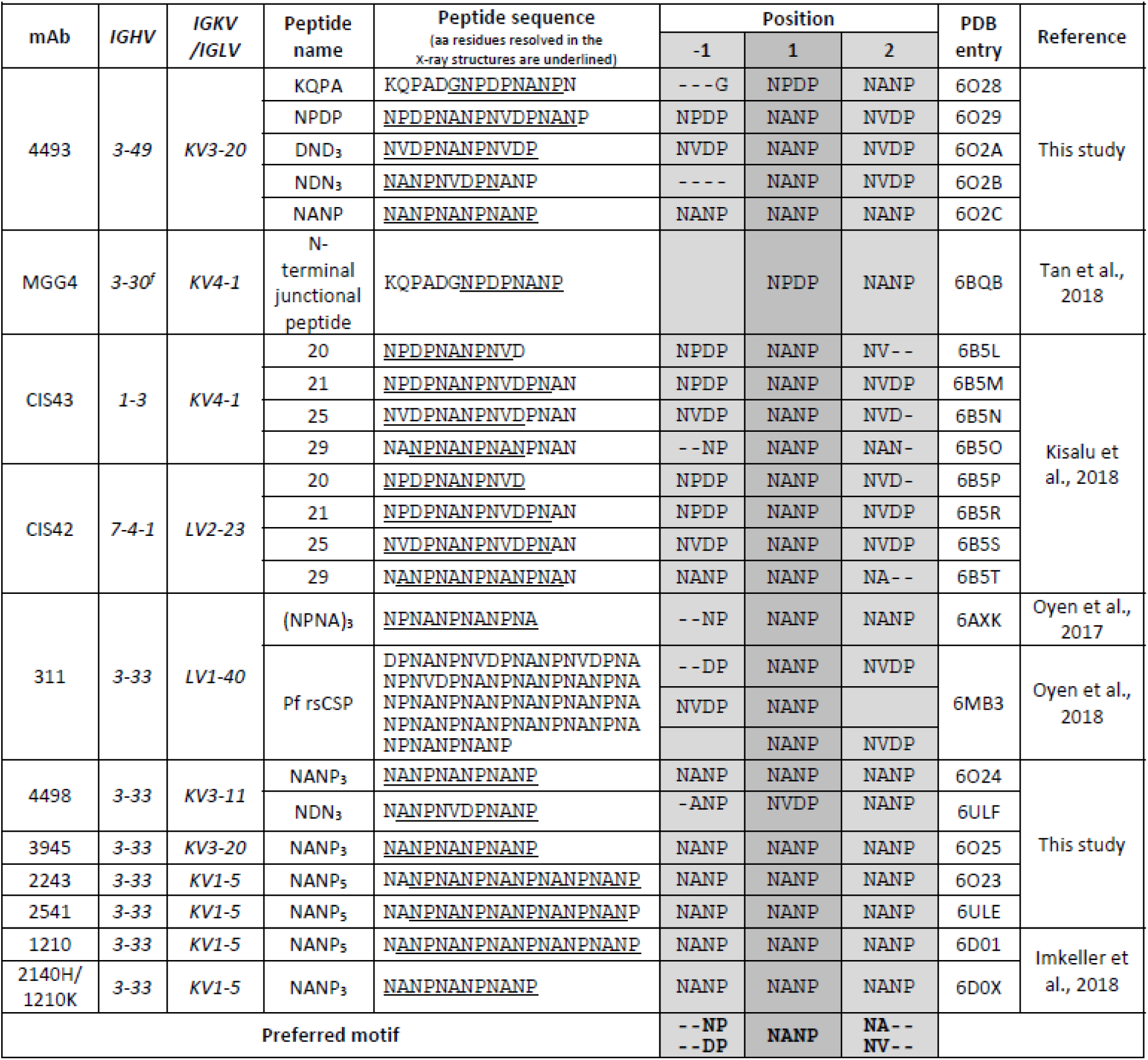
Conserved binding mode of anti-PfCSP antibodies in co-complexes with N-terminal junction and NANP peptides.

### Evolution of anti-PfCSP antibody binding profiles

To determine the role of NANP binding in the development of cross-reactive antibodies, we assessed the evolution of the response over time. The frequency of NANP-binding memory B cell antibodies with strong cross-reactivity to the junctional peptides, especially NPDP and NVDP, increased with repeated parasite exposure and was higher after the third immunization and challenge compared to the second immunization (Figure 5A). The vast majority was also class-switched to IgG (Figure 5B), including many *IGHV3-33*-, *IGKV1-5*-encoded but also rare antibodies with non-prominent gene combinations such as mAb 4493 (Figure 5C; Table S1). Most cross-reactive antibodies belonged to clonally expanded and diversified B cell clusters (Figures 5D and 5E; Table S1), but the majority showed no differences in their cross-reactivity profile independently of their binding preference and their absolute affinity (Figure 5D). In rare cases, individual members of a cluster gained or lost reactivity to one or all peptides (Figure 5E). However, this was independent of their absolute mutation load. Affinity to the junctional peptides was not always gained, and in some instances it was also lost with accumulating mutations, as previously demonstrated for NANP binding (Murugan et al., 2018).

**Figure 5:**
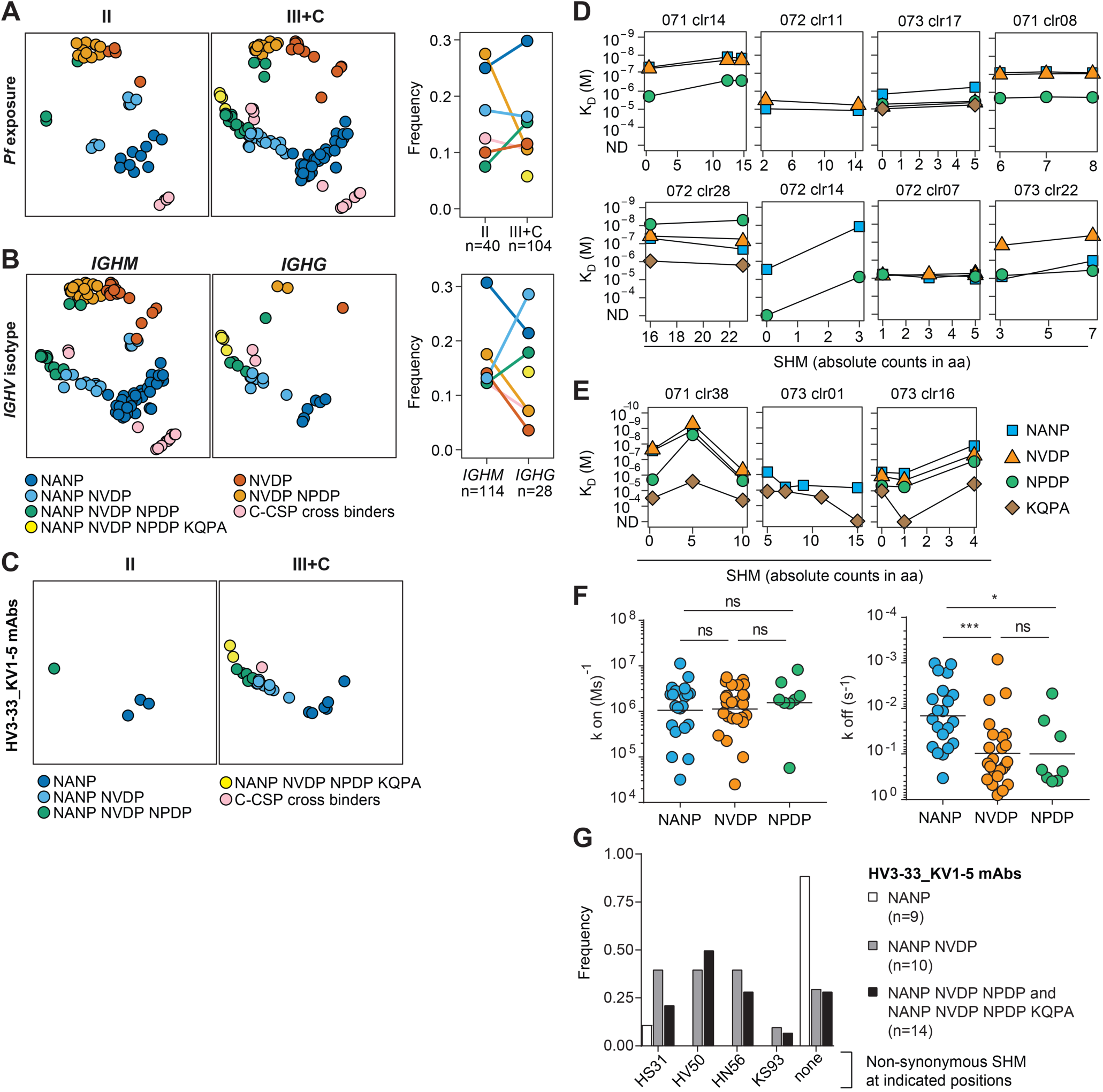
Maturation of anti-PfCSP cross-reactive antibody profiles over successive Pf exposures. (A and B) t-SNE clustering-based illustration of PfCSP-reactive antibodies with the indicated color-coded binding profiles (left and center). The respective frequencies are shown and the numbers of antibodies (n) are indicated (right). (A) After the second (II; left) or third (III) Pf exposure and challenge (C) infection (III+C; center). (B) Expressing *IGHM* (left) or *IGHG* (center) isotypes. (C) t-SNE clustering-based illustration of *IGHV3-33, IGKV1-5*-encoded PfCSP-reactive antibodies with the indicated color-coded binding profiles after the second (II; left) or third (III) Pf exposure and challenge (C) infection (III+C; right) as shown in (A). (D and E) Affinity of clonally related antibodies with different numbers of *IGH* and *IGL* somatic mutations (aa exchanges) to the indicated peptides determined by SPR. (D) Clusters of clonally related antibodies with similar binding profiles to the indicated peptides for all cluster members independently of their somatic mutation load. (E) Clusters of clonally related antibodies with mutated members that differ in their binding profiles defined by >10-fold differences in antibody affinity to any of the indicated peptides. (F) Kinetic on (k-on; left) and kinetic off (k-off; right) rates of NANP, NVDP, NPDP cross-reactive antibodies for binding to the indicated peptides. Black bars indicate geometric means. P-values were calculated by Mann-Whitney test. *P < 0.05; ***P < 0.001; ns indicates non-statistically significant differences. (G) Frequency of NANP-specific (n=9) *IGHV3-33*, *IGKV1-5*-encoded memory B cell and plasmablast antibodies (Murugan et al., 2018) with or without (none) the indicated aa somatic mutations (SHM) previously shown to directly (IGHV_S31, IGHV_V50I) or indirectly (IGHV_N56, IGKV_S93) increase NANP affinity (Imkeller et al., 2018) compared to NANP binders with cross-reactivity to NVDP (n=10) or NVDP and NPDP (n=14).

To assess what drove the selection of high-affinity cross-reactive antibodies, we compared the kinetic rates of cross-reactive antibodies to associate with (k_on_) or dissociate from (k_off_) NANP, NVDP, and NPDP peptides (Figure 5F). Despite similar k_on_ rates for all peptides, the antibodies differed in their k_off_, which was significantly lower for binding to the NANP repeat compared to NVDP or NPDP, suggesting that the antibodies had been primarily selected for their ability to bind the NANP repeat and not the junctional peptides. Although overall infrequent, many of the *IGHV3-33-* and *IGKV1-5*-encoded cross-reactive but only few NANP-specific antibodies carried select somatic mutations that have been shown to either directly (H.S31, H.V50) or indirectly (H.N56, K.S93) improve NANP binding through homotypic antibody-antibody interactions (Figure 5G; (Imkeller et al., 2018)). Thus, in line with our structural analyses, the accumulation of cross-reactive antibodies in response to repeated parasite exposure was likely due to the direct association of antibody cross-reactivity with NANP affinity in the core epitope and continuous selection of high-affinity clones from the naïve repertoire, or of B cells that had gained affinity through somatic mutations, especially of *IGHV3-33-*, *IGKV1-5*-encoded antibodies.

### Antibody binding to NANP but not the N-terminal junction or C-CSP correlates with potent Pf inhibition *in vitro*

To determine whether the observed differences in antibody-binding preferences affected their Pf-inhibitory capacity, we compared the activity of 139 antibodies representing all cross-reactivity profiles to inhibit the sporozoite traversal of hepatocytes *in vitro* (100 µg/ml; Figure 6; Table S1; (Murugan et al., 2018)). NVDP-specific antibodies were overall poor inhibitors (mean 55%; Figure 6A). Although the mean inhibitory activity was significantly higher for NVDP binders with cross-reactivity to NPDP (mean 74%), in line with their higher affinity (Figure 2A), none of the antibodies reached 100% inhibition. NANP-specific antibodies were overall better Pf inhibitors than NVDP-specific antibodies (mean 69%) with two antibodies that conferred complete inhibition (Figure 6B). The mean potency of NANP-binders increased significantly with cross-reactivity to the N-terminal junction and was higher for antibodies that bound to two or three junctional peptides (mean 92%) than for anti-NANP antibodies with limited cross-reactivity to NVDP only (mean 76%). C-CSP specific antibodies were the weakest inhibitors of all (mean 13%) but their potency was significantly improved with associated cross-reactivity and higher affinity to NANP as previously reported (mean 72%; (Scally et al., 2018)), and with additional binding to NVDP and NPDP (mean 95%; Figures 6C and 2C). Thus, NANP binding was associated with parasite inhibition, explaining the overall low protective activity of non-NANP-reactive antibodies.

**Figure 6:**
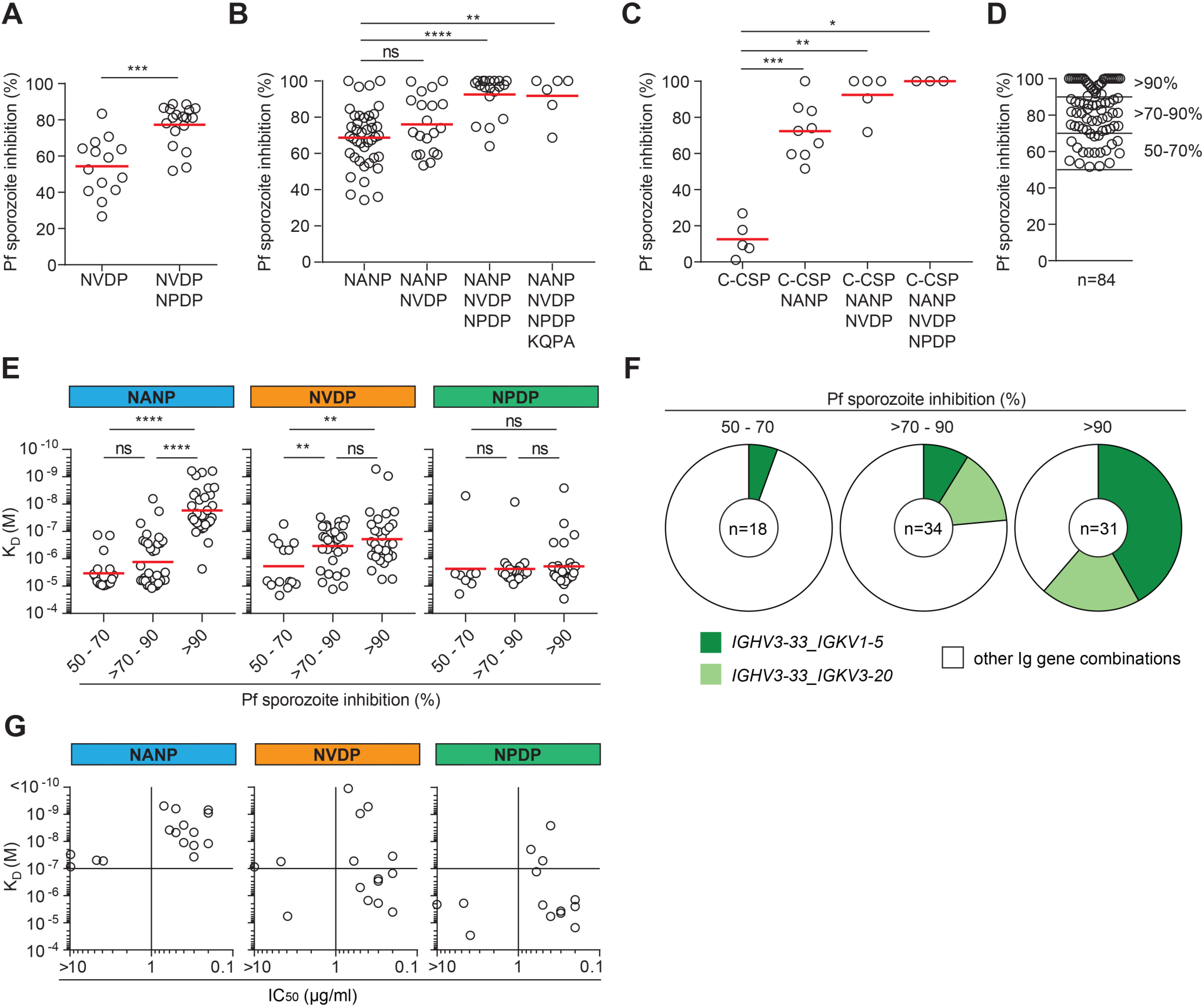
High-affinity cross-reactive antibodies are potent parasite inhibitors. (A-C) Capacity of antibodies (n=139, 100 µg/ml) with the indicated binding profiles to inhibit the hepatocyte traversal activity of Pf sporozoites *in vitro*. (A) NVDP-specific antibodies (n=14) and NVDP, NPDP cross-reactive antibodies (n=18). (B) NANP-specific antibodies (n=41) and NANP binders with cross-reactivity to NVDP (n=20), NVDP and NPDP (n=18), or NVDP, NPDP and KQPA (n=6). (C) C-CSP-specific antibodies (n=5) and C-CSP binders with cross-reactivity to NANP (n=9), NANP and NVDP (n=5), or NANP, NVDP and NPDP (n=3). (D) Cross-reactive PfCSP antibodies with low (50-70%; n=18), intermediate (>70-90%; n=33), or high (>90%; n=32) Pf sporozoite traversal inhibitory activity. (E) NANP (blue, left), NVDP (orange, middle), and NPDP (green, right) affinities of cross-reactive antibodies with low (50-70%), intermediate (>70-90%), or high (>90%) *in vitro* Pf hepatocyte traversal-inhibitory activity. (F) Frequency of cross-reactive antibodies encoded by *IGHV3-33* paired with either *IGKV1-5* (dark green) or *IGKV3-20* (light green) with low (50-70%), intermediate (>70-90%) or high (>90%) *in vitro* Pf hepatocyte traversal activity. n in the pie chart centers indicate the number of antibodies. (G) IC_50_ values versus NANP (left), NVDP (middle), and NPDP (right) of selected antibodies with >95% Pf hepatocyte traversal-inhibitory activity (D). (A-C, E) Red bars indicate arithmetic and geometric mean in A-C and E, respectively. (A-D) Data is representative of at least two independent experiments. P-values were calculated by Mann-Whitney test. *P < 0.05; **P < 0.01; ***P < 0.001; ****P < 0.0001; ns indicates non-significant statistical differences. (G) IC_50_ values were calculated from data of at least three independent experiments. See also Table S7.

We next examined whether the affinities (K_D_) of cross-reactive antibodies to NANP, NVDP, or NPDP were predictive of low (50-70%), intermediate (>70-90%) or high (>90%) levels of Pf inhibition (Figure 6D and E). NANP, but not NVDP or NPDP affinity was strongly associated with anti-Pf activity. The most potent inhibitors (>90%) were almost exclusively antibodies with NANP affinity below 10^-7^ M (Figure 6E). The NVDP affinity of the most potent inhibitors was overall weaker (mean K_D_= 1.9 x 10^-7^ M) than their NANP affinity (mean K_D_= 1.7 x 10^-8^ M), and comparable to antibodies with intermediate Pf-inhibitory activity (mean K_D_= 3.4 x 10^-7^ M). Although rare antibodies with high NPDP affinity were identified in all three groups, NPDP affinity was overall low in the range of K_D_ 10^-6^ M and did not discriminate between antibodies with low, intermediate, or high levels of Pf inhibition. In summary, cross-reactive antibodies with low anti-parasite activity were mostly weak binders, whereas high affinity to NANP but not to NVDP or NPDP discriminated the most potent cross-reactive inhibitors from antibodies with intermediate anti-parasite activity in the sporozoite traversal assay.

Next, we selected 16 of the most potent non-clonally-related antibodies with >95% inhibition activity, including one NANP-specific and 15 cross-reactive antibodies with different binding profiles, and determined their IC_50_ values (Table S7). Eleven antibodies were encoded by *IGHV3-33* in combination with *IGKV1-5* (9/11) or *IGKV3-20* (2/11), reflecting the strong enrichment of this gene combination among antibodies with >90% Pf-inhibitory activity (Figure 6F), whereas the other five antibodies were encoded by different *IGHV3* and various *IGKV* and *IGLV* light chain genes (Table S7). Regardless of their binding profile, the IC_50_ values of all 16 antibodies ranged from 0.2 µg/ml to >20 µg/ml and correlated with their NANP (K_D_ 10^-7^ – 10^-9^ M) but not their NVDP (K_D_ 10^-6^ – 10^-9^ M) or NPDP (K_D_ 10^-5^ – 10^-9^ M) affinities (Figure 6G; Table S7). Strikingly, with one exception, the IC_50_ values of all eleven *IGHV3-33-*encoded antibodies were <1.0 µg/ml (mean = 0.7 µg/ml). In contrast, only 2/5 antibodies with other gene combinations showed IC_50_ values <1.0 µg/ml, whereas for the other three, the values ranged from 4.7 µg/ml to >20.0 µg/ml. Thus, non-*IGHV3-33*-encoded antibodies with high Pf-inhibitory activity were rare. The most potent antibodies in this assay all showed NANP affinities <10^-7^ M independently of their binding profiles, and included cross-reactive and rare epitope-specific antibodies.

### Antibody-mediated protection from infection *in vivo*

To determine the inhibitory activity of the most potent antibodies *in vivo*, we measured the protective efficacy of selected antibodies with IC_50_ values <1.0 µg/ml as time to development of blood-stage parasites (prepatency) in mice infected with *Pf*CSP-expressing transgenic *Plasmodium berghei* parasites (*Pb*-*Pf*CSP) by mosquito bites 20 h after intraperitoneal antibody injection (Figure 7A). First, we assessed the potency of three cross-reactive *IGHV3-33*-encoded antibodies with or without C-CSP cross-reactivity (mAbs 1210, 2164, and 4476; Table S8) compared to a non-protective C-CSP-specific negative control (mAb 1710, (Scally et al., 2018); Figure 7B and 7C). All three antibodies showed a preference for NANP and weaker binding to NVDP and NPDP but not to KQPA. mAb 2164 showed overall higher affinities compared to mAbs 4476 and 1210, but only mAb 4476 recognized C-CSP (Figure 7B). After passive transfer of 300 µg antibody per mouse, the highest protection (56% parasitemia-free mice) was observed for mAb 1210, but the differences compared to mAb 2164 (33%) and mAb 4476 (30%) were not statistically significant (Figures 7C; Table S8). Thus, cross-reactivity with C-CSP and overall affinity did not predict the *in vivo* potency of these *IGHV3-33*-encoded antibodies with preference for NANP and limited binding to the N-terminal junction.

**Figure 7.**
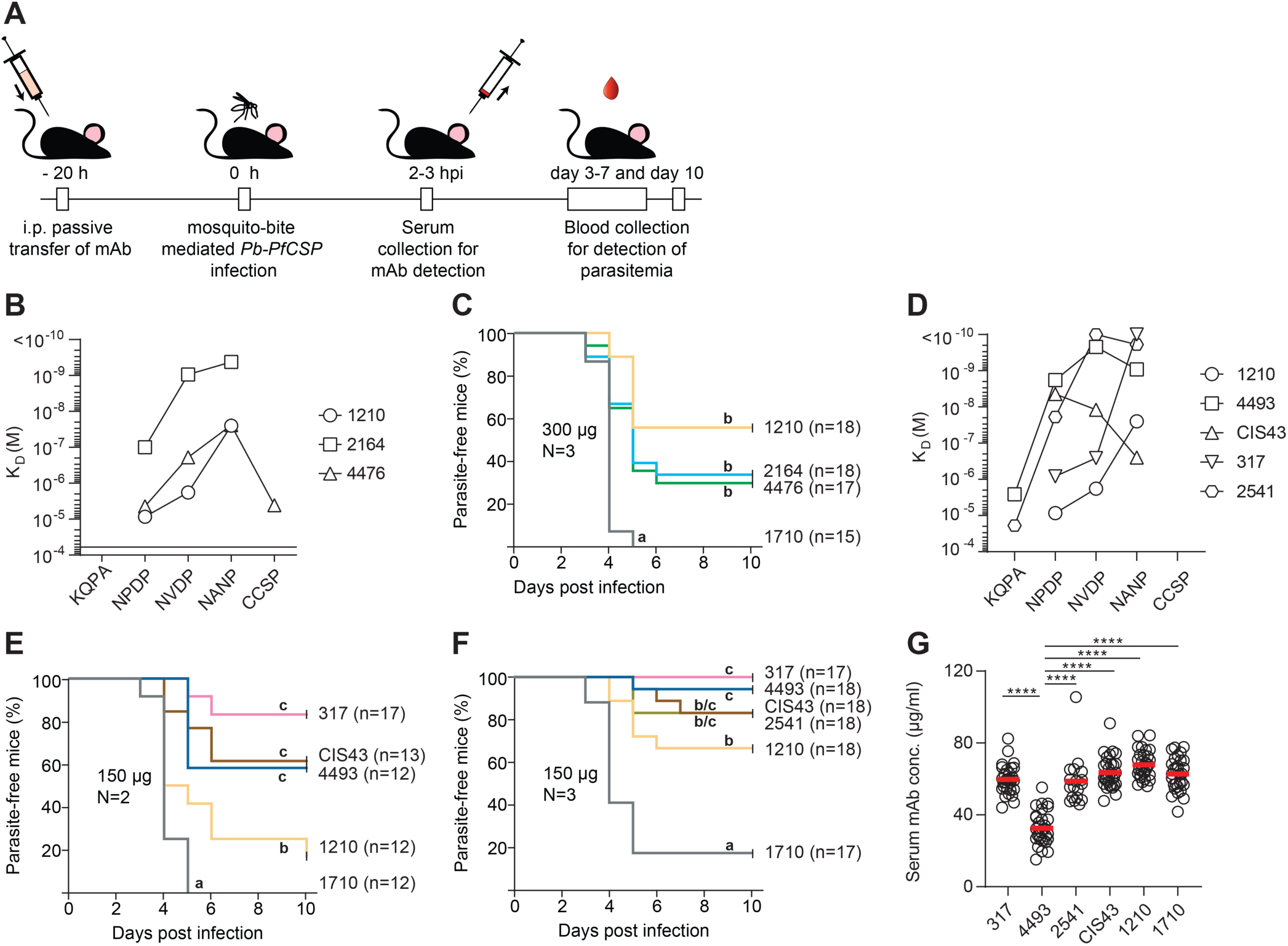
*In vivo* protective activity of cross-reactive *IGHV3-33*- and non-*IGHV3-33*-encoded PfCSP antibodies with different binding profiles. (A) Scheme and time line of the experiments. Antibody potency was assessed in C57BL/6 mice after intra peritoneal (i.p.) passive transfer of mAb 20 h before exposure to bites of mosquitoes infected with *Plasmodium berghei* transgenic parasites expressing *P. falciparum* CSP (*PbPfCSP*; (Triller et al., 2017)). Two to three h post infection (hpi) blood was collected and serum concentrations of the monoclonal antibodies were determined by ELISA. The time to blood-stage parasitemia was monitored by blood smear between day 3 and day 10 after the mosquito-bite exposure. (B, C) *In vivo* protective activity of cross-reactive *IGHV3-33*-encoded PfCSP antibodies with different binding profiles. (B) Affinity profiles of mAbs 1210 (Imkeller et al., 2018; Murugan et al., 2018), 2164 and 4476 to KQPA, NPDP, NVDP, NANP and C-CSP as determined by SPR. (C) Capacity of the indicated antibodies (300 µg/ml) to protect mice from infection by mosquito bites with *PbPfCSP* parasites compared to the non-protective C-CSP-specific antibody 1710 (Scally et al., 2018). Data shows the percentage of parasite-free mice in three independent experiments. The total number of mice per antibody is indicated (n). (D-G) *In vivo* protective activity of cross-reactive non-*IGHV3-33*-encoded PfCSP antibodies with different binding profiles. (D) Affinity profiles of mAbs 4493, CIS43 (Kisalu et al., 2018) and 317 (Oyen et al., 2017) to KQPA, NPDP, NVDP, NANP and C-CSP as determined by SPR in comparison to the *IGHV3-33*-encoded mAbs 1210 (Imkeller et al., 2018) and 2541. (E, F) Capacity of the indicated antibodies (150 µg/ml) to protect mice from infection by mosquito bites with *PbPfCSP* parasites compared to the non-protective C-CSP-specific antibody 1710 (Scally et al., 2018). Data show the percentage of parasite-free mice in two (E) or three (F) independent experiments. The total number of mice per antibody is indicated (n). (G) Serum concentrations in individual mice (open circles) after passive transfer of the indicated monoclonal antibodies (E and F; n=34 for mAb 317, n=31 for mAb 4493, n=18 for mAb 2541, n=31 for mAB CIS 43, n=30 for mAb 1210, n=29 for mAb 1710) at the time of parasite challenge. Horizontal red lines indicate means. P-values were calculated by Mann-Whitney test. ***P < 0.001. (B, D) Data represent the mean from three independent measurements. Horizontal lines indicate threshold K_D_ values of 5 x 10^-5^ M. (G) Data shows the mean of at least two independent measurements. See also Figure S5 and Tables S8-S11.

Next, we compared mAb 4493 (*IGHV3-49-, IGKV3-20*) with high affinity to NANP, NVDP and NPDP, and KQPA cross-reactivity to mAb 1210 in the same model (Figures 7D-7G; Figure S5). Additionally, we included two potent published antibodies. mAb CIS43 (*IGHV1-3*-, *IGKV4-1*-encoded; (Kisalu et al., 2018)) was similar to mAb 4493 in its strong affinity for NPDP, but showed a clear preference for this peptide with no measurable KQPA cross-reactivity, and about 10-fold and 100-fold lower NVDP and NANP affinity, respectively (Figure 7D). mAb 317 (*IGHV3-30-3*-, *IGKV1-5*-encoded; (Oyen et al., 2017)) had an exceptionally high NANP affinity (< 10^-10^ M) but low cross-reactivity to NVDP and NPDP. At a dose of 300 µg per mouse, mAbs 4493, CIS43, and 317 showed similar levels of protection from blood-stage parasitemia at 83%, 91%, and 100%, respectively (Figure S5; Table S9). To better resolve potential differences in the inhibitory activities of the antibodies, we halved the dose to 150 µg per mouse (Figures 7E and 7F). Although the degree of protection was overall lower, all three antibodies retained their high potency compared to mAb 1210. mAb 317 protected 83% of mice compared to 62% for mAb CIS43 and 58% for mAb 4493, but these differences were not statistically significant. Of note, independent of the dose, mAb 4493 consistently showed, on average, two-fold lower serum concentrations at the time of challenge than the other antibodies (Figures 7G and S5B; Table S10). To determine whether *IGHV3-33*-encoded antibodies could reach similar levels of potency, we compared the antibodies to mAb 2541 with higher affinity to NANP and NVDP than any *IGHV3-33*-, *IGKV1-5*-encoded antibody in our panel, as well as additional cross-reactivity to NPDP and KQPA (Figure 7F and Table S7). In direct comparison to mAbs CIS43, 4493, and 317, mAb 2541 was as protective as these non-*IGHV3-33*-encoded antibodies (Figures 7F and 7G; Table S11). Thus, the most potent PfCSP antibodies with high levels of *in vivo* protection against malaria parasites showed exceptional affinity to the repeat or the junctional epitopes and were encoded by *IGHV3-*33 and *IGHV3-30-3, IGKV1-5* (mAbs 2541 and 317, respectively; (Murugan et al., 2018; Oyen et al., 2017)) or other gene combinations (mAbs 4493 and CIS43; (Kisalu et al., 2018; Murugan et al., 2018)).

## Discussion

Our results uncover a molecular pattern underlying the binding of human antibodies to PfCSP and provide a link between epitope cross-reactivity, affinity, and parasite inhibitory capacity that is of great relevance for the design of a next-generation PfCSP vaccine. We found that sequence similarities between the 4-aa motifs in the N-terminal junction, central domain, and C terminus of PfCSP, and the repetitive organization of the NANP and NVDP motifs contribute to binding promiscuity and explain the high frequency of cross-reactive compared to epitope-specific PfCSP antibodies. Cross-reactivity was a feature of antibodies encoded by different Ig-gene combinations, including antibodies that developed in response to natural parasite exposure (Triller et al., 2017), but also of the well-characterized murine repeat binder mAb 2A10 with strong reactivity to the junctional peptides (Zavala et al., 1983), demonstrating that cross-reactivity was not limited to human antibodies (Figure S6). Although we identified a large variety of different PfCSP binding profiles, the vast majority of antibodies recognized NANP, either exclusively or with additional cross-reactivity to other motifs. The higher number of NANP compared to NVDP, NPDP, and NANA motifs per PfCSP molecule likely contributed to this bias in the response to live sporozoites, in addition to differences in immunogenicity and accessibility of PfCSP epitopes, as previously reported for the C terminus (Scally et al., 2018). Indeed, cross-reactive and epitope-specific antibodies that recognized the N-terminal junction with many 4-aa motifs were more frequent than those with reactivity to the C terminus, which only contains a single NANA sequence.

The direct association of affinity with epitope cross-reactivity explains the enrichment of cross-reactive antibodies after repeated antigen-exposure and demonstrates the selective advantage of cross-reactive clones over time in the natural B cell affinity maturation process. The overall slower dissociation rate from NANP compared to the N-terminal junction peptides suggests that NANP affinity rather than NVDP- or NPDP-binding drove the selection of these antibodies in the human host (Batista and Neuberger, 1998; Foote and Eisen, 2000; Victora and Nussenzweig, 2012). The observed accumulation of class-switched *IGHV3-33*-, *IGKV1-5*-encoded cross-reactive antibodies with somatic mutations that are known to increase NANP affinity either directly or indirectly through homotypic antibody interactions provides additional support for this hypothesis (Imkeller et al., 2018; Murugan et al., 2018; Oyen et al., 2018).

Our data demonstrate that cross-reactive antibodies bind their respective PfCSP target epitopes in nearly identical conformations. Strikingly, independently of the antibodies’ Ig gene usage, cellular origin, binding mode, and binding profile, all antibodies with cross-reactivity to the N-terminal junction showed a strong preference for NANP and bias against NVDP as the core epitope motif independently of their binding profile. Together, the data indicate that NANP serves as the primary anchor motif for binding to PfCSP for cross-reactive and non-cross-reactive central repeat binders and drives the evolution of the response to the N-terminal junction. Preferences for NANP as primary anchor motif likely limit the number of possible recognition sites for cross-reactive antibodies on full-length PfCSP. Although we have no information on the binding mode of non-NANP-reactive NVDP binders or NVDP-, NPDP-cross-reactive antibodies, they were overall weak inhibitors independently of their affinity.

Our data identified a stringent minimal NANP-affinity threshold of K_D_ <10^-7^ M for parasite-inhibitory antibodies with IC_50_ values below 1 µg/ml, but we did not observe a direct correlation with Pf-inhibitory activity for antibodies with affinities below <10^-8^ M in the *in vitro* assay or the *in vivo* model. These findings suggest that high affinity is required but not sufficient to predict the potency of PfCSP antibodies as previously observed for mAb 1210 and a mutated version with 10-fold higher affinity, which showed no difference in *in vivo* inhibitory activity (Imkeller et al., 2018). The lack of correlation between antibody potency and affinity beyond a certain threshold, as well as the high serum titers above 50 µg/ml of the most potent antibodies that were required to mediate sterile or near sterile protection in our *in vivo* model likely reflects the complex and only partially understood role of PfCSP antibodies in parasite inhibition (Julien and Wardemann, 2019). A better understanding of how antibodies recognize PfCSP on the parasite surface and the mechanisms underlying their function will be essential to define more precise correlates of inhibitory activity and protection. Recent data show that the antibody-mediated precipitation of PfCSP from the surface of motile sporozoites in the skin leads to their death upon loss of the PfCSP coat via exposure to cytotoxic Pf molecules that are released in the skin to mediate host cell traversal and invasion (Aliprandini et al., 2018; Flores-Garcia et al., 2018). It remains to be determined whether PfCSP antibodies with different binding profiles vary in their mechanism of action and to what extent they inhibit the parasite not only in the skin, but also in the circulation or the liver.

The fact that mAbs with strong preference for either the NANP repeat (mab 317, (Oyen et al., 2017)) or N-terminal junction (mAb CIS43, (Kisalu et al., 2018)) showed similar levels of *in vivo* inhibitory activity compared to antibodies with broader binding profiles (mAbs 4493 and 2541) suggests that affinity rather than the antibody binding profile determines protection, and that the NANP repeat and the N-terminal junction represent potent target epitopes. Thus, novel vaccination strategies will likely benefit from inducing responses against both domains but not the non-protective epitopes in the PfCSP C terminus that are part of RTS,S (Scally et al., 2018). However, the success of any future vaccine design approach will not only depend on the design of the immunogen, but also on the number of targetable B cells that express PfCSP-reactive antibodies. Several independent studies have identified antibodies encoded by *IGHV3-33* and highly similar *IGHV3-30* genes after immunization with live or attenuated sporozoites as well as RTS,S, demonstrating that B cells expressing these antibodies are abundantly recruited into the PfCSP response (Kisalu et al., 2018; Murugan et al., 2018; Oyen et al., 2017; Tan et al., 2018). Especially, *IGHV3-33*-encoded antibodies are strongly enriched among high-affinity PfCSP binders (Murugan et al., 2018). Many of these are germline binders, but mutations that directly or indirectly increase affinity contribute to the overall high Pf-inhibitory activity of these antibodies. As exemplified here by mAb 2541, *IGHV3-33*-encoded antibodies especially in combination with IGKV1-5 light chains and 8-aa-long KCDR3s can be highly potent, suggesting that B cells expressing *IGHV3-33*-encoded antibodies represent a promising target population for any PfCSP vaccine design strategy. In the same studies, potent high-affinity non-*IGHV3-33*-encoded PfCSP antibodies such as mAbs 4493 and CIS43 were less frequently identified, suggesting that they are overall rare (Kisalu et al., 2018; Murugan et al., 2018; Oyen et al., 2017; Tan et al., 2018). These antibodies showed strong reactivity to the N-terminal junction, in contrast to the vast majority of *IGHV3-3*3-encoded antibodies with strong preference for the repeat. However, independently of the binding profile or Ig gene usage, the paratope core of all protective antibodies described in this and other studies preferentially recognized NANP in the context of extended NP/DP-NANP-NA/NV motifs. Thus, NANP motifs represent potent B cell epitopes that induce not only antibodies with preferential binding to the repeat, but also antibodies with high affinity to the N-terminal junction. We conclude that high affinity to the NP/DP-NANP-NA/NV motif is a feature of protective PfCSP antibodies independently of their binding profile. Future studies will have to determine whether immunogen-design strategies that aim at targeting the N-terminal junction in addition to the repeat region will be more efficient than RTS,S in promoting protective humoral immune responses to vaccination.

## AUTHOR CONTRIBUTIONS

R.M., S.W.S., G.C., and E.T. designed and conducted the experiments, interpreted experimental results, and wrote the paper. G.M. performed analyses and interpreted results. T.D., A.B., K.P. conducted experiments. E.A.L., J.-P.J., H.W conceived the study, designed and supervised the experiments, interpreted all results, and wrote the paper.

## COMPETING INTERESTS

All authors declare no conflicts of interest.

## ACKNOWLEDGEMENTS

The authors thank C. Canetta, J. Gaertner, A. Knauf, C. Winter, and D. Foster (German Cancer Research Center, Heidelberg), C. Kreschel, L. Spohr, D. Eyermann and M. Andres (Max Planck Institute for Infection Biology, Berlin) and the DKFZ/European Molecular Biology Laboratory (EMBL)/Heidelberg University Chemical Biology Core Facility, especially P. Sehr, for technical assistance and services. The following reagents were obtained from BEI Resources, NIAID, NIH: HC-04, Hepatocyte (human), MRA-975, contributed by J. S. Prachumsri.

## FUNDING

S.W.S was supported by a Hospital for Sick Children Lap-Chee Tsui Postdoctoral Fellowship and a Canadian Institutes of Health Research (CIHR) fellowship, and E.T. was supported by a CIHR Canada Graduate Scholarship - Master’s Award. This work was undertaken, in part, thanks to funding from the Bill and Melinda Gates Foundation (OPP1179906; J.-P.J, H.W. and E.A.L.) and the Canada Research Chairs program (J.-P.J.).

## FIGURE LEGENDS

**Supplemental Figure 1. Reactivity of human anti-PfCSP antibodies to different PfCSP peptides.**

ELISA-reactivity of monoclonal human antibodies to KQPA, NPDP, NVDP, NANP and C-CSP at the indicated concentrations. Red and green lines indicate positive (2A10; (Triller et al., 2017)) and negative controls (mGO53; (Wardemann et al., 2003)), respectively. n indicates the number of tested antibodies.

See also Figure 1.

**Supplemental Figure 2. Affinity of human anti-PfCSP antibodies to different PfCSP peptides.**

(A) Antibody affinities to the indicated PfCSP peptides and C-CSP measured by SPR.

(B) t-SNE clustering-based illustration of the affinity of PfCSP-reactive antibodies (n=200) to the indicated peptides and C-CSP measured by SPR for antibodies with ELISA AUC values >5.

See also Figure 2.

**Supplemental Figure 3. Ig gene usage of anti-PfCSP antibodies with different reactivity profiles and cross-reactivity of *IGHV3-33*-encoded antibodies.**

(A) *IGHV* and paired *IGKV* or *IGLV* gene usage of antibodies with the indicated binding profiles: NANP-specific, NANP, NVDP cross-reactive and NANP, NVDP, NPDP cross-reactive antibodies; C-CSP cross-reactive antibodies, and NVDP-specific and NVDP, NPDP cross-reactive antibodies. Antibodies using the same *IGHV* are indicated as solid lines around the pie charts. n in the pie chart centers indicate the number of antibodies.

(B) ELISA PfCSP-reactivity of monoclonal plasmablast antibodies (n=111; Murugan) tested at 4 µg/ml (left). The PfCSP reactivity of *IGHV3-33*-, *IGKV*1-5-encoded antibodies (n=6) was confirmed at different concentrations and is indicated as mean area under the curve (AUC; center). Red and green filled circles indicate the positive (2A10; (Triller et al., 2017)) and negative (mGO53; (Wardemann et al., 2003)) control mAbs, respectively. The binding profiles of the six PfCSP-reactive *IGHV3-33*-, *IGKV*1-5-encoded plasmablast antibodies to the indicated peptides and C-CSP are shown (right). Mean AUC values from three independent experiments are shown (center, right).

(C) Isothermal titration calometry (ITC) measurements of *IGHV3-33-*encoded antibodies binding to NANP_3_ (mAbs 1210, 2243, 2541, 3945, 4498) or a 14-aa long C-CSP peptide (PNRNVDENANANSA; mAb 3246).

(D) Detailed interactions between mAbs 2243 and 2541 and NANP_5,_ mAbs 4498 and 3945 and NANP_3_, and mAb 3246 and NANA. H-bonds are shown as black dashes.

See also Figure 3.

**Supplemental Figure 4. Delineation of PfCSP binding by mAb 4493 and comparison to other mAbs of reported structures.**

(A) ITC measurements of mAb 4493 with the indicated peptides.

(B) Superposition of NPDP peptides recognized by mAbs 4493 and CIS43 (Kisalu et al., 2018) show that the peptides are recognized in a similar U-shaped conformation, but by different angles of approach.

(C-E) Overlay of peptide conformations observed in co-complexes with the indicated antibodies, for which structures were reported with multiple peptides (CIS43, CIS42 (Kisalu et al., 2018); 311 (Oyen et al., 2017)). The NPDP, NVDP, and NANP peptides are colored in green, orange, and blue, respectively. Information about the positioning of these motifs in the antibody paratope is provided. Three distinct paratope positions (−1, 1 and 2) are indicated. Amino acid residues resolved in the structures are underlined.

See also Figure 4.

**Supplemental Figure 5. Comparison of the *in vivo* protective capacity of mAbs 4493, 317, and CIS43**

(A) Capacity of mAbs 317 (Oyen et al., 2017), CIS43 (Kisalu et al., 2018), 4493 and 1210 (Imkeller et al., 2018; Murugan et al., 2018)) to protect mice from blood-stage parasitemia after passive i.p. mAb transfer (300 µg/mouse) and exposure to the bites of mosquitoes infected with *PbPfCSP* parasites. The percentage of parasite-free mice is indicated. The C-CSP-reactive non-inhibitory mAb 1710 (Scally et al., 2018) was used as negative control. Pooled data from two independent experiments is shown. The total number of mice per group is indicated (n).

(B) Serum concentration of the transferred monoclonal antibodies in individual mice at the time of parasite challenge. Data shows the mean of at least two independent measurements. Red bars indicate mean values.

See also Figure 7.

**Supplemental Figure 6. Cross-reactivity of mAbs 2A10, 580 and 663 determined by SPR.**

Affinities of the chimeric mAb 2A10 with mouse variable (Zavala et al., 1983) and human IgG1 constant region (Triller et al., 2017), and of mAbs 663 and 580 (Triller et al., 2017) to the indicated PfCSP peptides measured by SPR.

## SUPPLEMENTAL TABLES

**Table S1. Ig gene features and reactivity of anti-PfCSP antibodies.**

**Table S2. Binding profiles of PfCSP-reactive monoclonal antibodies isolated from CD19-CD27+CD38+PfCSP-plasmablasts.**

**Table S3. Data collection and refinement statistics for *IGHV3-33*-encoded antibodies.**

**Table S4. Table of contacts between *IGHV3-33*-encoded antibodies and PfCSP subdomains.**

**Table S5. Data collection and refinement statistics for mAb 4493.**

**Table S6. Table of contacts between mAb 4493 and PfCSP subdomains.**

**Table S7. IC_50_ values of representative potent antibodies.**

**Table S8. *In vivo* potency of *IGVH3-33*-encoded mAbs at 300 μg to protect mice from mosquito-bite mediated *Plasmodium* infection.**

**Table S9. *In vivo* potency of non-*IGVH3-33*-encoded mAbs at 300 μg to protect mice from mosquito-bite mediated *Plasmodium* infection.**

**Table S10. *In vivo* potency of non-*IGVH3-33*-encoded mAbs at 150 μg to protect mice from mosquito-bite mediated *Plasmodium* infection.**

**Table S11. *In vivo* potency of non-*IGVH3-33* and *IGHV3-33*-encoded mAbs at 150 μg to protect mice from mosquito-bite mediated *Plasmodium* infection.**

## STAR METHODS

### CONTACT FOR REAGENT AND RESOURCE SHARING

Further information and requests for resources and reagents should be directed to and will be fulfilled by the Lead Contact, Hedda Wardemann.

### EXPERIMENTAL MODEL AND SUBJECT DETAILS

#### Cell lines

HEK293F cells were cultured according to the manufacturer’s instructions. The cells were passaged at 37 °C, 8% CO_2_ and 180 rpm in a 50 ml Bioreactor in FreeStyle™ 293-F medium. HC-04 cells (MRA-975, deposited by Jetsumon Sattabongkot; Sattabongkot et al., 2006) were cultured at 37 °C and 5% CO2 using HC-04 complete culture medium (428.75 ml MEM (-L-glu), 428.75 ml F-12 Nutrient Mix (+ L-glu), 15 mM HEPES, 1.5 g/l NaHCO3, 2.5 mM L-glutamine and 10% FCS).

#### Bacteria

MAX Efficiency^®^ DH10B™ Competent Cells were cultured at 37 °C and 180 rpm in LB medium for maintenance and Terrific broth for plasmid production.

#### Plasmodium falciparum cultures

*Plasmodium falciparum Pf* NF54 (a kind gift of Prof. R. Sauerwein) were cultured in O+ human red blood cells at 37°C, 4% CO_2_ and 3% O_2_ in a Heracell 150i Tri-gas incubator (Thermo Scientific). For gametocyte production, asynchronous parasite cultures were diluted to 1% parasitaemia and maintained for 15-16 days with daily change of RPMI-1640 medium (Thermo Scientific) supplemented with 10% human A+ serum and 10 mM hypoxantine (c-c-Pro) until mosquito infections.

*Pb-PfCSP,* a replacement *P. berghei* line expressing *Pf* CSP (NF54) under the control of the *Pb csp* regulatory sequences (Triller et al. 2017), was obtained from Chris J. Janse and Shahid M. Khan and passaged every 3-4 days in CD1 female mice.

#### Mosquitoes

All mosquitoes were kept at 28-30 °C and 70-80% humidity. *Anopheles coluzzii* Ngousso S1 strain (Harris et al., 2010) were used for the production of *Pf* NF54 sporozoites for *in vitro* traversal assays. *A. gambiae* 7b line, immunocompromised transgenic mosquitoes derived from the G3 laboratory strain (Pompon and Levashina, 2015), were used for the production of *Pb-PfCSP* sporozoites for *in vivo* infections.

#### Mice

Female C57BL/6 mice (7-9 weeks old) and female CD-1 mice (8-12 weeks old) were bred in the MPIIB Experimental Animal Facility (Marienfelde, Berlin), handled in accordance with the German Animal Protection Law (§8 Tierschutzgesetz) and approved by the Landesamt für Gesundheit und Soziales (LAGeSo), Berlin, Germany (project numbers 368/12 and H0335/17).

### METHOD DETAILS

#### Ig gene cloning and recombinant antibody production

Ig heavy and light chain genes corresponding to antibody were cloned into human Igγ1 (AbVec2.0-IGHG1, Genbank ID: LT615368) and Igκ (AbVec1.1-IGKC, Genbank ID: LT615369) or Igλ (AbVec1.1-IGLC2-XhoI) expression vectors, respectively (Tiller et al., 2009). The cloning vectors are available from Addgene (Catalog number: 80795, 80796 and 99575). In brief, restriction site-tagged specific V and J-gene primers were used for amplifying Ig genes from single B cells and the amplicons were cloned into the above-mentioned vectors. Ig genes of mAbs CIS43 and 317 were obtained by reverse translation of the protein sequences deposited (PDB accession number 6B5M for CIS43 (Kisalu et al., 2018) and 6AXL for 317 (Oyen et al., 2017)). Ig genes of CIS43 were synthesized at MWG Eurofins Genomics with AgeI restriction site at the 5’ end and SalI and BsiWI restriction sites at the 3’end of the heavy and kappa Ig genes, respectively. Ig genes of 317 were synthesized at GeneArt (Thermofisher) and restriction sites were introduced via PCR. Upon successful cloning, recombinant monoclonal antibodies were expressed in HEK293F cells (ThermoFisher Scientific).

#### Enzyme-Linked Immunosorbent Assay

Recombinant monoclonal antibodies were purified using Protein G Sepharose beads (GE healthcare) and the IgG concentration was measured by ELISA as described (Tiller et al., 2008). Antigen ELISAs were performed as described (Triller et al., 2017). In brief, high-binding 384-well polystyrene plates (Corning) were coated overnight at 4 °C with KQPA, NPDP, NVDP, C-CSP or Streptavidin at 50 ng/well or PfCSP at 40 ng/well in 25 µl. Streptavidin-coated plates were incubated for 1 h with 200 ng/well biotinylated NANP5.5 in 25 µl. Plates were washed 3 times with 0.05% Tween 20 in PBS, blocked with 50 µl of 1% BSA in PBS for 1h at room temperature (RT), and washed again prior to incubation with diluted monoclonal antibodies at the indicated concentrations for 1.5 h at RT. Wells were washed and incubated with goat anti-human IgG-HRP at 1:1000 (Jackson Immuno Research) in PBS with 1% BSA. One-step ABTS substrate (RT, 20 µl/well; Roche) and 1X KPL ABTS® peroxidase stop solution (RT, 20 µl/well; SeraCare Life Sciences, Inc.) were used for detection. A chimeric version of the murine anti-PfCSP antibody 2A10 (Triller et al., 2017) with human Ig heavy and Ig kappa constant regions and the non-PfCSP-reactive antibody mGO53 (Wardemann et al., 2003) were used as a positive control and negative control, respectively. ELISA area under the curve (AUC) values were calculated using GraphPad Prism 7.04 (GraphPad).

#### Surface plasmon resonance (SPR)

SPR measurements were performed with a BIACORE T200 (GE Healthcare) as described (Murugan et al., 2018). In brief, the instrument was docked with a series S sensor chip CM5 (GE Healthcare). 10 mM HEPES with 150 mM NaCl at pH 7.4 was used as running buffer. All samples were immobilized by amine coupling using the human antibody capture kit (GE Healthcare) according to the manufacturer’s instructions. Sample antibodies and the non-PfCSP-reactive negative control antibody mGO53 (Wardemann et al., 2003) were captured in the sample and reference flow cell at equal concentrations, respectively. Flow cells were stabilized with running buffer at 10 µl/min flowrate for 20 min. The respective peptides were dissolved in running buffer and injected at 0, 0.015, 0.09, 0.55, 3.3, and 20 µM concentration. A flow rate of 30 µl/min was maintained, allowing the association and dissociation of the peptides for 60 s and 180 s respectively, at 25 °C. For high affinity antibodies (∼10^-10^ M), additional measurements at 0, 0.42, 2.57, 15.43, 92.6 and 555.5 nM concentrations were performed. After each run, both flow cells were regenerated with 3 M MgCl_2_. The data were fit using 1:1 binding model or steady state kinetic analysis using the BIACORE T200 software V2.0.

#### Pf sporozoite hepatocyte traversal assay

*Anopheles coluzzii* mosquitoes were infected with mature Pf gametocytes (NF54 strain) via artificial midi-feeders (Glass Instruments, The Netherlands) for 15 min and kept at 26°C and 80% humidity in a controlled S3 facility in accordance with local safety authorizations (Landesamt für Gesundheit und Soziales Berlin, Germany, LAGeSo, project number 411/08). Infected mosquitoes received an additional uninfected blood meal 7-8 days post infection (dpi) and were collected 13-15 dpi to isolate sporozoites. Sporozoites were isolated in HC-04 medium by dissecting and grinding mosquito thoraces containing salivary glands with glass pestles, followed by filtering the extracts with 100 µm and 40 µm cell strainers. The isolated salivary gland sporozoites were enumerated in a hemocytometer (Malassez, Marienfelde) and used for traversal assays as previously described (Triller et al., 2017). Briefly, salivary gland Pf sporozoites in HC-04 medium were pre-incubated with 100 µg/ml or serial dilutions (0.032, 0.16, 0.8, 4 and 20 µg/ml) of monoclonal antibodies in 27.5 µl for 30 min on ice and added to human hepatocytes (HC-04, (Sattabongkot et al., 2006)) for 2 h at 37 °C and 5% CO_2_ in the presence of 0.5 mg/ml dextran-rhodamine (Molecular Probes). Cells were washed and fixed with 1% PFA in PBS before measuring dextran positivity using FACS LSR II instrument (BD Biosciences). Data analysis was performed by subtraction of the background (dextran positivity in cells treated with uninfected mosquito salivary gland material) and normalization to the maximum Pf traversal capacity (dextran positivity in cells treated with salivary gland Pf sporozoites) using FlowJo V.10.0.8 (Tree Star). A chimeric humanized version of the PfCSP-reactive monoclonal antibody 2A10 (Triller et al., 2017) and of the non-PfCSP-reactive monoclonal antibody mGO53 (Wardemann et al., 2003) was used as a positive and negative control, respectively. IC_50_ values were calculated for each antibody by four-parameter logistic curve fitting in GraphPad Prism 7.04 (GraphPad) using the measurements from at least three independent experiments.

#### Fab production

Fabs of mAbs 2243, 4498, and 3945 were generated by papain digestion of IgG, purified via Protein A chromatography (GE Healthcare) followed by cation-exchange chromatography (MonoS, GE Healthcare) and size-exclusion chromatography (Superdex 200 Increase 10/300 GL, GE Healthcare). Fabs of mAbs 3246, 2541 and 4493 were generated by cloning of the *IGH* and *IGK* variable region gene segments into pcDNA3.4 TOPO expression vectors immediately upstream of human *CH1 and IGK* constant regions, respectively, followed by transient expression in HEK293F cells (Thermo Fisher Scientific) and purification via KappaSelect affinity chromatography (GE Healthcare), cation-exchange chromatography (MonoS, GE Healthcare) and size-exclusion chromatography (Superdex 200 Increase 10/300 GL, GE Healthcare).

#### Crystallization and structure determination

Purified Fabs of mAb 2243 and 2541 were mixed with NANP_5_ in a 3:1 molar ratio and excess Fab was purified away via size exclusion chromatography (Superdex 200 Increase 10/300 GL, GE Healthcare). Purified Fab 2243-NANP_5_ and 2541-NANP_5_ complexes were then concentrated to 10 mg/mL prior to crystallization trials. Purified 3246 Fab was concentrated to 9 mg/mL and diluted to 8 mg/mL with NANA (10 mg/mL), prior to crystallization trials. To enhance crystallizability, purified 4498, 3945 and 4493 Fabs were mixed with the anti-kappa V_H_H domain (Thermofisher) in a 1:2 molar ratio and excess V_H_H domain was removed away via size exclusion chromatography (Superdex 200 Increase 10/300 GL, GE Healthcare). The anti-kappa V_H_H domain has previously been shown to enhance the crystallizability of Fabs (Ereño-Orbea et al., 2018). Fab-V_H_H co-complexes were concentrated to 6 mg/mL and diluted to 5 mg/mL with the respective peptide. 2243-NANP_5_ co-crystals grew in 20 % (w/v) PEG 3350, 0.2 M ammonium sulfate and were cryoprotected in 15 % (w/v) ethylene glycol. 2541-NANP_5_ co-crystals grew in 1.8 M ammonium sulfate, 0.1 M citric acid pH 3.6 after microseeding from thin, layered plate-like crystals that grew in 1.6 M ammonium sulfate, 0.1 M citric acid pH 4 and were cryoprotected in 25 % (w/v) glycerol. 4498-V_H_H-NANP_3_ crystals grew in 10 % (w/v) 2-methyl-2,4-pentanediol, 0.1 M citric acid pH 2.5 and were cryoprotected in 15 % (w/v) ethylene glycol. 4498-V_H_H-NDN_3_ crystals grew in 0.1 M phosphate-citrate pH 4.2, 20 % (w/v) PEG 8000, 0.2 M sodium chloride and were cryoprotected in 20 % (w/v) glycerol. 3945-V_H_H-NANP_3_ crystals grew in 0.2 M di-sodium hydrogen phosphate, 20 % (w/v) PEG 3350 and were cryoprotected in 15 % (w/v) ethylene glycol. 3246-NANA crystals grew in 10 % (v/v) isopropanol, 20 % (w/v) PEG4000, 0.1 M HEPES pH 7.5 and 10 % (w/v) ethylene glycol. 4493-V_H_H-KQPA crystals grew in 0.1 M HEPES pH 7.0, 1 M lithium chloride, 20 % (w/v) PEG 6000 and were cryoprotected in 15 % (w/v) ethylene glycol. 4493-V_H_H-NPDP crystals grew in 20 % (w/v) PEG 3350, 0.2 M sodium nitrate and were cryoprotected in 15 % (w/v) ethylene glycol. 4493-V_H_H-NDN_3_ crystals grew in 20 % (w/v) PEG 3350, 0.2 M potassium nitrate and were cryoprotected in 15 % (w/v) ethylene glycol. 4493-V_H_H-DND_3_ crystals grew in 20 % (w/v) PEG 8000, 0.1 M MES pH 6.0 and 0.2 M calcium acetate and were cryoprotected in 20 % (w/v) glycerol. 4493-V_H_H-NANP_3_ grew in 40 % (w/v) PEG 600, 0.1 M sodium citrate pH 5.5 after microseeding from thin needle-like crystals that grew in 0.1 M HEPES pH 7.5, 20 % (w/v) PEG 8000 and were cryoprotected in 15 % (w/v) ethylene glycol. Data were collected at the 08ID-1 beamline at the Canadian Light Source (CLS), the 23-ID beamline at the Advanced Photon Source (APS), or the 17-ID-2 beamline at the National Synchrotron Light Source II (NSLS-II), and processed and scaled using XDS (Kabsch et al., 2010). The structures were determined by molecular replacement using Phaser (McCoy et al., 2007). Refinement of the structures was carried out using phenix.refine (Adams et al., 2010) and iterations of refinement using Coot (Emsley et al., 2010). Software were accessed through SBGrid (Morin et al., 2013).

#### Isothermal titration calorimetry

Calorimetric titration experiments were performed with an Auto-iTC200 instrument (Malvern) at either 15 °C or 25 °C. Proteins were dialyzed against 20 mM Tris pH 8.0 and 150 mM sodium chloride overnight at 4 °C. Fabs were concentrated to 10 µM and added to the calorimetric cell, which was titrated with peptide (100 µM) in 15 successive injections of 2.5 µl. Experiments were performed at least twice, and the mean and standard error of the mean were reported (Fig. 3G, 4A, S3C and S4A). The experimental data were analyzed according to a 1:1 binding model by means of Origin 7.0.

#### Plasmodium berghei infections in vivo

*A. gambiae* 7b mosquitoes were fed on female CD-1 mice infected with *Pb-PfCSP* parasites (0.1-0.8% gametocytemia) and kept at 20 °C and 80% humidity until further usage. Infected mosquitoes were offered an additional uninfected blood meal 7 dpi and 20 mosquitoes were dissected for oocyst counts 17 dpi. Female C57BL/6 mice were passively immunized by i.p. injection of 150 or 300 µg of monoclonal antibodies in 200 µl of PBS. After 20 h (18 dpi), mice were exposed to *Pb-PfCSP*-infected mosquitoes (infection prevalence between 75% and 100%; Supplemental tables S8-S11). All blood-fed mosquitoes were collected individually for gDNA extraction (NucleoMag VET, Macherey-Nagel) followed by PCR to determine their *Pb-PfCSP* infectivity status. Specific primers amplifying *P. berghei* 18s RNA gene (Friesen et al. 2010) and control primers amplifying *A. gambiae* AGAP001076 gene (Gildenhard et al. 2019) were used. Mosquitoes positive for both PCR reactions were considered infected. Antibody titers were measured by ELISA in serum samples collected from the submandibular vein 2-3 h post mosquito bite. Blood parasitaemia was assayed by daily tests of a minimum of 100 microscopic fields per Giemsa-stained thin blood smears 3 - 7 days and 10 days post mosquito bite. Infected mice were sacrificed two days after the detection of parasitaemia.

#### Statistics

Statistics was performed on Prism 7.04 (GraphPad) or RStudio (version 3.2.2) using two-tailed Mann-Whitney assuming non-normal distribution or Mantel-Cox log-rank test for *in vivo* experiments, as described in the figure legends. P values less than 0.05 were considered significant (*P < 0.05; **P < 0.01; ***P < 0.001; ****P < 0.0001) as indicated in the figure legends.

#### Data and Software Availability

The data that support the findings of this report are available from the corresponding authors upon reasonable request. The crystal structures reported in this manuscript have been deposited in the Protein Data Bank, www.rcsb.org (PDB ID codes 6O23, 6O24, 6O25, 6O26, 6O28, 6O29, 6O2A, 6O2B, 6O2C, 6ULE, 6ULF).

