## Supplemental Figures and Tables for "Evolution of protective human antibodies against *Plasmodium falciparum* circumsporozoite protein repeat motifs"

### Figure S1

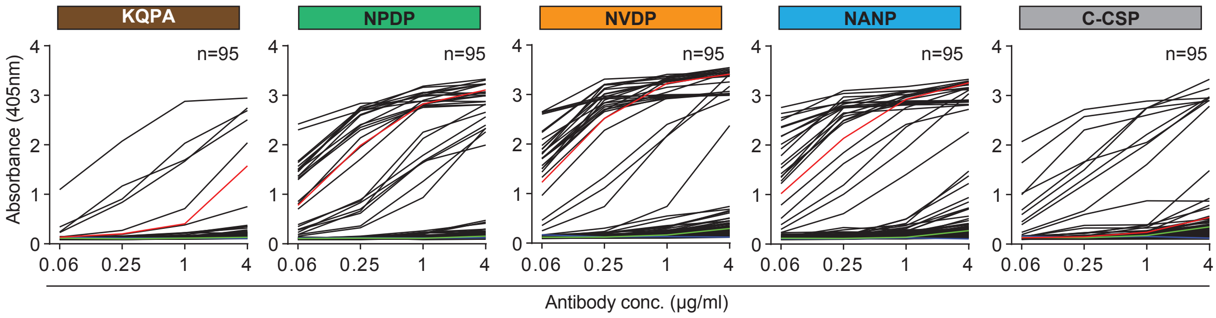
KQPA
NPDP
NVDP
NANP
C-CSP
n=95
n=95
n=95
n=95
n=95
Absorbance (405nm)
0
1
2
3
4
0.06
0.25
1
4
Antibody conc. (µg/ml)

### Figure S2

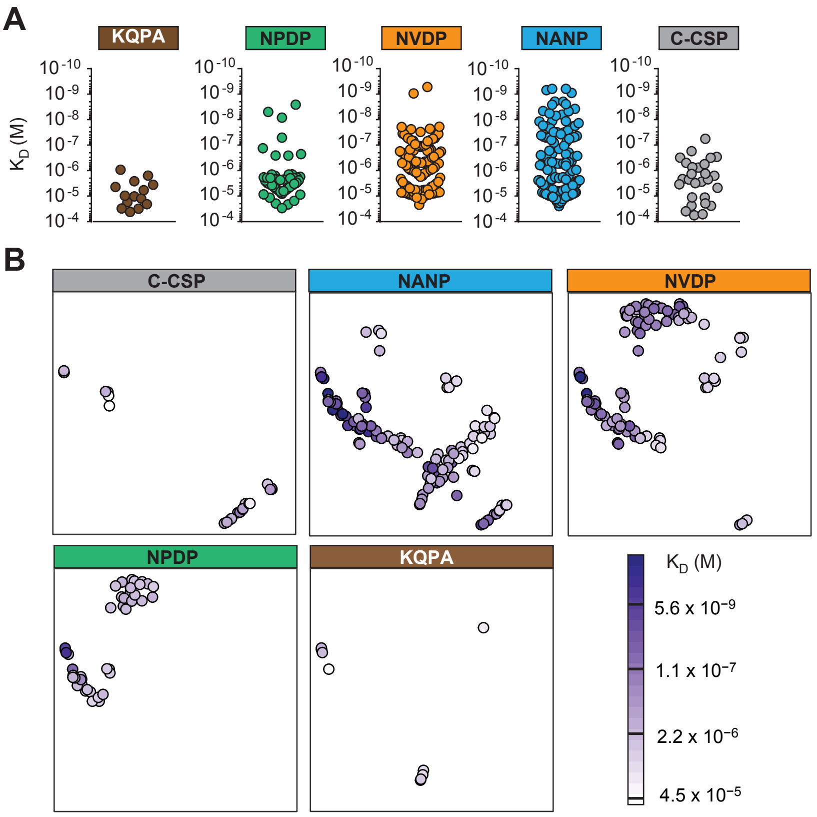
A
KQPA
NPDP
NVDP
NANP
C-CSP
$K_D$ (M)
$10^{-10}$
$10^{-9}$
$10^{-8}$
$10^{-7}$
$10^{-6}$
$10^{-5}$
$10^{-4}$
B
C-CSP
NANP
NVDP
NPDP
KQPA
$K_D$ (M)
$5.6 \times 10^{-9}$
$1.1 \times 10^{-7}$
$2.2 \times 10^{-6}$
$4.5 \times 10^{-5}$

#### Figure S3

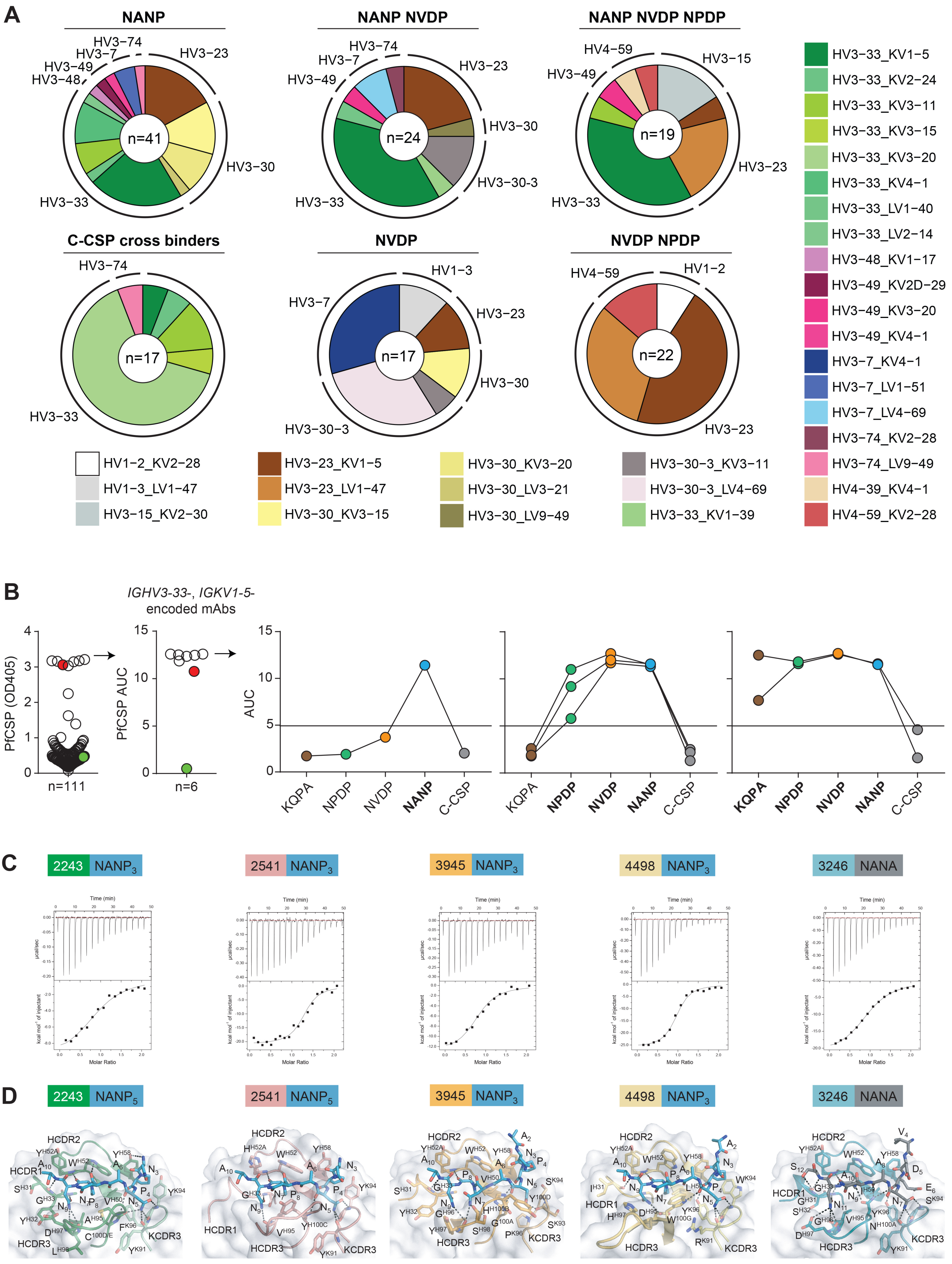

A
NANP
NANP NVDP
NANP NVDP NPDP
C-CSP cross binders
NVDP
NVDP NPDP
n=41
n=24
n=19
n=17
n=17
n=22
HV3−74
HV3−7
HV3−49
HV3−48
HV3−23
HV3−30
HV3−33
HV3−30-3
HV4−59
HV3−15
HV1−3
HV1−2
HV1−2_KV2−28
HV1−3_LV1−47
HV3−15_KV2−30
HV3−23_KV1−5
HV3−23_LV1−47
HV3−30_KV3−15
HV3−30_KV3−20
HV3−30_LV3−21
HV3−30_LV9−49
HV3−30−3_KV3−11
HV3−30−3_LV4−69
HV3−33_KV1−39
HV3−33_KV1−5
HV3−33_KV2−24
HV3−33_KV3−11
HV3−33_KV3−15
HV3−33_KV3−20
HV3−33_KV4−1
HV3−33_LV1−40
HV3−33_LV2−14
HV3−48_KV1−17
HV3−49_KV2D−29
HV3−49_KV3−20
HV3−49_KV4−1
HV3−7_KV4−1
HV3−7_LV1−51
HV3−7_LV4−69
HV3−74_KV2−28
HV3−74_LV9−49
HV4−39_KV4−1
HV4−59_KV2−28
B
IGHV3-33-, IGKV1-5-encoded mAbs
PfCSP (OD405)
PfCSP AUC
AUC
n=111
n=6
KQPA
NPDP
NVDP
NANP
C-CSP
C
2243 NANP3
2541 NANP3
3945 NANP3
4498 NANP3
3246 NANA
Time (min)
μcal/sec
kcal mol⁻¹ of injectant
Molar Ratio
D
2243 NANP5
2541 NANP5
3945 NANP3
4498 NANP3
3246 NANA
HCDR1
HCDR2
HCDR3
KCDR3

#### Figure S4

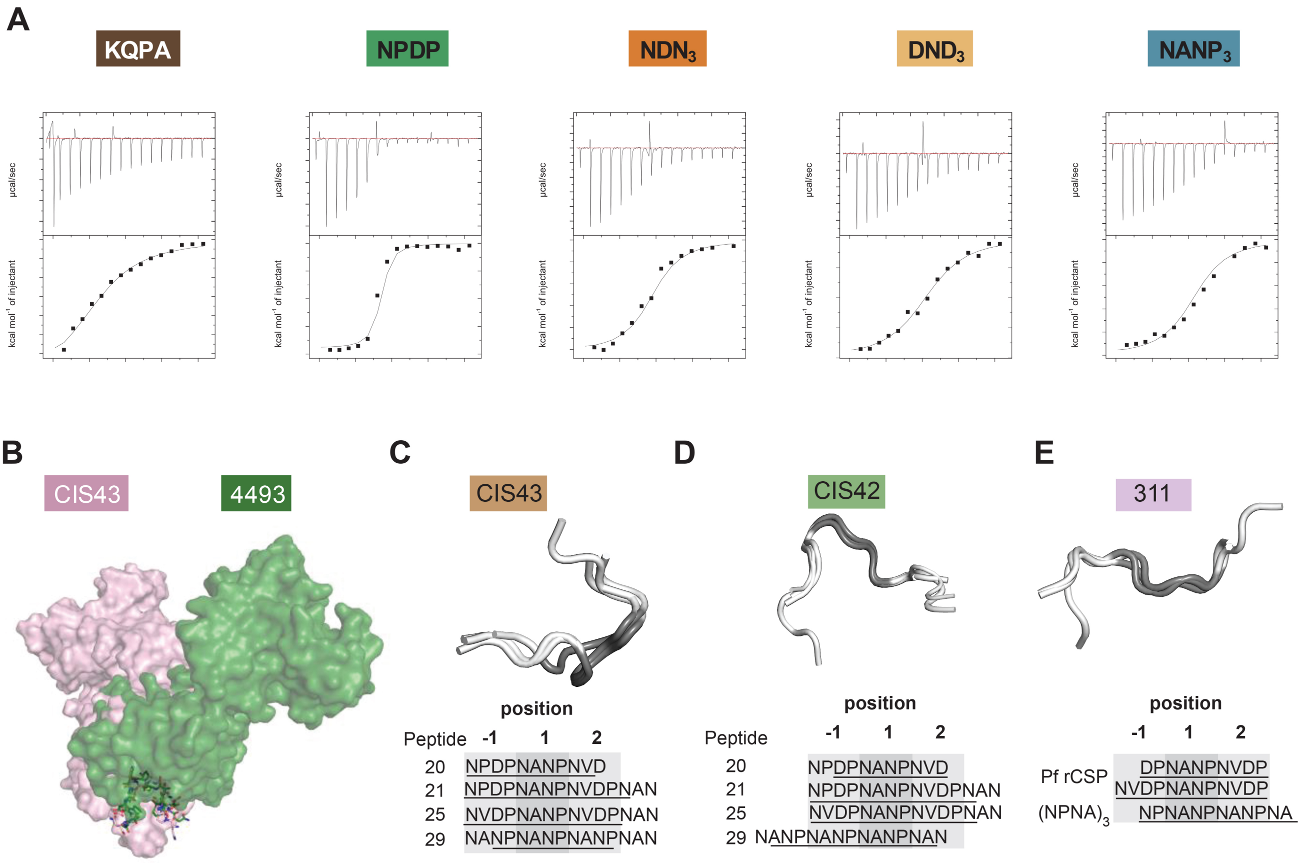
A
KQPA
NPDP
NDN3
DND3
NANP3
µcal/sec
kcal mol-1 of injectant
B
CIS43
4493
C
CIS43
position
Peptide -1 1 2
20 NPDPNANPNVD
21 NPDPNANPNVDPNAN
25 NVDPNANPNVDPNAN
29 NANPNANPNANPNAN
D
CIS42
position
Peptide -1 1 2
20 NPDPNANPNVD
21 NPDPNANPNVDPNAN
25 NVDPNANPNVDPNAN
29 NANPNANPNANPNAN
E
311
position
-1 1 2
Pf rCSP DPNANPNVDP
NVDPNANPNVDP
(NPNA)3 NPNANPNANPNA

#### Figure S5

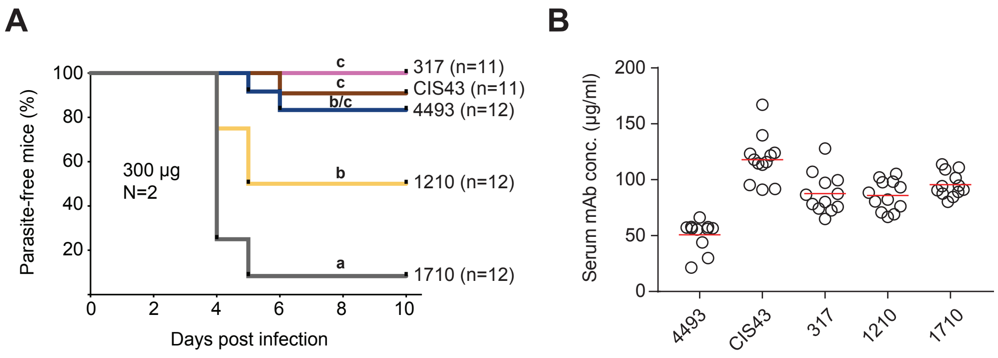

A
Parasite-free mice (%)
100
80
60
40
20
0
2
4
6
8
10
Days post infection
300 µg
N=2
c
c
b/c
b
a
317 (n=11)
CIS43 (n=11)
4493 (n=12)
1210 (n=12)
1710 (n=12)
B
Serum mAb conc. (µg/ml)
200
150
100
50
0
4493
CIS43
317
1210
1710

### Figure S6

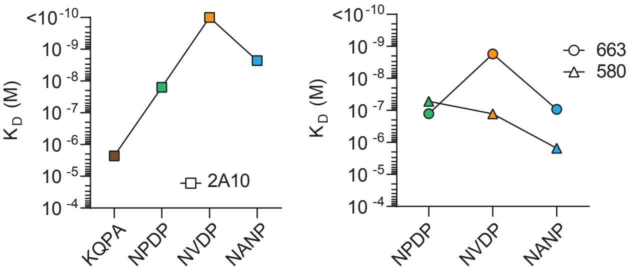

K_D (M)
<10^-10
10^-9
10^-8
10^-7
10^-6
10^-5
10^-4
2A10
KQPA
NPDP
NVDP
NANP
663
580

### Table S1. Ig gene features and reactivity of anti-PfCSP antibodies.

| Antibody name | donor | Pf exposure | Ig gene information | | | | | | | | | | ELISA AUC | | | | | SPR KD (M) | | | | | Pf sporozoite Inhibition (%) |
|---|---|---|---|---|---|---|---|---|---|---|---|---|---|---|---|---|---|---|---|---|---|---|---|
| | | | *IGHV* | *IGHD* | *IGHJ* | light | *IGKV/ IGLV* | *IGKJ/ IGLJ* | *IGHC* | cluster id | IGHV SHM | IGKV/IGLV SHM | KQPA | NPDP | NVDP | NANP | C-CSP | KQPA | NPDP | NVDP | NANP | C-CSP | |
| L1517x0075 | 17 | I | *4-59* | *6-13* | *6* | kappa | *3-20* | *1* | *IGHG2* | 19 | 4 | 3 | 2.4 | 3.3 | 2.5 | 0.5 | 1.3 | | > E-04 | > E-04 | > E-04 | | |
| L1517x0521 | 17 | II | *4-59* | *2-15* | *4* | lambda | *1-47* | *3* | *IGHM* | 0 | 7 | 5 | 0.7 | 0.8 | 0.8 | 0.5 | 1.4 | | | | | | |
| L1517x0618 | 17 | III | *3-33* | *3-22* | *4* | kappa | *1-12* | *4* | *IGHM* | 0 | 7 | 3 | 1.6 | 1.5 | 1.9 | 0.5 | 1.3 | | | | 1.2E-05 | > E-04 | |
| L1517x0692 | 17 | III | *3-33* | *1-1* | *4* | kappa | *4-1* | *1* | *IGHM* | 0 | 0 | 0 | 1.5 | 1.4 | 1.6 | 2.4 | 7.4 | | | | 3.5E-06 | 3.1E-06 | 26.9 |
| L1517x0622 | 17 | III | *3-74* | *2-2* | *4* | lambda | *1-47* | *1* | *IGHM* | 0 | 8 | 6 | 1.3 | 1 | 1.2 | 1.7 | 0.8 | | | | 9.3E-07 | | |
| L1426x0864 | 26 | III | *3-7* | *7-27* | *4* | kappa | *2-30* | *2* | *IGHM* | 0 | 1 | 0 | 2 | 1.7 | 1.6 | 0.6 | 1.3 | | | | 9.1E-06 | | |
| L1435x1741 | 35 | C | *4-39* | *6-19* | *4* | kappa | *4-1* | *1* | *IGHG1* | 0 | 26 | 8 | 1.4 | 6.2 | 9.7 | 7.4 | 0.8 | | | 2.1E-07 | 1.8E-06 | | 79 |
| D0835x0465 | 35 | I | *3-33* | *6-19* | *4* | kappa | *2-24* | *1* | *IGHM* | 0 | 6 | 0 | 2 | 2 | 2 | 7.9 | 8.1 | | | 2.2E-05 | 6.8E-06 | 2.7E-06 | 59.3 |
| L1435x2690 | 35 | III | *3-30-3* | *1-26* | *4* | kappa | *3-11* | *1* | *IGHM* | 0 | 4 | 1 | 1.6 | 1.5 | 1.9 | 3.6 | 1.2 | | | | 7.3E-06 | | |
| L1435x2867 | 35 | III | *3-30-3* | *6-19* | *4* | kappa | *3-11* | *1* | *IGHM* | 0 | 2 | 0 | 1.5 | 2 | 6.2 | 4.9 | 1.1 | | 1.1E-05 | 6.9E-06 | 1.2E-05 | | 67.7 |
| L1435x2991 | 35 | III | *3-33* | *3-22* | *4* | kappa | *3-20* | *2* | *IGHM* | 0 | 0 | 0 | 2.2 | 1.9 | 1.7 | 2.8 | 1.7 | | | | 1.6E-05 | | |
| L1435x3008 | 35 | III | *3-33* | *4-17* | *4* | kappa | *1-5* | *1* | *IGHM* | 0 | 0 | 0 | 1.9 | 1.9 | 3.1 | 6.8 | 1.6 | | | | 1.2E-05 | | 54.6 |
| L1542x1690 | 42 | III | *3-33* | *1-7* | *4* | kappa | *2-24* | *1* | *IGHM* | 20 | 12 | 5 | 1.6 | 1.5 | 1.4 | 0.4 | 2.1 | | | | 1.0E-05 | | |
| L1542x1837 | 42 | III | *3-33* | *7-27* | *4* | kappa | *2-24* | *1* | *IGHM* | 19 | 9 | 7 | 1.9 | 1.7 | 1.5 | 0.5 | 2.6 | | | | 1.3E-05 | | |
| L1542x1606 | 42 | III | *3-30* | *1-26* | *4* | lambda | *2-14* | *2* | *IGHM* | 14 | 1 | 1 | 1.3 | 1.2 | 1.2 | 1.3 | 1.6 | | | | 1.7E-05 | | |
| L1542x1769 | 42 | III | *3-30* | *4/OR15-4a* | *3* | kappa | *3-15* | *2* | *IGHM* | 12 | 8 | 3 | 1.3 | 1.5 | 1.6 | 5 | 1.3 | | | | 1.6E-05 | | |
| L1542x1811 | 42 | III | *3-30* | *4/OR15-4a* | *3* | kappa | *3-15* | *2* | *IGHM* | 12 | 7 | 3 | 1.3 | 1.3 | 1.3 | 1.2 | 1.1 | | | | 1.3E-05 | | |
| L1542x1238 | 42 | II | *3-30* | *1-1* | *4* | kappa | *3-15* | *1* | *IGHM* | 11 | 7 | 6 | 2.5 | 2.7 | 6.8 | 3.3 | 1.8 | | | 4.9E-06 | > E-04 | | 62.4 |
| L1542x1539 | 42 | III | *3-30* | *1-1* | *4* | kappa | *3-15* | *1* | *IGHM* | 11 | 7 | 6 | 2.5 | 2.5 | 5 | 2.6 | 1.4 | | > E-04 | | > E-04 | | |
| L1542x1773 | 42 | III | *3-30* | *1-1* | *4* | kappa | *3-15* | *1* | *IGHM* | 11 | 7 | 6 | 3 | 3.1 | 5.9 | 1 | 2.1 | | | | > E-04 | | 48.1 |
| L1542x1565 | 42 | III | *3-30* | *2-8* | *4* | kappa | *3-15* | *1* | *IGHM* | 10 | 8 | 4 | 1.4 | 1.3 | 1.2 | 6.5 | 1.3 | | | | 2.8E-06 | | 63.5 |
| L1542x1633 | 42 | III | *3-30* | *2-8* | *4* | kappa | *3-15* | *1* | *IGHM* | 10 | 8 | 4 | 1.3 | 1.3 | 1.2 | 6.9 | 1.8 | | | | 1.8E-05 | | 67.4 |
| L1542x1648 | 42 | III | *3-30* | *2-8* | *4* | kappa | *3-15* | *1* | *IGHM* | 10 | 8 | 4 | 1.3 | 1.2 | 1.1 | 8.1 | 1.7 | | | | 1.9E-05 | | 66.6 |
| L1542x1671 | 42 | III | *3-30* | *2-8* | *4* | kappa | *3-15* | *1* | *IGHM* | 10 | 8 | 4 | 1.8 | 1.7 | 1.5 | 6.3 | 1.5 | | | | 5.3E-06 | | 46.9 |
| L1542x1746 | 42 | III | *3-30* | *2-8* | *4* | kappa | *3-15* | *1* | *IGHM* | 10 | 8 | 4 | 1.4 | 1.1 | 1.4 | 5.8 | 1.2 | | | | 3.8E-06 | | 37.3 |
| L1542x2133 | 42 | C | *3-23* | *6-19* | *4* | lambda | *2-14* | *2* | *IGHM* | 7 | 11 | 6 | 1.4 | 2.6 | 2.1 | 0.4 | 1.8 | | | 8.6E-06 | 1.0E-05 | | |
| L1542x2151 | 42 | C | *3-33* | *6-19* | *4* | kappa | *1-5* | *4* | *IGHM* | 0 | 0 | 1 | 1.6 | 1.9 | 2.4 | 9.7 | 1.2 | | | | 6.4E-07 | | 73.2 |
| L1542x2159 | 42 | C | *3-33* | *1-26* | *4* | kappa | *3-11* | *2* | *IGHM* | 0 | 0 | 0 | 1.6 | 1.6 | 1.8 | 2.8 | 1.3 | | | | 8.1E-06 | | |
| L1542x2207 | 42 | C | *3-33* | *3-16* | *3* | kappa | *1-5* | *1* | *IGHM* | 0 | 3 | 3 | 2 | 4.4 | 11.8 | 12.6 | 1.5 | | 6.3E-06 | 4.4E-07 | 5.6E-08 | | 94.3 |
| L1542x1955 | 42 | C | *3-33* | *6-19* | *2* | lambda | *1-40* | *1* | *IGHM* | 0 | 1 | 0 | 2 | 2.1 | 8.4 | 12.7 | 1.5 | | 8.5E-06 | 2.8E-06 | 1.7E-07 | | 87.2 |
| L1542x1369 | 42 | II | *3-30-3* | *1-26* | *3* | kappa | *3-15* | *1* | *IGHM* | 0 | 2 | 0 | 1.5 | 1.6 | 1.6 | 0.4 | 1.1 | | | | > E-04 | | |

| Antibody name | donor | Pf exposure | Ig gene information | | | | | | | | | | ELISA AUC | | | | | SPR KD (M) | | | | | Pf sporozoite Inhibition (%) |
|---|---|---|---|---|---|---|---|---|---|---|---|---|---|---|---|---|---|---|---|---|---|---|---|
| | | | *IGHV* | *IGHD* | *IGHJ* | light | *IGKV/ IGLV* | *IGKJ/ IGLJ* | *IGHC* | cluster id | IGHV SHM | IGKV/IGLV SHM | KQPA | NPDP | NVDP | NANP | C-CSP | KQPA | NPDP | NVDP | NANP | C-CSP | |
| L1542x1571 | 42 | III | *3-33* | *6-13* | *4* | kappa | *4-1* | *1* | *IGHM* | 0 | 0 | 0 | 1.9 | 1.6 | 1.7 | 8.9 | 4.2 | | | | 8.2E-06 | | 44.2 |
| L1542x1578 | 42 | III | *3-33* | *1-1* | *6* | kappa | *1-5* | *2* | *IGHM* | 0 | 0 | 0 | 2 | 2.4 | 5.9 | 11.3 | 1.7 | | | 1.2E-05 | 4.9E-07 | | 69.7 |
| L1542x1601 | 42 | III | *3-33* | *2-2* | *5* | kappa | *1-5* | *4* | *IGHM* | 0 | 0 | 0 | 1.4 | 1.3 | 1.3 | 0.5 | 0.9 | | | | 1.1E-05 | | |
| L1542x1616 | 42 | III | *3-33* | *6-19* | *4* | kappa | *4-1* | *1* | *IGHM* | 0 | 0 | 3 | 1.5 | 1.3 | 1.2 | 4.3 | 2.9 | | | | 1.8E-06 | | |
| L1542x1744 | 42 | III | *3-49* | *4-17* | *4* | kappa | *2D-29* | *2* | *IGHM* | 0 | 1 | 0 | 8.9 | 2.1 | 2.6 | 12.9 | 1.2 | 4.4E-06 | 3.3E-06 | 5.5E-06 | 2.6E-07 | | |
| L1542x1755 | 42 | III | *3-30* | *2-15* | *3* | kappa | *3-15* | *1* | *IGHM* | 0 | 8 | 1 | 1.9 | 2 | 2.4 | 1.6 | 1 | | | | > E-04 | | |
| L1542x1762 | 42 | III | *3-33* | *3/OR15-3a* | *4* | kappa | *4-1* | *1* | *IGHM* | 0 | 8 | 6 | 1.5 | 1.2 | 1.3 | 1.6 | 11.2 | | | | 2.2E-06 | 5.6E-07 | 17.7 |
| L1542x1777 | 42 | III | *3-30* | *2-15* | *5* | kappa | *3-11* | *2* | *IGHM* | 0 | 1 | 0 | 1.7 | 1.6 | 2.3 | 4.5 | 1.1 | | > E-04 | | > E-04 | | |
| L1542x1914 | 42 | III | *3-33* | *6-13* | *3* | kappa | *3-15* | *1* | *IGHM* | 0 | 4 | 0 | 1.8 | 1.6 | 1.5 | 0.4 | 3.6 | | | | 1.1E-05 | | |
| D0751x0500 | 51 | II | *4-39* | *6-19* | *3* | lambda | *1-40* | *1* | *IGHM* | 16 | 8 | 3 | 7.8 | 4.4 | 5.7 | 0.4 | 1.6 | 1.7E-05 | 2.5E-06 | 5.7E-06 | > E-04 | | |
| D0851x1758 | 51 | II | *1-3* | *6-13* | *4* | lambda | *1-47* | *3* | *IGHM* | 2 | 16 | 6 | 1.4 | 1.6 | 1.8 | 0.4 | 1 | | | | > E-04 | | |
| D0751x0169 | 51 | III | *1-3* | *6-13* | *4* | lambda | *1-47* | *3* | *IGHG1* | 2 | 14 | 9 | 1.5 | 3.6 | 5.3 | 0.4 | 1.2 | | | 3.8E-06 | > E-04 | | |
| D0751x0322 | 51 | III | *1-3* | *6-13* | *4* | lambda | *1-47* | *3* | *IGHM* | 2 | 18 | 7 | 1.4 | 2.4 | 2.4 | 0.3 | 0.9 | | | | > E-04 | | |
| D0751x0901 | 51 | III | *1-3* | *6-13* | *4* | lambda | *1-47* | *2* | *IGHM* | 2 | 14 | 5 | 1.5 | 3.7 | 4.5 | 0.3 | 0.9 | | | 6.3E-06 | > E-04 | | |
| D0751x0968 | 51 | III | *1-3* | *6-13* | *4* | lambda | *1-47* | *2* | *IGHM* | 2 | 15 | 6 | 1.8 | 4.5 | 5 | 0.3 | 1.1 | | > E-04 | 6.0E-06 | > E-04 | | |
| D0851x1592 | 51 | III | *1-3* | *6-13* | *4* | lambda | *1-47* | *3* | *IGHM* | 2 | 11 | 4 | 1.9 | 3.6 | 5.4 | 0.3 | 1.2 | | | 7.1E-06 | > E-04 | | |
| D0851x1742 | 51 | III | *1-3* | *6-13* | *4* | lambda | *1-47* | *3* | *IGHM* | 2 | 18 | 7 | 1.5 | 1.9 | 1.7 | 0.4 | 1.1 | | | 7.7E-06 | > E-04 | | |
| D0851x1710 | 51 | III | *3-21* | *6-13* | *5* | lambda | *3-1* | *2* | *IGHG1* | 0 | 1 | 2 | 1.7 | 1.5 | 1.3 | 0.4 | 12 | | | | | 5.6E-08 | 1.1 |
| L1471x4237 | 71 | C | *4-4* | *2-15* | *6* | kappa | *3-20* | *2* | *IGHM* | 57 | 1 | 2 | 1.3 | 1.4 | 1.2 | 0.5 | 10.8 | | | | > E-04 | | 9.3 |
| L1471x3919 | 71 | III | *4-4* | *2-15* | *6* | kappa | *3-20* | *2* | *IGHM* | 57 | 1 | 2 | 1.7 | 1.6 | 1.4 | 0.5 | 10.9 | | | | > E-04 | 5.0E-07 | 7.6 |
| L1471x3710 | 71 | II | *3-7* | *3-3* | *3* | lambda | *1-51* | *2* | *IGHM* | 42 | 7 | 2 | 1.8 | 1.9 | 1.7 | 8.1 | 1.2 | | | | 7.0E-06 | | 71.6 |
| L1471x4112 | 71 | III | *3-7* | *3-3* | *3* | lambda | *1-51* | *2* | *IGHM* | 42 | 7 | 3 | 2 | 2 | 2.1 | 7.5 | 1.1 | | | | 6.0E-06 | | 69.4 |
| L1471x4493 | 71 | C | *3-49* | *6-13* | *4* | kappa | *3-20* | *1* | *IGHG1* | 38 | 5 | 1 | 10.5 | 11.9 | 11.6 | 11.4 | 4.1 | 2.6E-06 | 2.6E-09 | 5.2E-10 | 2.5E-09 | 5.1E-05 | 100 |
| L1471x4560 | 71 | C | *3-49* | *6-13* | *4* | kappa | *3-20* | *1* | *IGHM* | 38 | 6 | 3 | 3 | 5.5 | 11.3 | 11.4 | 2.8 | 4.2E-05 | 2.3E-06 | 5.0E-07 | 2.4E-06 | | 94.1 |
| L1471x4142 | 71 | III | *3-49* | *6-13* | *4* | kappa | *3-20* | *1* | *IGHM* | 38 | 1 | 0 | 3.2 | 2.7 | 10.7 | 13.5 | 2.3 | 3.0E-05 | 2.0E-06 | 2.3E-08 | 4.1E-08 | | 99.7 |
| L1471x4353 | 71 | C | *3-48* | *4-23* | *3* | kappa | *1-17* | *4* | *IGHM* | 36 | 10 | 1 | 2.7 | 2.8 | 2.3 | 5.6 | 1.7 | | | | 1.2E-05 | | 96.8 |
| L1471x4041 | 71 | III | *3-48* | *4-23* | *3* | kappa | *1-17* | *4* | *IGHG1* | 36 | 10 | 1 | 3.2 | 3 | 2.7 | 3.1 | 1.8 | | | | 1.5E-05 | | |
| L1471x4055 | 71 | III | *3-48* | *4-23* | *3* | kappa | *1-17* | *4* | *IGHA1* | 36 | 12 | 6 | 1.5 | 1.6 | 1.4 | 0.7 | 1.6 | | | | 2.4E-05 | | |
| L1471x4476 | 71 | C | *3-33* | *4-11* | *4* | kappa | *3-20* | *1* | *IGHG1* | 34 | 1 | 2 | 1.6 | 9.1 | 12.3 | 13.1 | 9.4 | | 4.4E-06 | 2.9E-07 | 3.7E-08 | 1.4E-06 | 100 |
| L1471x3945 | 71 | III | *3-33* | *4-11* | *4* | kappa | *3-20* | *1* | *IGHM* | 34 | 2 | 0 | 2.5 | 7 | 11.2 | 12 | 10 | | 5.2E-06 | 5.2E-07 | 4.4E-08 | 2.2E-06 | 100 |
| L1471x4092 | 71 | III | *3-33* | *4-11* | *4* | kappa | *3-20* | *1* | *IGHM* | 34 | 0 | 0 | 2.1 | 5.7 | 12.4 | 13.5 | 8.4 | | 4.5E-06 | 7.4E-07 | 6.2E-08 | 3.9E-05 | 100 |
| L1471x3246 | 71 | II | *3-33* | *4-11* | *6* | kappa | *3-20* | *2* | *IGHM* | 33 | 5 | 1 | 1.6 | 1.7 | 2.6 | 10.3 | 10.6 | | | | 2.7E-07 | 2.9E-07 | 74 |
| L1471x3374 | 71 | II | *3-33* | *4-11* | *6* | kappa | *3-20* | *2* | *IGHM* | 33 | 5 | 1 | 1.1 | 1 | 1.7 | 9.9 | 10.8 | | | 1.0E-05 | 2.9E-07 | 3.1E-07 | 72.5 |

| Antibody name | donor | Pf exposure | Ig gene information | | | | | | | | | | ELISA AUC | | | | | SPR KD (M) | | | | | Pf sporozoite Inhibition (%) |
|---|---|---|---|---|---|---|---|---|---|---|---|---|---|---|---|---|---|---|---|---|---|---|---|
| | | | IGHV | IGHD | IGHJ | light | IGKV/ IGLV | IGKJ/ IGLJ | IGHC | cluster id | IGHV SHM | IGKV/IGLV SHM | KQPA | NPDP | NVDP | NANP | C-CSP | KQPA | NPDP | NVDP | NANP | C-CSP | |
| L1471x3639 | 71 | II | 3-33 | 4-11 | 6 | kappa | 3-20 | 2 | IGHM | 33 | 8 | 1 | 2 | 2.2 | 5.6 | 11.4 | 10.3 | | | 1.3E-05 | 2.4E-07 | 3.4E-06 | 71.9 |
| L1471x3767 | 71 | II | 3-33 | 4-11 | 6 | kappa | 3-20 | 2 | IGHM | 33 | 14 | 5 | 1.4 | 1.3 | 1.7 | 10 | 11.2 | | | | 2.8E-07 | 1.7E-07 | 83.5 |
| L1471x3768 | 71 | II | 3-33 | 4-11 | 6 | kappa | 3-20 | 2 | IGHM | 33 | 8 | 3 | 1.3 | 1.4 | 1.6 | 8.8 | 10.8 | | | | 2.2E-07 | 3.1E-07 | 85.4 |
| L1471x3896 | 71 | III | 3-33 | 4-11 | 6 | kappa | 3-20 | 2 | IGHM | 33 | 7 | 5 | 1.7 | 1.8 | 5.1 | 11.7 | 10.6 | | 2.9E-05 | 5.7E-06 | 5.2E-08 | 1.6E-06 | 100 |
| L1471x3960 | 71 | III | 3-33 | 4-11 | 6 | kappa | 3-20 | 2 | IGHM | 33 | 10 | 3 | 1.7 | 2.6 | 9.2 | 11.9 | 10.5 | | 8.6E-06 | 6.2E-07 | 2.6E-09 | 7.6E-07 | 100 |
| L1471x4130 | 71 | III | 3-33 | 4-11 | 6 | kappa | 3-20 | 2 | IGHM | 33 | 7 | 2 | 1.8 | 2.1 | 7.1 | 13.4 | 11.2 | | | 2.8E-06 | 5.1E-08 | 1.3E-06 | 90.5 |
| L1471x4059 | 71 | III | 3-33 | 2-21 | 4 | kappa | 3-11 | 1 | IGHA1 | 32 | 8 | 5 | 1.3 | 1.4 | 1.3 | 9 | 10.2 | | | | 7.8E-06 | 7.4E-07 | 59.6 |
| L1471x4074 | 71 | III | 3-33 | 2-21 | 4 | kappa | 3-11 | 1 | IGHM | 32 | 1 | 1 | 1.2 | 1.4 | 1.3 | 6.7 | 8.9 | | | | 9.2E-06 | 2.2E-06 | 51.6 |
| L1471x3922 | 71 | III | 3-33 | 1-1 | 5 | kappa | 3-11 | 2 | IGHM | 31 | 2 | 3 | 1.6 | 1.8 | 2 | 11.5 | 1.3 | | | | 1.9E-06 | | 79.7 |
| L1471x3978 | 71 | III | 3-33 | 5-24 | 6 | kappa | 2-24 | 2 | IGHG1 | 29 | 5 | 2 | 1.5 | 1.5 | 1.5 | 9.6 | 1.1 | | | | 1.7E-05 | | 57.8 |
| L1471x4310 | 71 | C | 3-33 | 2-15 | 4 | kappa | 1-5 | 1 | IGHG1 | 27 | 0 | 2 | 1.7 | 4.7 | 11.4 | 13.5 | 1.4 | | 6.7E-06 | 8.4E-07 | 3.1E-08 | | 100 |
| L1471x3965 | 71 | III | 3-33 | 2-15 | 4 | kappa | 1-5 | 1 | IGHG1 | 27 | 0 | 2 | 2.4 | 5.9 | 11.4 | 11.9 | 1.3 | | 5.1E-06 | 9.3E-07 | 3.1E-08 | | 96.5 |
| L1471x3659 | 71 | II | 3-30 | 2-2 | 4 | lambda | 3-21 | 3 | IGHM | 26 | 7 | 5 | 1.8 | 1.4 | 1.5 | 11 | 1.1 | | | | 6.1E-07 | | 84.6 |
| L1471x3865 | 71 | III | 3-30 | 4-17 | 3 | kappa | 3-20 | 2 | IGHM | 25 | 3 | 1 | 2.6 | 2.1 | 1.9 | 11.6 | 1.8 | | | | 5.2E-07 | | 80.8 |
| L1471x3921 | 71 | III | 3-30 | 4-17 | 3 | kappa | 3-20 | 2 | IGHM | 25 | 5 | 1 | 3 | 2.3 | 2.9 | 11.3 | 1.7 | | | | 4.6E-07 | | 74.1 |
| L1471x4087 | 71 | III | 3-30 | 4-17 | 3 | kappa | 3-20 | 2 | IGHG3 | 25 | 3 | 1 | 2.8 | 2.5 | 2.4 | 13.5 | 2 | | | | 4.6E-07 | | 80.9 |
| L1471x4115 | 71 | III | 3-30 | 4-17 | 3 | kappa | 3-20 | 2 | IGHG1 | 25 | 4 | 1 | 3.6 | 3.6 | 3.2 | 13.3 | 2.5 | | | | 4.6E-07 | | 86.2 |
| L1471x4239 | 71 | C | 3-23 | 6-13 | 3 | kappa | 3-20 | 1 | IGHM | 16 | 15 | 5 | 1.7 | 2.1 | 2.1 | 0.6 | 3.6 | | | | > E-04 | | |
| L1471x4467 | 71 | C | 3-23 | 6-13 | 3 | kappa | 3-20 | 1 | IGHM | 16 | 14 | 5 | 1.3 | 1.4 | 2 | 0.5 | 2.7 | | | | > E-04 | | |
| L1471x4052 | 71 | III | 3-23 | 6-13 | 3 | kappa | 3-20 | 1 | IGHM | 16 | 13 | 4 | 2.2 | 2.3 | 2.5 | 0.7 | 4.1 | | | | > E-04 | | |
| L1471x4267 | 71 | C | 3-23 | 6-19 | 4 | lambda | 1-47 | 2 | IGHM | 14 | 2 | 0 | 2.2 | 13 | 13.2 | 12.8 | 1.7 | | 1.9E-06 | 5.5E-08 | 4.9E-08 | > E-04 | 100 |
| L1471x3426 | 71 | II | 3-23 | 6-19 | 4 | lambda | 1-47 | 1 | IGHM | 14 | 13 | 7 | 1.8 | 11.1 | 11.2 | 11.5 | 1.4 | | 2.6E-07 | 1.9E-08 | 1.2E-08 | | 100 |
| L1471x3933 | 71 | III | 3-23 | 6-19 | 4 | lambda | 1-47 | 1 | IGHM | 14 | 15 | 7 | 1.8 | 11.2 | 11.4 | 11.7 | 1.3 | | 2.6E-07 | 1.9E-08 | 1.7E-08 | | 97.9 |
| L1471x4264 | 71 | C | 3-15 | NA | 4 | kappa | 2-30 | 1 | IGHM | 8 | 12 | 2 | 2.9 | 11.7 | 12.9 | 12.9 | 1.7 | | 1.9E-06 | 9.2E-08 | 1.1E-07 | > E-04 | 98.9 |
| L1471x3870 | 71 | III | 3-15 | NA | 4 | kappa | 2-30 | 1 | IGHM | 8 | 11 | 1 | 2.3 | 11.3 | 11.5 | 11.4 | 1.5 | | 2.1E-06 | 1.0E-07 | 7.9E-08 | > E-04 | 96.9 |
| L1471x4067 | 71 | III | 3-15 | NA | 4 | kappa | 2-30 | 1 | IGHM | 8 | 12 | 1 | 2.9 | 12.8 | 13.4 | 13.2 | 2.2 | | 1.8E-06 | 9.4E-08 | 7.3E-08 | | 97.6 |
| L1471x3726 | 71 | II | 1-2 | 2-15 | 4 | kappa | 2-28 | 2 | IGHG1 | 2 | 6 | 5 | 1.9 | 5.4 | 10.2 | 0.7 | 1.5 | | 5.3E-06 | 6.1E-08 | 1.2E-05 | | 86.6 |
| L1471x4167 | 71 | III | 1-2 | 2-15 | 4 | kappa | 2-28 | 2 | IGHG1 | 2 | 6 | 5 | 1.8 | 7.3 | 11.5 | 0.7 | 1.6 | | 3.7E-06 | 6.5E-08 | 7.9E-06 | | 88.5 |
| L1471x4415 | 71 | C | 3-33 | 1-1 | 4 | kappa | 4-1 | 1 | IGHM | 0 | 0 | 0 | 0.8 | 0.8 | 0.9 | 9.2 | 4.2 | | | | 9.3E-07 | 1.1E-05 | 65.7 |
| L1471x4496 | 71 | C | 3-33 | 4-17 | 1 | kappa | 3-15 | 3 | IGHM | 0 | 0 | 0 | 0.7 | 1 | 1.1 | 6.3 | 9.4 | | | | 5.5E-06 | 1.9E-05 | 65.7 |
| L1471x4498 | 71 | C | 3-33 | 3-22 | 4 | kappa | 3-11 | 2 | IGHG1 | 0 | 9 | 2 | 1.4 | 8.6 | 11.3 | 11.4 | 1.2 | | 2.9E-06 | 7.0E-08 | 6.2E-10 | > E-04 | 93.3 |
| L1471x3269 | 71 | II | 3-33 | 3-22 | 4 | lambda | 2-14 | 1 | IGHM | 0 | 1 | 3 | 1.5 | 1.4 | 1.4 | 9.3 | 3.8 | | | | 1.0E-06 | | 69.9 |
| D0972x1572 | 72 | II | 4-59 | 3-16 | 3 | kappa | 2-28 | 2 | IGHM | 32 | 1 | 1 | 1.1 | 6 | 10.3 | 3.7 | 0.9 | | 2.7E-06 | 1.6E-07 | 9.8E-06 | | 81.2 |

| Antibody name | donor | Pf exposure | Ig gene information | | | | | | | | | | ELISA AUC | | | | | SPR KD (M) | | | | | Pf sporozoite Inhibition (%) |
|---|---|---|---|---|---|---|---|---|---|---|---|---|---|---|---|---|---|---|---|---|---|---|---|
| | | | IGHV | IGHD | IGHJ | light | IGKV/ IGLV | IGKJ/ IGLJ | IGHC | cluster id | IGHV SHM | IGKV/IGLV SHM | KQPA | NPDP | NVDP | NANP | C-CSP | KQPA | NPDP | NVDP | NANP | C-CSP | |
| D0972x1765 | 72 | II | 4-59 | 3-16 | 3 | kappa | 2-28 | 2 | IGHM | 32 | 0 | 0 | 1.2 | 7.8 | 10.4 | 2.8 | 1 | | 2.8E-06 | 1.7E-07 | 1.0E-05 | > E-04 | 76.8 |
| D0972x1985 | 72 | III | 4-59 | 3-16 | 3 | kappa | 2-28 | 2 | IGHM | 32 | 3 | 0 | 1.3 | 7.9 | 10.9 | 5.4 | 0.9 | | 2.8E-06 | 6.5E-08 | 9.9E-06 | | 74.7 |
| D0972x1994 | 72 | III | 4-59 | 3-16 | 3 | kappa | 2-28 | 2 | IGHM | 32 | 2 | 1 | 1.2 | 8.7 | 11.6 | 2 | 1 | | 2.2E-06 | 1.7E-07 | 9.5E-06 | | |
| D0972x2159 | 72 | III | 4-39 | 3-10 | 5 | lambda | 9-49 | 2 | IGHA1 | 28 | 18 | 10 | 12.1 | 13.1 | 11.9 | 9.4 | 6.2 | 9.3E-07 | 8.4E-09 | 3.8E-08 | 5.1E-08 | 4.7E-06 | 86.8 |
| D0972x2296 | 72 | III | 4-39 | 2-21 | 5 | lambda | 9-49 | 2 | IGHA1 | 28 | 22 | 15 | 11.6 | 12.3 | 11.5 | 8.1 | 9.5 | 1.6E-06 | 5.0E-09 | 5.3E-08 | 1.4E-07 | 2.1E-06 | 68.7 |
| D0972x2249 | 72 | III | 4-31 | 2-15 | 4 | kappa | 4-1 | 1 | IGHM | 25 | 8 | 3 | 1.9 | 2.3 | 2.1 | 1.5 | 0.6 | > E-04 | | | 8.4E-06 | | |
| D0972x2147 | 72 | III | 3-33 | 2-15 | 4 | kappa | 1-5 | 2 | IGHG1 | 17 | 3 | 1 | 0.8 | 4.6 | 12.1 | 12.5 | 5.9 | | 2.2E-06 | 5.0E-07 | 4.7E-09 | 5.6E-05 | 100 |
| D0972x2262 | 72 | III | 3-33 | 2-15 | 4 | kappa | 1-5 | 2 | IGHG1 | 17 | 3 | 2 | 1.3 | 2.8 | 9.4 | 11.7 | 3.1 | | 2.4E-06 | 5.6E-07 | 1.3E-07 | | 89.3 |
| D0972x1797 | 72 | II | 3-33 | 3-22 | 4 | kappa | 1-5 | 1 | IGHG1 | 16 | 0 | 0 | 1.1 | 1.7 | 1.7 | 12.9 | 1.4 | | | 6.1E-06 | 1.9E-08 | | 81.7 |
| D0972x2248 | 72 | III | 3-33 | 3-22 | 4 | kappa | 1-5 | 1 | IGHG1 | 16 | 1 | 1 | 1 | 1.2 | 1.4 | 10.5 | 1.2 | | > E-04 | 5.7E-06 | 1.8E-08 | | 80.8 |
| D0972x2292 | 72 | III | 3-33 | 3-22 | 4 | kappa | 1-5 | 1 | IGHM | 16 | 3 | 0 | 1.1 | 1.5 | 2.2 | 12.8 | 1.1 | | > E-04 | 1.9E-06 | 1.4E-08 | > E-04 | 97.5 |
| D0972x1888 | 72 | II | 3-33 | 2-15 | 4 | kappa | 1-5 | 1 | IGHG1 | 15 | 2 | 0 | 1.1 | 10.2 | 12.1 | 11.9 | 1.3 | | 3.7E-06 | 2.4E-07 | 4.6E-09 | > E-04 | 100 |
| D0972x2163 | 72 | III | 3-33 | 2-15 | 4 | kappa | 1-5 | 1 | IGHM | 15 | 0 | 0 | 1 | 7.6 | 12 | 12.1 | 1.2 | | 4.6E-06 | 3.0E-07 | 7.0E-09 | | 100 |
| D0972x2219 | 72 | III | 3-33 | 4-23 | 3 | kappa | 1-5 | 4 | IGHM | 14 | 3 | 1 | 0.7 | 6.8 | 9.3 | 12 | 0.7 | | 7.4E-06 | 1.2E-06 | 5.7E-09 | | 91.5 |
| D0972x2290 | 72 | III | 3-33 | 4-23 | 3 | kappa | 1-5 | 4 | IGHM | 14 | 0 | 0 | 1.2 | 1.4 | 1.3 | 10.6 | 0.6 | | > E-04 | | 1.6E-06 | | 52.9 |
| D0972x1987 | 72 | III | 3-33 | 4-17 | 3 | kappa | 1-5 | 3 | IGHM | 13 | 0 | 0 | 1.2 | 5.6 | 11 | 12.3 | 1.1 | | 4.7E-06 | 2.9E-07 | 2.2E-08 | | 100 |
| D0972x2100 | 72 | III | 3-33 | 4-17 | 3 | kappa | 1-5 | 3 | IGHG1 | 13 | 3 | 1 | 1.3 | 4.4 | 11.9 | 13.5 | 0.9 | | 3.1E-06 | 1.0E-07 | 6.3E-09 | | 86.7 |
| D0972x2151 | 72 | III | 3-33 | 2-15 | 2 | kappa | 4-1 | 1 | IGHG1 | 12 | 0 | 0 | 0.9 | 1.2 | 1.3 | 10.9 | 0.8 | | | | 5.0E-06 | | 55.8 |
| D0972x2235 | 72 | III | 3-33 | 2-15 | 2 | kappa | 4-1 | 1 | IGHM | 12 | 0 | 0 | 1.2 | 1.7 | 1.5 | 10 | 1.1 | | | | 5.2E-06 | | 57.5 |
| D0972x1694 | 72 | II | 3-30 | 2-15 | 4 | lambda | 9-49 | 1 | IGHM | 11 | 2 | 2 | 1.6 | 3 | 7.6 | 6 | 4.4 | | | 3.2E-06 | 6.5E-06 | 1.1E-05 | 71.3 |
| D0972x1791 | 72 | II | 3-30 | 2-8 | 4 | lambda | 9-49 | 1 | IGHM | 11 | 11 | 3 | 1 | 1.4 | 2.3 | 0.7 | 1.3 | | | 5.9E-06 | 1.0E-05 | | |
| D0972x1196 | 72 | I | 3-30-3 | 6-19 | 4 | lambda | 4-69 | 3 | IGHM | 9 | 2 | 2 | 1.9 | 3.4 | 9.6 | 1.6 | 1.2 | | > E-04 | 1.6E-06 | 4.5E-06 | | 40.7 |
| D0972x1713 | 72 | II | 3-30-3 | 6-19 | 4 | lambda | 4-69 | 3 | IGHM | 9 | 1 | 2 | 1.1 | 2.3 | 9.4 | 0.9 | 0.9 | | | 1.3E-06 | 5.8E-06 | | 52.6 |
| D0972x1846 | 72 | II | 3-30-3 | 6-19 | 4 | lambda | 4-69 | 3 | IGHM | 9 | 3 | 2 | 1.2 | 1.8 | 10.7 | 1.2 | 0.8 | | | 9.5E-07 | 6.9E-06 | | 64.2 |
| D0972x1928 | 72 | III | 3-30-3 | 6-19 | 4 | lambda | 4-69 | 3 | IGHM | 9 | 3 | 2 | 1 | 1.8 | 9.2 | 0.6 | 1 | | | 1.7E-06 | 4.6E-06 | | 26.7 |
| D0972x2031 | 72 | III | 3-30-3 | 6-19 | 4 | lambda | 4-69 | 3 | IGHM | 9 | 3 | 2 | 1.2 | 2.5 | 10.7 | 0.7 | 1 | | | 1.1E-06 | 6.6E-06 | | 34.6 |
| D0972x1576 | 72 | II | 3-30-3 | 1-14 | 3 | kappa | 3-11 | 1 | IGHM | 8 | 8 | 0 | 1.3 | 2.2 | 6.7 | 7.1 | 1.2 | | 3.1E-06 | 3.6E-06 | 9.4E-06 | | 71.7 |
| D0972x1593 | 72 | II | 3-30-3 | 1-26 | 3 | kappa | 3-11 | 4 | IGHM | 7 | 7 | 0 | 1 | 1.7 | 1.9 | 5 | 1.1 | | | 5.3E-06 | 9.9E-06 | | |
| D0972x1647 | 72 | II | 3-30-3 | 1-26 | 3 | kappa | 3-11 | 4 | IGHM | 7 | 2 | 1 | 1.4 | 2.1 | 6.2 | 6.9 | 1.1 | | 8.1E-06 | 6.8E-06 | 9.6E-06 | | 59.4 |
| D0972x1782 | 72 | II | 3-30-3 | 1-26 | 3 | kappa | 3-11 | 4 | IGHM | 7 | 7 | 1 | 1.3 | 2 | 1.9 | 1.6 | 1.2 | | | 6.2E-06 | 7.5E-06 | | |
| D0972x1783 | 72 | II | 3-30-3 | 1-26 | 3 | kappa | 3-11 | 4 | IGHM | 7 | 0 | 1 | 1.2 | 2 | 5.4 | 5.1 | 1.2 | | 6.2E-06 | 7.2E-06 | 8.6E-06 | | 53.4 |
| D0972x1547 | 72 | II | 3-33 | 2-2 | 3 | kappa | 1-5 | 1 | IGHM | 0 | 5 | 1 | 0.9 | 1.5 | 2.3 | 9.7 | 0.6 | | | | 1.9E-07 | | 66.1 |
| D0972x1788 | 72 | II | 3-30-3 | 1-26 | 5 | kappa | 3-11 | 1 | IGHM | 0 | 9 | 2 | 2.7 | 4.1 | 3.5 | 0.9 | 2.1 | | | | 7.5E-06 | | |

| Antibody name | donor | Pf exposure | Ig gene information | | | | | | | | | | ELISA AUC | | | | | SPR KD (M) | | | | | Pf sporozoite Inhibition (%) |
|---|---|---|---|---|---|---|---|---|---|---|---|---|---|---|---|---|---|---|---|---|---|---|---|
| | | | *IGHV* | *IGHD* | *IGHJ* | light | *IGKV/ IGLV* | *IGKJ/ IGLJ* | *IGHC* | cluster id | IGHV SHM | IGKV/IGLV SHM | KQPA | NPDP | NVDP | NANP | C-CSP | KQPA | NPDP | NVDP | NANP | C-CSP | |
| D0972x1834 | 72 | II | *3-33* | *3-22* | *4* | kappa | *1-5* | *2* | *IGHM* | 0 | 0 | 0 | 1.2 | 2 | 1.7 | 9.2 | 1.5 | | | | 4.6E-06 | | 36 |
| D0972x1949 | 72 | III | *3-33* | *5-24* | *3* | kappa | *1-5* | *1* | *IGHG1* | 0 | 4 | 0 | 1.6 | 4.9 | 10.8 | 11.9 | 1.3 | | > E-04 | 7.9E-07 | 2.6E-08 | | 96.7 |
| D0972x1954 | 72 | III | *3-33* | *3-22* | *4* | kappa | *1-5* | *1* | *IGHG1* | 0 | 7 | 3 | 1.1 | 2.1 | 9.6 | 12 | 0.9 | | 3.2E-06 | 3.0E-08 | 1.8E-08 | | 86.4 |
| D0972x2032 | 72 | III | *3-74* | *1-14* | *5* | kappa | *2-28* | *4* | *IGHM* | 0 | 0 | 0 | 1.8 | 3.5 | 6.9 | 10.3 | 1.7 | | | | 2.4E-06 | | |
| D0972x2103 | 72 | III | *3-33* | *2-8* | *3* | kappa | *1-5* | *2* | *IGHG1* | 0 | 5 | 0 | 1.5 | 10.6 | 12.4 | 13.4 | 1.4 | | 2.5E-06 | 1.5E-07 | 6.9E-10 | > E-04 | 100 |
| D0972x2140 | 72 | III | *3-33* | *3-22* | *5* | kappa | *1-5* | *1* | *IGHG1* | 0 | 6 | 1 | 7.9 | 13.8 | 13.5 | 12.5 | 8.2 | 2.1E-05 | 1.4E-06 | 3.5E-08 | 9.0E-10 | > E-04 | 100 |
| D0972x2164 | 72 | III | *3-33* | *4-23* | *3* | kappa | *1-5* | *1* | *IGHG1* | 0 | 2 | 1 | 5.7 | 12.7 | 12.5 | 12 | 1.4 | 3.1E-05 | 5.1E-08 | 9.4E-10 | 6.2E-10 | > E-04 | 100 |
| D0972x2243 | 72 | III | *3-33* | *2-2* | *4* | kappa | *1-5* | *3* | *IGHM* | 0 | 0 | 0 | 1.2 | 2.1 | 3 | 12.4 | 1.4 | | 1.5E-05 | 4.0E-06 | 1.2E-08 | > E-04 | 100 |
| D0873x0668 | 73 | III | *3-30-3* | *3-22* | *4* | kappa | *3-15* | *1* | *IGHM* | 29 | 7 | 3 | 0.7 | 1 | 1 | 0.4 | 0.8 | | | | | | |
| D0773x0982 | 73 | III | *3-23* | *4-11* | *4* | lambda | *1-47* | *2* | *IGHM* | 28 | 0 | 0 | 0.8 | 9.8 | 10.4 | 4 | 0.8 | | 3.4E-06 | 4.3E-07 | 2.4E-06 | | |
| D0873x0860 | 73 | III | *3-23* | *4-11* | *4* | lambda | *1-47* | *2* | *IGHM* | 28 | 1 | 0 | 0.8 | 10.7 | 10.9 | 7.2 | 0.9 | | 3.4E-06 | 4.8E-07 | 2.4E-06 | | 64 |
| D0773x1256 | 73 | III | *3-74* | *3-10* | *4* | lambda | *9-49* | *2* | *IGHM* | 24 | 0 | 0 | 1.8 | 2.8 | 2.2 | 12.2 | 5.8 | | | | 4.7E-08 | 2.4E-05 | 100 |
| D0873x0684 | 73 | III | *3-74* | *3-10* | *4* | lambda | *9-49* | *2* | *IGHM* | 24 | 0 | 0 | 1.4 | 2 | 1.8 | 13.4 | 2.8 | | > E-04 | > E-04 | 3.1E-08 | > E-04 | 100 |
| D0773x1096 | 73 | III | *3-7* | *3-16* | *4* | lambda | *4-69* | *3* | *IGHM* | 22 | 6 | 1 | 0.8 | 4 | 11.9 | 8.5 | 0.9 | | 6.1E-06 | 1.5E-07 | 4.5E-06 | | 78.1 |
| D0773x1233 | 73 | III | *3-7* | *3-16* | *4* | lambda | *4-69* | *3* | *IGHG1* | 22 | 9 | 1 | 1 | 3 | 12.2 | 8.5 | 0.7 | | 3.8E-06 | 4.4E-08 | 5.3E-07 | | 74 |
| D0773x1204 | 73 | III | *3-7* | *3-10* | *4* | kappa | *4-1* | *4* | *IGHM* | 21 | 2 | 2 | 1.1 | 1.9 | 7.2 | 0.4 | 1.2 | | | 5.2E-06 | | | 63.8 |
| D0773x1284 | 73 | III | *3-7* | *3-10* | *4* | kappa | *4-1* | *4* | *IGHM* | 21 | 1 | 2 | 0.9 | 2.4 | 11.6 | 0.6 | 1 | | 7.7E-06 | 2.0E-07 | 7.1E-06 | | |
| D0873x0588 | 73 | III | *3-7* | *3-10* | *4* | kappa | *4-1* | *4* | *IGHM* | 21 | 2 | 0 | 1.1 | 2.4 | 12.1 | 0.5 | 0.9 | | | 4.2E-08 | 3.4E-06 | | 41.3 |
| D0873x0615 | 73 | III | *3-7* | *3-10* | *4* | kappa | *4-1* | *4* | *IGHM* | 21 | 5 | 0 | 1.4 | 2.9 | 12.3 | 0.5 | 0.9 | | 6.2E-06 | 2.4E-08 | 3.2E-06 | | 45.3 |
| D0773x0980 | 73 | III | *3-49* | *3-10* | *4* | kappa | *1-5* | *1* | *IGHM* | 18 | 0 | 0 | 2.1 | 2.6 | 2.3 | 3.7 | 1.7 | | | | 3.4E-06 | | |
| D0773x1239 | 73 | III | *3-49* | *3-10* | *4* | kappa | *1-5* | *1* | *IGHM* | 18 | 0 | 0 | 2.6 | 3.1 | 2.7 | 4.6 | 1.9 | | | | 3.5E-06 | | |
| D0873x1096 | 73 | II | *3-49* | *2-15* | *6* | kappa | *2D-29* | *2* | *IGHM* | 17 | 0 | 0 | 6.6 | 1.9 | 2.7 | 13.2 | 1.1 | 9.6E-06 | 5.3E-06 | 7.4E-06 | 4.2E-07 | | |
| D0873x1308 | 73 | II | *3-49* | *2-15* | *6* | kappa | *2D-29* | *2* | *IGHM* | 17 | 0 | 0 | 4.1 | 0.8 | 0.9 | 10.3 | 0.8 | | 5.5E-06 | 5.6E-06 | 5.3E-07 | | 73 |
| D0873x0689 | 73 | III | *3-49* | *2-15* | *6* | kappa | *2D-29* | *2* | *IGHM* | 17 | 3 | 3 | 10.1 | 1.7 | 2.5 | 12.6 | 1.3 | 5.8E-06 | 3.7E-06 | 4.3E-06 | 2.2E-07 | | |
| D0773x1090 | 73 | III | *3-49* | *6-19* | *4* | kappa | *4-1* | *2* | *IGHM* | 16 | 1 | 0 | 2.3 | 2.2 | 3.1 | 12.7 | 1.2 | > E-04 | 5.7E-06 | 2.1E-06 | 6.6E-07 | | 76.8 |
| D0773x1259 | 73 | III | *3-49* | *6-19* | *4* | kappa | *4-1* | *2* | *IGHM* | 16 | 0 | 0 | 5.5 | 2.1 | 4.2 | 12 | 1.7 | 1.0E-05 | 4.8E-06 | 1.2E-06 | 5.0E-07 | | |
| D0773x1305 | 73 | III | *3-49* | *6-19* | *4* | kappa | *4-1* | *2* | *IGHG1* | 16 | 2 | 5 | 11.5 | 8.8 | 11.4 | 11.9 | 7 | 3.6E-06 | 1.3E-07 | 5.3E-08 | 3.8E-09 | 2.4E-05 | 95.2 |
| D0773x1216 | 73 | III | *3-48* | *3-10* | *4* | kappa | *2D-29* | *4* | *IGHA1* | 14 | 9 | 8 | 2.3 | 2.9 | 2.4 | 3.7 | 1.4 | | | 3.5E-06 | 5.3E-06 | | |
| D0873x0896 | 73 | III | *3-48* | *3-10* | *4* | kappa | *2D-29* | *4* | *IGHA1* | 14 | 9 | 8 | 1.9 | 2.2 | 2.2 | 3.2 | 1.3 | | | | 5.4E-06 | | |
| D0873x0800 | 73 | III | *3-33* | *3-22* | *4* | kappa | *3-11* | *4* | *IGHM* | 13 | 2 | 3 | 0.9 | 1.2 | 1.1 | 13.2 | 0.7 | | | | 2.5E-06 | | 60.2 |
| D0873x0647 | 73 | III | *3-33* | *3-22* | *6* | kappa | *1-5* | *1* | *IGHM* | 10 | 12 | 3 | 0.8 | 2.4 | 8.5 | 12.5 | 0.9 | | 3.2E-06 | 1.1E-07 | 2.0E-09 | | |
| D0773x1210 | 73 | III | *3-33* | *3-22* | *3* | kappa | *1-5* | *1* | *IGHM* | 9 | 3 | 4 | 1.1 | 8.9 | 10.5 | 11.9 | 1.1 | > E-04 | 5.8E-06 | 1.5E-06 | 1.1E-08 | > E-04 | 100 |
| D0773x1001 | 73 | III | *3-23* | *6-19* | *4* | lambda | *1-47* | *2* | *IGHM* | 7 | 0 | 0 | 0.9 | 11.5 | 11.7 | 2.6 | 0.8 | | | 3.8E-07 | 6.2E-06 | | 73.8 |

| Antibody name | donor | Pf exposure | Ig gene information | | | | | | | | | | ELISA AUC | | | | | SPR KD (M) | | | | | Pf sporozoite Inhibition (%) |
|---|---|---|---|---|---|---|---|---|---|---|---|---|---|---|---|---|---|---|---|---|---|---|---|
| | | | *IGHV* | *IGHD* | *IGHJ* | light | *IGKV/ IGLV* | *IGKJ/ IGLJ* | *IGHC* | cluster id | IGHV SHM | IGKV/IGLV SHM | KQPA | NPDP | NVDP | NANP | C-CSP | KQPA | NPDP | NVDP | NANP | C-CSP | |
| D0773x1006 | 73 | III | *3-23* | *6-19* | *4* | lambda | *1-47* | *2* | *IGHM* | 7 | 0 | 0 | 1 | 12.5 | 12.4 | 3.8 | 1 | | | 2.2E-07 | 5.1E-06 | | 78.1 |
| D0773x1015 | 73 | III | *3-23* | *6-13* | *4* | lambda | *1-47* | *2* | *IGHM* | 7 | 0 | 0 | 1 | 12.1 | 12.3 | 4 | 1 | | | 3.1E-07 | 5.0E-06 | | 85.9 |
| D0773x1199 | 73 | III | *3-23* | *6-19* | *4* | lambda | *1-47* | *2* | *IGHM* | 7 | 0 | 0 | 1 | 12 | 11.8 | 3.8 | 1.3 | | | 4.4E-07 | 5.9E-06 | | 81.8 |
| D0773x1340 | 73 | III | *3-23* | *6-19* | *4* | lambda | *1-47* | *2* | *IGHM* | 7 | 0 | 0 | 1.1 | 10.6 | 11.2 | 2.1 | 1 | | | 4.5E-07 | 6.0E-06 | | 81.2 |
| D0873x0829 | 73 | III | *3-23* | *6-19* | *4* | lambda | *1-47* | *2* | *IGHM* | 7 | 0 | 0 | 1 | 11.8 | 12 | 3.1 | 0.9 | | 3.1E-06 | 4.4E-07 | 6.2E-06 | | 88.7 |
| D0773x1444 | 73 | II | *3-23* | *2-2* | *6* | kappa | *1-5* | *4* | *IGHM* | 6 | 3 | 4 | 1.5 | 1.9 | 2.2 | 6.6 | 0.9 | | | | 2.2E-06 | | 34.3 |
| D0773x1366 | 73 | II | *3-23* | *4-11* | *4* | kappa | *1-5* | *5* | *IGHM* | 5 | 2 | 2 | 0.6 | 6.7 | 9.2 | 0.4 | 1 | | 6.1E-06 | 4.9E-07 | 9.3E-06 | | 53.7 |
| D0773x1368 | 73 | II | *3-23* | *3-10* | *4* | kappa | *1-5* | *5* | *IGHM* | 5 | 7 | 5 | 1.2 | 8.6 | 10.8 | 0.4 | 1.4 | | 3.5E-06 | 1.8E-07 | 8.3E-06 | | 62.1 |
| D0773x1401 | 73 | II | *3-23* | *4-11* | *4* | kappa | *1-5* | *5* | *IGHM* | 5 | 13 | 10 | 1.2 | 8.6 | 11.1 | 2.5 | 1.4 | | 1.6E-06 | 1.2E-07 | 4.7E-06 | | |
| D0773x1419 | 73 | II | *3-23* | *4-11* | *4* | kappa | *1-5* | *5* | *IGHM* | 5 | 12 | 11 | 0.8 | 1.4 | 2.5 | 0.4 | 1 | | | 4.3E-06 | 5.1E-06 | | |
| D0773x1463 | 73 | II | *3-23* | *4-11* | *4* | kappa | *1-5* | *5* | *IGHM* | 5 | 7 | 9 | 0.9 | 11 | 11.3 | 1.8 | 1 | | 1.7E-06 | 1.1E-07 | 6.4E-06 | | 81.1 |
| D0773x1551 | 73 | II | *3-23* | *3-9* | *4* | kappa | *1-5* | *5* | *IGHM* | 5 | 9 | 8 | 1 | 8.5 | 11.3 | 5.3 | 1.7 | | 1.4E-06 | 5.9E-08 | 3.4E-06 | | 74 |
| D0773x1628 | 73 | II | *3-23* | *4-11* | *4* | kappa | *1-5* | *5* | *IGHM* | 5 | 12 | 11 | 1.1 | 5.1 | 9.6 | 2.9 | 1.2 | | 1.8E-06 | 1.6E-07 | 4.1E-06 | | 82.8 |
| D0873x1035 | 73 | II | *3-23* | *3-22* | *4* | kappa | *1-5* | *5* | *IGHM* | 5 | 9 | 8 | 1 | 8.8 | 12.2 | 4.1 | 1.1 | | 1.4E-06 | 6.8E-08 | 3.1E-06 | | 85.3 |
| D0873x1097 | 73 | II | *3-23* | *4-11* | *4* | kappa | *1-5* | *5* | *IGHM* | 5 | 23 | 13 | 1.2 | 3.3 | 10.1 | 1.4 | 1.1 | | 4.1E-06 | 4.4E-07 | 6.5E-06 | | 59.2 |
| D0873x1264 | 73 | II | *3-23* | *4-11* | *4* | kappa | *1-5* | *5* | *IGHM* | 5 | 7 | 8 | 0.9 | 6.7 | 8.5 | 0.5 | 1.1 | | 2.1E-06 | 1.5E-07 | 6.3E-06 | | |
| D0873x1342 | 73 | II | *3-23* | *4-11* | *4* | kappa | *1-5* | *5* | *IGHM* | 5 | 6 | 8 | 1.2 | 7.7 | 10.8 | 3.9 | 1.4 | | 1.9E-06 | 1.9E-07 | 6.3E-06 | > E-04 | 86.4 |
| D0773x1041 | 73 | III | *3-23* | *3-22* | *4* | kappa | *1-5* | *5* | *IGHM* | 5 | 5 | 5 | 0.9 | 8.8 | 12.1 | 0.5 | 1.2 | | 4.0E-06 | 2.6E-07 | 8.0E-06 | | 65.5 |
| D0873x0706 | 73 | III | *3-23* | *3-22* | *4* | kappa | *1-5* | *5* | *IGHM* | 5 | 5 | 5 | 1.1 | 6.9 | 11.3 | 0.4 | 1.2 | | 3.7E-06 | 2.8E-07 | 8.7E-06 | | 51.9 |
| D0873x0898 | 73 | III | *3-23* | *4-11* | *4* | kappa | *1-5* | *5* | *IGHM* | 5 | 18 | 14 | 0.8 | 3.8 | 11.2 | 2.3 | 1 | | 2.3E-07 | 3.7E-08 | 2.4E-06 | | 83.4 |
| D0773x1427 | 73 | II | *3-23* | *2-2* | *4* | kappa | *1-5* | *1* | *IGHM* | 4 | 15 | 3 | 1 | 1.9 | 6.5 | 10.1 | 1 | | | 6.5E-06 | 4.5E-06 | | 59.4 |
| D0773x1450 | 73 | II | *3-23* | *2-2* | *4* | kappa | *1-5* | *1* | *IGHA1* | 4 | 17 | 10 | 1.3 | 2.5 | 6.6 | 11.6 | 1.1 | | 1.9E-05 | 9.0E-06 | 2.4E-06 | | 60.5 |
| D0873x1316 | 73 | II | *3-23* | *2-2* | *4* | kappa | *1-5* | *1* | *IGHM* | 4 | 15 | 6 | 0.8 | 0.8 | 2.5 | 11.9 | 0.7 | | | 9.8E-06 | 2.2E-06 | | 77.6 |
| D0873x1337 | 73 | II | *3-23* | *2-2* | *4* | kappa | *1-5* | *1* | *IGHM* | 4 | 17 | 3 | 0.7 | 0.7 | 0.7 | 6.2 | 0.7 | | | 5.3E-07 | 5.2E-06 | | 51.7 |
| D0773x1316 | 73 | III | *3-23* | *2-2* | *4* | kappa | *1-5* | *1* | *IGHM* | 4 | 18 | 6 | 0.9 | 1.9 | 6.9 | 9.7 | 0.8 | | | 8.5E-06 | 4.0E-06 | | 59 |
| D0773x1320 | 73 | III | *3-23* | *2-2* | *4* | kappa | *1-5* | *1* | *IGHM* | 4 | 13 | 6 | 0.8 | 1.5 | 4.6 | 11.3 | 0.8 | | | 8.7E-06 | 2.1E-06 | | 74.7 |
| D0773x1356 | 73 | III | *3-23* | *2-2* | *4* | kappa | *1-5* | *1* | *IGHM* | 4 | 20 | 5 | 0.7 | 1.2 | 2.2 | 8.8 | 0.8 | | | 7.7E-06 | 3.5E-06 | | 69.1 |
| D0873x0623 | 73 | III | *3-23* | *2-2* | *4* | kappa | *1-5* | *1* | *IGHM* | 4 | 21 | 8 | 1.3 | 3.3 | 8.3 | 10.2 | 0.7 | | 2.1E-05 | 7.0E-06 | 3.6E-06 | | 54.9 |
| D0873x0900 | 73 | III | *3-23* | *2-2* | *4* | kappa | *1-5* | *1* | *IGHM* | 4 | 15 | 4 | 0.7 | 1.1 | 3.1 | 10.9 | 0.7 | | | 7.9E-06 | 3.6E-06 | | 68.2 |
| D0873x0905 | 73 | III | *3-23* | *2-2* | *4* | kappa | *1-5* | *1* | *IGHM* | 4 | 16 | 2 | 0.7 | 0.9 | 1.1 | 9.1 | 0.7 | | | 9.7E-07 | 5.3E-06 | | 58.2 |
| D0873x0619 | 73 | III | *1-18* | *4-17* | *4* | kappa | *3-11* | *3* | *IGHM* | 1 | 13 | 2 | 2.9 | 2 | 1.6 | 1.1 | 0.9 | 1.2E-05 | | | 7.0E-06 | | |
| D0773x1505 | 73 | II | *3-23* | *2-2* | *6* | kappa | *1-5* | *1* | *IGHM* | 0 | 12 | 5 | 1.1 | 1.9 | 5.6 | 9.8 | 1 | | | | 1.0E-06 | | |
| D0773x1288 | 73 | III | *3-33* | *6-13* | *4* | kappa | *1-39* | *3* | *IGHG1* | 0 | 0 | 0 | 1.1 | 1.7 | 5.7 | 12.6 | 1 | | > E-04 | | 1.4E-07 | | 68.3 |

| Antibody name | donor | Pf exposure | Ig gene information | | | | | | | | | | ELISA AUC | | | | | SPR KD (M) | | | | | Pf sporozoite Inhibition (%) |
|---|---|---|---|---|---|---|---|---|---|---|---|---|---|---|---|---|---|---|---|---|---|---|---|
| | | | ***IGHV*** | ***IGHD*** | ***IGHJ*** | **light** | ***IGKV/ IGLV*** | ***IGKJ/ IGLJ*** | ***IGHC*** | **cluster id** | **IGHV SHM** | **IGKV/IGLV SHM** | **KQPA** | **NPDP** | **NVDP** | **NANP** | **C-CSP** | **KQPA** | **NPDP** | **NVDP** | **NANP** | **C-CSP** | |
| D0873x0614 | 73 | III | *3-33* | *3-22* | *4* | kappa | *1-5* | *1* | *IGHG1* | 0 | 1 | 1 | 1.4 | 4.2 | 9.7 | 12.2 | 1.5 | | 3.8E-06 | 3.5E-07 | 2.0E-09 | | |
| D0873x0922 | 73 | III | *3-33* | *6-13* | *4* | kappa | *1-12* | *4* | *IGHM* | 0 | 2 | 1 | 1.1 | 1.8 | 1.7 | 1.9 | 1.1 | | | | 5.9E-06 | | |

Pf exposures I, II, III and C refer to first, second, third and challenge Pf sporozoites exposures, respectively

cluster ids are unique within donors and 0 indicates non-clonally expanded single events

SHM: somatic hypermutations, represented in nucleotide base pairs

AUC: Area under ELISA curve

#### Table S2. Binding profiles of PfCSP-reactive antibodies isolated from CD19+CD27+CD38+PfCSP- plasmablasts.

| Antibody name | Ig gene information | | | | | | | ELISA AUC | | | | | SPR $K_D$ (M) | | | | Pf sporozoite inhibition (%) |
|---|---|---|---|---|---|---|---|---|---|---|---|---|---|---|---|---|---|
| | *IGHV* | *IGKV* | IgH CDR3 | Igκ/Igλ CDR3 | *IGHV* SHM | *IGKV/IGLV* SHM | *IGHC* | KQPA | NPDP | NVDP | NANP | C-CSP | KQPA | NPDP | NVDP | NANP | |
| D0972x2261 | 3-33 | 1-5 | ARPGAGSTGYVAFDI | QQYNSYWT | 4 | 0 | *IGHG1* | 2.6 | 11 | 12.7 | 11.5 | 2.5 | | 1.5E-06 | 2.4E-08 | < E-10 | 97.2 |
| L1372x1220 | 3-33 | 1-5 | AKVGEGQVGDSSGYYDH | QQYKSFWT | 6 | 1 | *IGHG1* | 7.7 | 11.6 | 12.6 | 11.6 | 4.6 | 2.1E-05 | 1.0E-06 | 5.6E-08 | 5.5E-10 | 96.8 |
| L1372x1240 | 3-33 | 1-5 | ARVALRATVVWGDAFDI | QQYNSYWT | 1 | 0 | *IGHG1* | 1.8 | 5.8 | 11.7 | 11.3 | 2.2 | | 4.2E-06 | 1.3E-06 | 3.6E-09 | 94.3 |
| L1372x1315 | 3-33 | 1-5 | ARVGDPEYYYDSSGFYDY | QQYNSYWT | 6 | 0 | *IGHG1* | 1.9 | 9.2 | 12 | 11.6 | 1.3 | | 3.8E-06 | 4.0E-07 | 3.4E-10 | 97.7 |
| L1372x2541 | 3-33 | 1-5 | ARVGDYSDFKYGAFDI | QQYNSYWT | 14 | 4 | *IGHG1* | 12.5 | 11.8 | 12.7 | 11.5 | 1.6 | 1.5E-05 | 1.9E-08 | < E-10 | 4.8E-10 | 97.3 |
| D0773x1333 | 3-33 | 1-5 | ARPKNYYESTGDYSGAFDI | QQYNSYWT | 6 | 0 | *IGHG1* | 1.7 | 1.9 | 3.7 | 11.4 | 2 | | 2.5E-05 | 8.3E-06 | 6.0E-08 | 89.2 |

SHM: somatic hypermutations, indicated in nucleotide basepairs

AUC: Area under ELISA curve

#### Table S3: Data collection and refinement statistics for *IGHV3-33*-encoded mAbs.

| | 3246-NANA | 2243-NANP$_5$ | 2541-NANP$_5$ | 3945-V$_H$H-NANP$_3$ | 4498-V$_H$H-NANP$_3$ | 4498-V$_H$H-NDN$_3$ |
|---|---|---|---|---|---|---|
| **Beamline** | CLS-08ID-1 | CLS-08ID-1 | APS-23-ID-B | NSLS-II-17-ID-2 | NSLS-II-17-ID-2 | CLS-08ID-1 |
| **Wavelength (Å)** | 1.03320 | 0.97949 | 1.033167 | 0.979360 | 0.979341 | 0.97949 |
| **Space group** | $P2_122_1$ | $P2_1$ | $P2_1$ | $P2_12_12$ | $P2_1$ | $P2_1$ |
| **Cell dimensions** | | | | | | |
| ***a,b,c*** **(Å)** | 67.1, 68.7, 104.2 | 79.1, 67.9, 100.4 | 66.8, 63.7, 146.7 | 156.2, 230.7, 48.2 | 63.0, 76.8, 63.7 | 61.5, 76.1, 63.2 |
| **α, β, γ (°)** | 90, 90, 90 | 90, 106.2, 90 | 90, 101.0, 90 | 90, 90, 90 | 90, 98.9, 90 | 90, 98.3, 90 |
| **Resolution (Å)[a]** | 40.0-1.80 (1.83-1.80) | 40.0-2.00 (2.03-2.00) | 40.0-2.55 (2.59-2.55) | 40.0-2.90 (2.95-2.90) | 40.0-1.40 (1.42-1.40) | 40.0-1.70 (1.73-1.70) |
| **No. molecules in ASU** | 1 | 1 | 1 | 3 | 1 | 1 |
| **No. observations** | 294,782 (14,082) | 228,124 (8,715) | 268,610 (11,187) | 590,243 (28,817) | 786,144 (33,433) | 423,380 (22,202) |
| **No. unique observations** | 45,439 (2,139) | 68,815 (2,909) | 39,583 (1,755) | 39,920 (1,998) | 117,560 (4,896) | 62,305 (3,120) |
| **Multiplicity** | 6.5 (6.6) | 3.3 (2.9) | 6.8 (6.3) | 14.8 (14.7) | 6.7 (6.8) | 6.7 (6.9) |
| **$R_{merge}$ (%)[b]** | 8.2 (71.7) | 13.9 (57.1) | 9.5 (74.9) | 24.2 (81.6) | 7.8 (62.5) | 4.5 (28.3) |
| **$R_{pim}$ (%)[c]** | 3.5 (30.2) | 8.9 (38.5) | 3.9 (32.2) | 6.5 (22.1) | 3.3 (26.0) | 1.9 (11.4) |
| **<I/σ I>** | 14.9 (1.5) | 6.5 (1.4) | 13.8 (1.5) | 10.0 (1.7) | 12.7 (1.6) | 26.5 (5.9) |
| **$CC_{½}$** | 99.9 (53.2) | 98.9 (54.2) | 99.8 (70.1) | 99.7 (66.4) | 99.8 (79.1) | 99.9 (97.8) |
| **Completeness (%)** | 100.0 (100.0) | 99.3 (98.0) | 99.4 (98.9) | 99.9 (100.0) | 99.9 (100.0) | 98.1 (96.9) |
| **Refinement Statistics** | | | | | | |
| **Reflections used in refinement** | 45,389 | 68,790 | 39,506 | 39,826 | 117,483 | 62,263 |
| **Reflections used for R-free** | 2,270 | 1,999 | 1,976 | 1,989 | 2,010 | 2,002 |
| **Non-hydrogen atoms** | 3,708 | 7,624 | 6,872 | 12,760 | 5,108 | 5,010 |
| **Macromolecule** | 3,307 | 6,699 | 6,779 | 12,757 | 4,404 | 4,375 |
| **Water** | 385 | 815 | 93 | 3 | 675 | 635 |
| **Heteroatom** | 16 | 110 | 0 | 0 | 29 | 0 |
| **$R_{work}$[d] / $R_{free}$[e]** | 16.3 / 20.7 | 18.5 / 22.6 | 20.4 / 25.6 | 21.0 / 26.5 | 16.1 / 18.6 | 15.8 / 18.4 |
| **Rms deviations from ideality** | | | | | | |
| **Bond lengths (Å)** | 0.010 | 0.003 | 0.004 | 0.002 | 0.02 | 0.008 |
| **Bond angle (°)** | 1.15 | 0.65 | 0.99 | 0.54 | 1.50 | 1.24 |
| **Ramachandran plot** | | | | | | |
| **Favoured regions (%)** | 97.9 | 97.0 | 96.9 | 96.6 | 97.7 | 98.1 |
| **Allowed regions (%)** | 2.1 | 3.0 | 2.9 | 3.4 | 2.3 | 1.9 |
| **B-factors (Å$^2$)** | | | | | | |
| **Wilson B-value** | 24.3 | 20.2 | 69.8 | 50.4 | 16.9 | 18.9 |
| **Average B-factors** | 32.9 | 28.1 | 86.5 | 49.0 | 28.5 | 26.3 |
| **Average macromolecule** | 31.7 | 26.6 | 86.7 | 49.0 | 26.6 | 24.6 |
| **Average heteroatom** | 45.3 | 55.1 | - | - | 67.8 | - |
| **Average water molecule** | 43.1 | 36.2 | 72.1 | 21.0 | 39.2 | 37.9 |

[a] Values in parentheses refer to the highest resolution bin.

[b] $R_{merge} = \Sigma_{hkl} \Sigma_i | I_{hkl,i} - \langle I_{hkl} \rangle | / \Sigma_{hkl} \langle I_{hkl} \rangle$

[c] $R_{pim} = \Sigma_{hkl} [1/(N-1)]^{1/2} \Sigma_i | I_{hkl,i} - \langle I_{hkl} \rangle | / \Sigma_{hkl} \langle I_{hkl} \rangle$

[d] $R_{work} = (\Sigma \, ||F_o| - |F_c||) / (\Sigma \, |F_o|)$ - for all data except as indicated in footnote e.

[e] 5% of data were used for the $R_{free}$ calculation

#### Table S4: Table of contacts between *IGHV3-33*-encoded mAbs and PfCSP subdomains.

| **Ab** | **3246** | **2243** | **2541** | **3945** | **4498** | **4498** | | | | | | | | |
|---|---|---|---|---|---|---|---|---|---|---|---|---|---|---|
| **Antigen** | **NANA (BSA Å$^{2}$)** | **NANP$_{5}$ (BSA Å$^{2}$)** | **NANP$_{5}$ (BSA Å$^{2}$)** | **NANP$_{3}$ (BSA Å$^{2}$)** | **NANP$_{5}$ (BSA Å$^{2}$)** | **NDN$_{3}$ (BSA Å$^{2}$)** | **Interaction** | **3246** | **2243** | **2541** | **3945** | **4498** | **4498** | **Total** |
| | **Val4 (50)** | | | **Ala2 (32)** | **Ala2 (23)** | **Ala2 (14)** | | | | | | | | |
| | **Asp5 (10)** | **Asn3 (26)** | **Asn3 (44)** | **Asn3 (17)** | **Asn3 (9)** | **Asn3 (17)** | | | | | | | | |
| | | Asn$^{N}$ | | | | | HB | | H-Tyr58$^{OH}$ | | | | | 1 |
| | **Glu6 (139)** | **Pro4 (130)** | **Pro4 (132)** | **Pro4 (89)** | **Pro4 (130)** | **Pro4 (126)** | | | | | | | | |
| | Glu$^{O}$ | | | | Pro$^{O}$ | Pro$^{O}$ | HB | K-Tyr96$^{OH}$ | | | | K-Tyr96$^{OH}$ | K-Tyr96$^{OH}$ | 3 |
| | Glu$^{O}$ | | | | | | HB | H-Asn100A$^{ND2}$ | | | | | | 1 |
| | | | | Pro$^{O}$ | | | HB | | | | His100B$^{NE2}$ | | | 1 |
| | **Asn7 (112)** | **Asn5 (138)** | **Asn5 (138)** | **Asn5 (136)** | **Asn5 (93)** | **Asn5 (123)** | | | | | | | | |
| | Asn$^{OD1}$ | Asn$^{OD1}$ | Asn$^{OD1}$ | | Asn$^{OD1}$ | Asn$^{OD1}$ | HB | K-Ser94$^{N}$ | K-Tyr94$^{N}$ | K-Tyr94$^{N}$ | | K-Trp94$^{N}$ | K-Trp94$^{N}$ | 5 |
| | Asn$^{ND2}$ | Asn$^{ND2}$ | Asn$^{ND2}$ | | Asn$^{ND2}$ | Asn$^{ND2}$ | HB | K-Tyr91$^{O}$ | K-Tyr91$^{O}$ | K-Tyr91$^{O}$ | | K-Arg91$^{O}$ | K-Arg91$^{O}$ | 5 |
| | Asn$^{ND2}$ | Asn$^{ND2}$ | Asn$^{ND2}$ | | Asn$^{ND2}$ | Asn$^{ND2}$ | HB | K-Ser94$^{O}$ | K-Tyr94$^{O}$ | K-Tyr94$^{O}$ | | K-Trp94$^{O}$ | K-Trp94$^{O}$ | 5 |
| | | | | Asn$^{ND2}$ | | | HB | | | | K-Ser93$^{O}$ | | | 1 |
| | | Asn$^{ND2}$ | | | | | HB | | H-Cys100D$^{O}$ | | | | | 1 |
| | | | | Asn$^{O}$ | | | HB | | | | H-Gly100A$^{N}$ | | | 1 |
| | | | Asn$^{O}$ | | | | HB | | | H-Tyr100C$^{N}$ | | | | 1 |
| | | | | | Asn$^{O}$ | Asn$^{O}$ | HB | | | | | H-Trp100G$^{NE1}$ | H-Trp100G$^{NE1}$ | 2 |
| | Asn$^{O}$ | | | | | | WMHB | K-Tyr32$^{OH}$ | | | | | | 1 |
| | Asn$^{O}$ | | | | | | WMHB | K-Tyr91$^{O}$ | | | | | | 1 |
| | Asn$^{O}$ | | | | | | WMHB | H-Asn100A$^{N}$ | | | | | | 1 |
| | **Ala8 (17)** | **Ala6 (17)** | **Ala6 (23)** | **Ala6 (30)** | **Ala6 (28)** | **Val6 (28)** | | | | | | | | |
| | | | | | Ala$^{O}$ | | WMHB | | | | | H-Trp52$^{NE1}$ | | 1 |
| | Ala$^{O}$ | | | | | | HB | H-Asn100A$^{ND2}$ | | | | | | 1 |
| | **Asn9 (27)** | **Asn7 (23)** | **Asn7 (56)** | **Asn7 (75)** | **Asn7 (66)** | **Asp7 (57)** | | | | | | | | |
| | | | | | | Asp$^{OD1}$ | HB | | | | | | H-His97$^{NE2}$ | 1 |
| | Asn$^{OD1}$ | | | | | | WMHB | H-Asn100A$^{OD1}$ | | | | | | 1 |
| | | | | Asn$^{ND2}$ | | | HB | | | | H-Gly96$^{O}$ | | | 1 |
| | | | | Asn$^{ND2}$ | | | HB | | | | H-Ser98$^{OG}$ | | | 1 |
| | | | | Asn$^{ND2}$ | | | HB | | | | H-Ser98$^{O}$ | | | 1 |
| | | Asn$^{O}$ | | | | | HB | | H-Trp52$^{NE1}$ | | | | | 1 |
| | **Ala10 (106)** | **Pro8 (120)** | **Pro8 (119)** | **Pro8 (125)** | **Pro8 (125)** | **Pro8 (123)** | | | | | | | | |
| | Ala$^{O}$ | Pro$^{O}$ | Pro$^{O}$ | Pro$^{O}$ | Pro$^{O}$ | Pro$^{O}$ | HB | H-Tyr52A$^{N}$ | H-Tyr52A$^{N}$ | H-His52A$^{N}$ | H-Tyr52A$^{N}$ | H-Tyr52A$^{N}$ | H-Tyr52A$^{N}$ | 6 |
| | Ala$^{N}$ | | | | | | HB | H-Asn100A$^{OD1}$ | | | | | | 1 |
| | **Asn11 (104)** | **Asn9 (88)** | **Asn9 (107)** | **Asn9 (90)** | **Asn9 (100)** | **Asn9 (103)** | | | | | | | | |
| | Asn$^{OD1}$ | Asn$^{OD1}$ | Asn$^{OD1}$ | Asn$^{OD1}$ | Asn$^{OD1}$ | Asn$^{OD1}$ | HB | H-Gly33$^{N}$ | H-Gly33$^{N}$ | H-Gly33$^{N}$ | H-Gly33$^{N}$ | H-Gly33$^{N}$ | H-Gly33$^{N}$ | 6 |
| | Asn$^{ND2}$ | Asn$^{ND2}$ | Asn$^{ND2}$ | | | | HB | H-Val95$^{O}$ | H-Ala95$^{O}$ | H-Val95$^{O}$ | | | | 3 |
| | | | | | Asn$^{ND2}$ | Asn$^{ND2}$ | HB | | | | | H-Asp95$^{OD1}$ | H-Asp95$^{OD1}$ | 2 |
| | Asn$^{ND2}$ | Asn$^{ND2}$ | | Asn$^{ND2}$ | | | HB | H-Gly96$^{O}$ | H-Leu96$^{O}$ | | H-Gly96$^{O}$ | | | 3 |
| | Asn$^{ND2}$ | | | | | | HB | H-Asp97$^{OD1}$ | | | | | | 1 |
| | **Ser12 (65)** | **Ala10 (66)** | **Ala10 (73)** | **Ala10 (52)** | **Ala10 (73)** | **Ala10 (66)** | | | | | | | | |
| | Ser$^{OG}$ | | | | | | HB | H-Gly31$^{O}$ | | | | | | 1 |
| | Ser$^{N}$ | Ala$^{N}$ | | Ala$^{N}$ | Ala$^{N}$ | Ala$^{N}$ | HB | H-Gly31$^{O}$ | H-Ser31$^{O}$ | | H-Ser31$^{O}$ | H-Ile31$^{O}$ | H-Ile31$^{O}$ | 5 |
| | | | **Asn11 (7)** | **Asn11 (0)** | **Asn11 (9)** | **Asn11 (22)** | | | | | | | | |
| | | | **Pro12 (0)** | **Pro12 (59)** | **Pro12 (10)** | **Pro12 (18)** | | | | | | | | |
| **H-bonds** | 14 | 11 | 7 | 10 | 10 | 10 | | | | | | | | |
| **BSA Total (Å2)** | 630 | 608 | 699 | 705 | 666 | 697 | | | | | | | | |

#### Table S5: Data collection and refinement statistics for 4493 mAb.

| | 4493-$V_HH$-KQPA | 4493-$V_HH$-NPDP | 4493-$V_HH$-NDN$_3$ | 4493-$V_HH$-DND$_3$ | 4493-$V_HH$-NANP$_3$ |
|---|---|---|---|---|---|
| **Beamline** | APS-23ID-D | APS-23ID-B | APS-23ID-D | APS-23ID-D | NSLS-II-17-ID-2 |
| **Wavelength (Å)** | 1.033200 | 1.033167 | 1.033200 | 1.033200 | 0.979329 |
| **Space group** | $P2_12_12_1$ | $P2_12_12_1$ | P1 | P1 | $P2_1$ |
| **Cell dimensions** | | | | | |
| ***a,b,c*** **(Å)** | 77.0, 94.7, 144.0 | 54.5, 79.8, 281.8 | 82.7, 90.7, 92.5 | 83.1, 93.4, 94.2 | 64.0, 68.1, 84.6 |
| **α, β, γ (°)** | 90, 90, 90 | 90, 90, 90 | 89.4, 64.3, 76.9 | 82.7, 76.9, 64.5 | 90, 102.4, 90 |
| **Resolution (Å)**[a] | 30-1.93 (1.96-1.93) | 30-2.4 (2.44-2.4) | 30-2.15 (2.18-2.15) | 30-2.1 (2.13-2.1) | 30- 2.02 (2.05-2.02) |
| **No. molecules in ASU** | 2 | 2 | 4 | 4 | 1 |
| **No. observations** | 1,041,011 (45,384) | 656,780 (33,011) | 441,691 (18,037) | 502,150 (19,722) | 316,197 (12,374) |
| **No. unique observations** | 79,786 (3,549) | 49,310 (2,353) | 124,730 (5,069) | 141,733 (5,870) | 46,495 (1,934) |
| **Multiplicity** | 13.0 (12.8) | 14.0 (14.0) | 3.5 (3.5) | 3.5 (3.3) | 6.7 (6.4) |
| **$R_{merge}$ (%)**[b] | 18.7 (81.7) | 15.1 (83.0) | 7.0 (54.4) | 6.9 (53.8) | 11.6 (58.2) |
| **$R_{pim}$ (%)**[c] | 5.4 (23.9) | 4.3 (23.0) | 4.4 (33.9) | 5.3 (34.9) | 4.8 (24.2) |
| **<I/σ I>** | 9.4 (1.6) | 12.0 (1.5) | 10.5 (1.7) | 9.4 (1.7) | 9.7 (2.1) |
| **$CC_{½}$** | 99.7 (65.0) | 99.7 (76.1) | 99.7 (72.2) | 99.7 (64.1) | 99.5 (74.1) |
| **Completeness (%)** | 99.9 (100) | 99.9 (100) | 97.7 (96.9) | 97.7 (96.7) | 99.0 (87.4) |
| **Refinement Statistics** | | | | | |
| **Reflections (work)** | 79,565 | 49,165 | 124,663 | 141,701 | 46,490 |
| **Reflections (test)** | 1,530 | 1,549 | 1,357 | 1,995 | 1,998 |
| **Non-hydrogen atoms** | 9,453 | 8,768 | 17,873 | 18,502 | 4,810 |
| **Macromolecule** | 8,492 | 8,637 | 17,120 | 17,139 | 4,328 |
| **Water** | 937 | 119 | 681 | 1,327 | 441 |
| **Heteroatom** | 24 | 12 | 72 | 36 | 41 |
| **$R_{work}$[d] / $R_{free}$[e]** | 18.8 / 21.9 | 21.0 / 24.0 | 19.3 / 22.1 | 17.7 / 21.0 | 16.2 / 19.5 |
| **Rms deviations from ideality** | | | | | |
| **Bond lengths (Å)** | 0.004 | 0.002 | 0.003 | 0.003 | 0.004 |
| **Bond angle (°)** | 0.72 | 0.55 | 0.59 | 0.66 | 0.72 |
| **Ramachandran plot** | | | | | |
| **Favoured regions (%)** | 98.6 | 97.8 | 98.3 | 98.5 | 99.1 |
| **Allowed regions (%)** | 1.4 | 2.2 | 1.7 | 1.5 | 0.9 |
| **B-factors (Å$^2$)** | | | | | |
| **Wilson B-value** | 25.3 | 45.2 | 39.7 | 31.7 | 30.3 |
| **Average B-factors** | 30.4 | 60.1 | 54.7 | 44.8 | 34.3 |
| **Average macromolecule** | 29.5 | 60.2 | 54.8 | 44.6 | 33.0 |
| **Average heteroatom** | 55.8 | 62.5 | 65.5 | 78.3 | 56.8 |
| **Average water molecule** | 37.7 | 52.3 | 51.7 | 46.4 | 44.5 |

[a] Values in parentheses refer to the highest resolution bin.

[b] $R_{merge} = \Sigma_{hkl} \Sigma_i \mid I_{hkl,i} - <I_{hkl}> \mid / \Sigma_{hkl} <I_{hkl}>$

[c] $R_{pim} = \Sigma_{hkl} [1/(N-1)]^{1/2} \Sigma_i \mid I_{hkl,i} - <I_{hkl}> \mid / \Sigma_{hkl} <I_{hkl}>$

[d] $R_{work} = (\Sigma \mid |F_o| - |F_c| \mid) / (\Sigma \mid |F_o| )$ - for all data except as indicated in footnote e.

[e] 5% of data were used for the $R_{free}$ calculation

#### Table S6: Table of contacts between 4493 mAb and PfCSP subdomains.

| **Ab** | **4493** | **4493** | **4493** | **4493** | **4493** | | | |
|---|---|---|---|---|---|---|---|---|
| **Antigen** | **KQPA (BSA Å$^2$)** | **NPDP (BSA Å$^2$)** | **NDN$_3$ (BSA Å$^2$)** | **DND$_3$ (BSA Å$^2$)** | **NANP$_3$ (BSA Å$^2$)** | **Interaction** | **4493** | **Total** |
| | | **Pro2 (18)** | | **Val2 (82)** | **Ala2 (73)** | | | |
| | | **Asp3 (38)** | | **Asp3 (7)** | **Asn3 (9)** | | | |
| | | Asp$^{OD1}$ | | | | HB | K-Ser94$^{N}$ | 1 |
| | **Gly6 (30)** | **Pro4 (1)** | | **Pro4 (2)** | **Pro4 (4)** | | | |
| | | Pro$^{O}$ | | Pro$^{O}$ | Pro$^{O}$ | HB | H-Arg52$^{NH2}$ | 3 |
| | **Asn7 (38)** | **Asn5 (74)** | **Asn1 (59)** | **Asn5 (81)** | **Asn5 (81)** | | | |
| | Asn$^{N}$ | | Asn$^{N}$ | | | HB | H-Glu58$^{OE1}$ | 2 |
| | | Asn$^{ND2}$ | | | | HB | H-Glu58$^{OE2}$ | 1 |
| | Asn$^{O}$ | Asn$^{O}$ | Asn$^{O}$ | | | HB | H-Arg52$^{NH2}$ | 3 |
| | **Pro8 (121)** | **Ala6 (79)** | **Ala2 (99)** | **Ala6 (70)** | **Ala6 (86)** | | | |
| | Pro$^{O}$ | Ala$^{O}$ | Ala$^{O}$ | Ala$^{O}$ | Ala$^{O}$ | HB | K-Trp96$^{NE1}$ | 5 |
| | Pro$^{O}$ | Ala$^{O}$ | Ala$^{O}$ | Ala$^{O}$ | Ala$^{O}$ | HB | H-Trp100A$^{NE1}$ | 5 |
| | **Asp9 (35)** | **Asn7 (36)** | **Asn3 (38)** | **Asn7 (29)** | **Asn7 (30)** | | | |
| | Asp$^{O}$ | Asn$^{O}$ | Asn$^{O}$ | Asn$^{O}$ | Asn$^{O}$ | HB | H-Arg52$^{NE}$ | 5 |
| | | Asn$^{O}$ | | Asn$^{O}$ | | HB | H-Arg52$^{NH2}$ | 2 |
| | **Pro10 (122)** | **Pro8 (125)** | **Pro4 (121)** | **Pro8 (123)** | **Pro8 (122)** | | | |
| | Pro$^{O}$ | | Pro$^{O}$ | Pro$^{O}$ | Pro$^{O}$ | WMHB | H-Ser52A$^{N}$ | 4 |
| | **Asn11 (110)** | **Asn9 (117)** | **Asn5 (111)** | **Asn9 (113)** | **Asn9 (105)** | | | |
| | Asn$^{OD1}$ | | Asn$^{OD1}$ | Asn$^{OD1}$ | Asn$^{OD1}$ | WMHB | H-Ala33$^{N}$ / H-Val95$^{O}$ | 4 |
| | Asn$^{ND2}$ | Asn$^{ND2}$ | Asn$^{ND2}$ | Asn$^{ND2}$ | Asn$^{ND2}$ | HB | H-Glu96$^{O}$ | 5 |
| | Asn$^{ND2}$ | Asn$^{ND2}$ | Asn$^{ND2}$ | Asn$^{ND2}$ | Asn$^{ND2}$ | HB | H-Gly98$^{O}$ | 5 |
| | **Ala12 (19)** | **Val10 (7)** | **Val6 (10)** | **Val10 (11)** | **Ala10 (14)** | | | |
| | Ala$^{O}$ | Val$^{O}$ | Val$^{O}$ | Val$^{O}$ | Ala$^{O}$ | HB | H-Arg52$^{NE}$ | 5 |
| | **Asn13 (46)** | **Asp11 (44)** | **Asp7 (54)** | **Asp11 (55)** | **Asn11 (52)** | | | |
| | Asn$^{ND2}$ | | | | | WMHB | H-Ala52$^{O}$ | 1 |
| | | | Asp$^{N}$ | | Asn$^{N}$ | HB | H-Asn53$^{OD1}$ | 2 |
| | **Pro14 (31)** | **Pro12 (32)** | **Pro8 (33)** | **Pro12 (29)** | **Pro12 (35)** | | | |
| | **Asn15 (27)** | **Asn13 (12)** | | | | | | |
| | Asn$^{OD1}$ | | | | | WMHB | H-Arg56$^{NH2}$ | 1 |
| **H-bonds** | 8 | 11 | 9 | 8 | 8 | | | |
| **BSA Total (Å2)** | 579 | 583 | 525 | 602 | 611 | | | |

HB: hydrogen bond (3.89 Å cut-off)
WMHB: water-mediated hydrogen bond (3.89 Å cut-off)

#### Table S7. $IC_{50}$ values of representative potent antibodies.

| Antibody name | Ig gene information | | | | | SPR $K_D$ (M) | | | | | Traversal assay | |
|---|---|---|---|---|---|---|---|---|---|---|---|---|
| | *IGHV* | Light chain | *IGKV/ IGLV* | IgH CDR3 | Igκ/Igλ CDR3 | KQPA | NPDP | NVDP | NANP | C-CSP | Pf sporozoite inhibition (%) | $IC_{50}$ (µg/ml) |
| L1471x3870 | 3-15 | κ | 2-30 | TTASFDY | MQGTHWPFST | | 2.1E-06 | 1.0E-07 | 7.9E-08 | > E-04 | 96.9 | >20 |
| L1471x4267 | 3-23 | λ | 1-47 | AKDVDSGWPSYFDY | AAWDDSLSGLV | | 1.9E-06 | 5.5E-08 | 4.9E-08 | > E-04 | 100 | 4.7 |
| D0972x2243 | 3-33 | κ | 1-5 | ARALDDASSTSCCALDY | QQYNSYFT | | 1.5E-05 | 4.0E-06 | 1.2E-08 | > E-04 | 100 | 0.2 |
| D0972x2292 | 3-33 | κ | 1-5 | ARVGDGNYYDSSGYYDY | QQYNSLWT | | > E-04 | 1.9E-06 | 1.4E-08 | > E-04 | 97.5 | 0.3 |
| D0972x2103 | 3-33 | κ | 1-5 | ARVLGEDCNGGVCSCCAFDI | QQYNSYYT | | 2.5E-06 | 1.5E-07 | 6.9E-10 | > E-04 | 100 | 0.2 |
| D0972x1888 | 3-33 | κ | 1-5 | ARVQTTTGGGSCCPFDY | QQYNSYWT | | 3.7E-06 | 2.4E-07 | 4.6E-09 | > E-04 | 100 | 0.3 |
| D0773x1210 | 3-33 | κ | 1-5 | ARVRDSSDYYGDAFDI | QQYNNYWT | > E-04 | 5.8E-06 | 1.5E-06 | 1.1E-08 | > E-04 | 100 | 0.4 |
| D0972x2140 | 3-33 | κ | 1-5 | AKVGEGQVGDSSGYYDH | QQYKSFWT | 2.1E-05 | 1.4E-06 | 3.5E-08 | 9.0E-10 | | 100 | 0.2 |
| D0972x2164 | 3-33 | κ | 1-5 | ARVGVVTPGGACCAFDI | QQYNSYWT | 3.1E-05 | 5.1E-08 | 9.4E-10 | 6.2E-10 | > E-04 | 100 | 0.5 |
| D0972x2147 | 3-33 | κ | 1-5 | ARTAQFSGGSCCHFDY | QQYNTSYT | | 2.2E-06 | 5.0E-07 | 4.7E-09 | 5.6E-05 | 100 | 0.5 |
| L1372x2541 | 3-33 | κ | 1-5 | ARVGDYSDFKYGAFDI | QQYNSYWT | 1.5E-05 | 1.9E-08 | < E-10 | 4.8E-10 | | 97.3 | 0.7 |
| L1471x3896 | 3-33 | κ | 3-20 | ARVGDGDGNYMDL | QQYGSLYT | | 2.9E-05 | 5.7E-06 | 5.2E-08 | 1.6E-06 | 100 | 3.9 |
| L1471x4476 | 3-33 | κ | 3-20 | ARAGYSNYGHYFDY | QQYGSSPPT | | 4.4E-06 | 2.9E-07 | 3.7E-08 | 1.4E-06 | 100 | 0.3 |
| L1471x4493 | 3-49 | κ | 3-20 | TRVELGSSWSLGY | QQYGSSPWT | 2.6E-06 | 2.6E-09 | 5.2E-10 | 2.5E-09 | | 100 | 0.4 |
| D0773x1305 | 3-49 | κ | 4-1 | TRVLYSSGWYPFDY | QQYYSIPYT | 3.6E-06 | 1.3E-07 | 5.3E-08 | 3.8E-09 | 2.4E-05 | 95.2 | 0.6 |
| D0873x0684 | 3-74 | λ | 9-49 | ARVDDYYGSGSYGLDY | GADHGSGRV | | >E-04 | > E-04 | 3.1E-08 | > E-04 | 100 | >20 |

#### Table S8. *In vivo* potency of *IGVH3-33*-encoded mAbs at 300 µg in protection against mosquito bite challenge infection.

**A. Protection and prepatency status of mice immunized with different antibodies**

| | mAbs | 1210 | | | | 2164 | | | | 4476 | | | | 1710 | | | |
|---|---|---|---|---|---|---|---|---|---|---|---|---|---|---|---|---|---|
| Independent experiments | Mouse | Serum conc. (µg/ml) | Protection | Prepatency | Inf. bites | Serum conc. (µg/ml) | Protection | Prepatency | Inf. bites | Serum conc. (µg/ml) | Protection | Prepatency | Inf. bites | Serum conc. (µg/ml) | Protection | Prepatency | Inf. bites |
| Experiment 1 | | 1210 (n=6) | | | | 2164 (n=6) | | | | 4476 (n=6) | | | | 1710 (n=4) | | | |
| | Mouse_1 | 40 | NP | 5 | 7 | 35 | P | | 3 | 36 | NP | 4 | 6 | 49 | NP | 4 | 7 |
| | Mouse_2 | 57 | P | | 2 | 36 | NP | 6 | 4 | 36 | P | | 3 | 20 | NP | 4 | 5 |
| | Mouse_3 | 24 | NP | 5 | 4 | 46 | NP | 5 | 6 | 38 | NP | 4 | 5 | 98 | NP | 4 | 3 |
| | Mouse_4 | 56 | P | | 4 | 54 | P | | 3 | 37 | P | | 4 | 54 | NP | 4 | 3 |
| | Mouse_5 | 43 | P | | 3 | 56 | P | | 4 | 34 | NP | 5 | 3 | | | | |
| | Mouse_6 | 55 | P | | 4 | ND | NP | 5 | 7 | 45 | P | | 3 | | | | |
| Experiment 2 | | 1210 (n=6) | | | | 2164 (n=6) | | | | 4476 (n=6) | | | | 1710 (n=5) | | | |
| | Mouse_1 | 24 | P | | 5 | 60 | NP | 4 | 5 | 46 | NP | 4 | 4 | 92 | NP | 4 | 2 |
| | Mouse_2 | 53 | NP | 5 | 4 | 40 | NP | 5 | 4 | 42 | NP | 4 | 2 | 61 | NP | 3 | 5 |
| | Mouse_3 | 65 | NP | 5 | 5 | 34 | NP | 4 | 3 | 51 | NP | 6 | 4 | 42 | NP | 4 | 5 |
| | Mouse_4 | 70 | NP | 4 | 4 | 37 | P | | 3 | 52 | NP | 5 | 4 | 49 | NP | 4 | 7 |
| | Mouse_5 | 92 | NP | 4 | 7 | 57 | NP | 5 | 5 | 97 | NP | 5 | 6 | 49 | NP | 3 | 4 |
| | Mouse_6 | 58 | NP | 5 | 4 | 64 | NP | 4 | 3 | 47 | NP | 5 | 2 | | | | |
| Experiment 3 | | 1210 (n=6) | | | | 2164 (n=6) | | | | 4476 (n=5) | | | | 1710 (n=6) | | | |
| | Mouse_1 | 84 | P | | 1 | 54 | NP | 5 | 2 | 64 | NP | 4 | 2 | 79 | NP | 5 | 2 |
| | Mouse_2 | 69 | P | | 2 | 59 | P | | 2 | 57 | NP | 3 | 3 | 65 | NP | 4 | 2 |
| | Mouse_3 | 78 | P | | 2 | 76 | NP | 3 | 2 | 56 | NP | 5 | 2 | 61 | NP | 4 | 2 |
| | Mouse_4 | 95 | NP | 5 | 2 | 72 | NP | 3 | 4 | 60 | P | | 2 | 70 | NP | 4 | 3 |
| | Mouse_5 | 76 | P | | 1 | 60 | NP | 4 | 3 | 61 | P | | 1 | 70 | NP | 4 | 1 |
| | Mouse_6 | 91 | P | | 2 | 72 | P | | 1 | | | | | 70 | NP | 4 | 2 |

Protection: P and NP indicate protected and non-protected mice, respectively
Prepatency: Number of days post infection when parasitemia was detected
Inf. bites: Number of *Pb-PfCSP* infected blood-fed mosquitoes

**B. P values of statistical analysis performed using Mantel-Cox log-rank test**

| mAbs | 1210 (n=18) | 2164 (n=18) | 4476 (n=17) |
|---|---|---|---|
| 1710 (n=16) | <0.0001 | 0.0013 | 0.0010 |
| 4476 (n=17) | 0.0740 | 0.8709 | |
| 2164 (n=18) | 0.1124 | | |

#### Table S9. *In vivo* potency of non-*IGVH3-33*-encoded mAbs at 300 µg in protection against mosquito bite challenge infection.

**A. Protection and prepatency status of mice immunized with different antibodies**

| | mAbs | 1210 | | | | 4493 | | | | CIS43 | | | | 317 | | | | 1710 | | | |
|---|---|---|---|---|---|---|---|---|---|---|---|---|---|---|---|---|---|---|---|---|---|
| **Independent experiments** | **Mouse** | **Serum conc. (µg/ml)** | **Protection** | **Prepatency** | **Inf. bites** | **Protection** | **Prepatency** | **Inf. bites** | **Inf. bites** | **Serum conc. (µg/ml)** | **Protection** | **Prepatency** | **Inf. bites** | **Serum conc. (µg/ml)** | **Protection** | **Prepatency** | **Inf. bites** | **Serum conc. (µg/ml)** | **Protection** | **Prepatency** | **Inf. bites** |
| **Experiment 1** | | **1210 (n=6)** | | | | **4493 (n=6)** | | | | **CIS43 (n=6)** | | | | **317 (n=6)** | | | | **1710 (n=6)** | | | |
| | Mouse_1 | 69 | NP | 4 | 5 | 66 | NP | 5 | 2 | 91.8 | P | | 2 | 74 | P | | 3 | 91 | NP | 4 | 3 |
| | Mouse_2 | 98 | NP | 5 | 3 | 56 | P | | 4 | 167 | P | | 4 | 78 | P | | 3 | 80 | NP | 5 | 2 |
| | Mouse_3 | 88 | NP | 4 | 3 | 56 | P | | 2 | 139.7 | P | | 3 | 73 | P | | 3 | 102 | NP | 4 | 5 |
| | Mouse_4 | 105 | NP | 5 | 4 | 55 | P | | 2 | 123 | P | | 4 | 87 | P | | 5 | 88 | NP | 4 | 3 |
| | Mouse_5 | 71 | P | | 4 | 57 | NP | 6 | 2 | 115 | P | | 3 | 87 | P | | 5 | 85 | NP | 4 | 3 |
| | Mouse_6 | 102 | NP | 4 | 2 | 58 | P | | 5 | 124 | P | | 4 | 107 | P | | 3 | 94 | NP | 5 | 4 |
| **Experiment 2** | | **1210 (n=6)** | | | | **4493 (n=6)** | | | | **CIS43 (n=6)** | | | | **317 (n=5)** | | | | **1710 (n=6)** | | | |
| | Mouse_1 | 98 | P | | 3 | 58 | P | | 4 | | | | | 80 | P | | 1 | 91 | NP | 4 | 3 |
| | Mouse_2 | 93 | P | | 3 | ND | P | | 1 | 115.8 | P | | 2 | 65 | P | | 2 | 89 | NP | 4 | 3 |
| | Mouse_3 | 76 | P | | 2 | 44 | P | | 2 | 113.4 | NP | 6 | 1 | 76 | P | | 3 | 95 | P | | 3 |
| | Mouse_4 | 82 | P | | 2 | 21 | P | | 1 | 91 | P | | 2 | 100 | P | | 2 | 114 | NP | 4 | 1 |
| | Mouse_5 | 67 | NP | 5 | 2 | 57 | P | | 4 | 95 | P | | 2 | 128 | P | | 2 | 111 | NP | 4 | 2 |
| | Mouse_6 | 81 | P | | 4 | 30 | P | | 1 | 122 | P | | 1 | | | | | 109 | NP | 4 | 1 |

Protection: P and NP indicate protected and non-protected mice, respectively
Prepatency: Number of days post infection when parasitemia was detected
Inf. bites: Number of *Pb-PfCSP* infected blood-fed mosquitoes
ND: Not determined

**B. P values of statistical analysis performed using Mantel-Cox log-rank test**

| **mAbs** | **1210 (n=12)** | **4493 (n=12)** | **CIS43 (n=12)** | **317 (n=11)** |
|---|---|---|---|---|
| **1710 (n=12)** | 0.0108 | <0.0001 | <0.0001 | <0.0001 |
| **317 (n=11)** | 0.0078 | 0.1663 | 0.3173 | |
| **CIS43 (n=12)** | 0.0294 | 0.5745 | | |
| **4493 (n=12)** | 0.0656 | | | |

#### Table S10. *In vivo* potency of non-*IGVH3-33*-encoded mAbs at 150 µg in protection against mosquito bite challenge infection.

**A. Protection and prepatency status of mice immunized with different antibodies**

| | **mAbs** | **1210** | | | | **4493** | | | | **CIS43** | | | | **317** | | | | **1710** | | | |
|---|---|---|---|---|---|---|---|---|---|---|---|---|---|---|---|---|---|---|---|---|---|
| **Independent experiments** | **Mouse** | **Serum conc. (µg/ml)** | **Protection** | **Prepatency** | **Inf. bites** | **Protection** | **Prepatency** | **Inf. bites** | **Inf. bites** | **Serum conc. (µg/ml)** | **Protection** | **Prepatency** | **Inf. bites** | **Serum conc. (µg/ml)** | **Protection** | **Prepatency** | **Inf. bites** | **Serum conc. (µg/ml)** | **Protection** | **Prepatency** | **Inf. bites** |
| **Experiment 1** | | **1210 (n=6)** | | | | **4493 (n=6)** | | | | **CIS43 (n=6)** | | | | **317 (n=6)** | | | | **1710 (n=6)** | | | |
| | Mouse_1 | 67 | P | | 3 | 27 | P | | 2 | 68.6 | P | | 5 | 66 | P | | 2 | 66 | NP | 4 | 3 |
| | Mouse_2 | 84 | NP | 4 | 2 | 29 | NP | 5 | 4 | 90.9 | NP | 6 | 5 | 58 | P | | 4 | 66 | NP | 5 | 2 |
| | Mouse_3 | 76 | NP | 6 | 4 | 29 | NP | 5 | 2 | 63.3 | P | | 3 | 60 | P | | 2 | 55 | NP | 5 | 1 |
| | Mouse_4 | 65 | NP | 4 | 5 | 30 | P | | 4 | | | | | 55 | P | | 3 | 42 | NP | 4 | 2 |
| | Mouse_5 | 62 | NP | 4 | 4 | 43 | P | | 3 | 72 | NP | 4 | 5 | 59 | P | | 2 | 57 | NP | 4 | 2 |
| | Mouse_6 | 73 | P | | 3 | 32 | P | | 2 | 68 | P | | 3 | 53.7 | P | | 3 | 63 | NP | 4 | 2 |
| **Experiment 2** | | **1210 (n=6)** | | | | **4493 (n=6)** | | | | **CIS43 (n=8)** | | | | **317 (n=6)** | | | | **1710 (n=6)** | | | |
| | Mouse_1 | 76 | NP | 4 | 4 | 46 | P | | 4 | 60.5 | P | | 4 | 63 | P | | 1 | 58 | NP | 3 | 4 |
| | Mouse_2 | 78 | NP | 10 | 4 | 55 | P | | 3 | 57.1 | P | | 2 | 82 | NP | 6 | 3 | 63 | NP | 4 | 4 |
| | Mouse_3 | 65 | NP | 5 | 3 | 28 | NP | 5 | 3 | 58.2 | P | | 2 | 66 | P | | 2 | 57 | NP | 4 | 3 |
| | Mouse_4 | 65 | NP | 4 | 6 | 20 | NP | 5 | 4 | 55 | NP | 5 | 5 | 68 | P | | 3 | 60 | NP | 4 | 4 |
| | Mouse_5 | 62 | NP | 3 | 5 | 36 | NP | 5 | 4 | 63 | P | | 2 | 60 | P | | 4 | 63 | NP | 5 | 3 |
| | Mouse_6 | 69 | NP | 6 | 3 | 46 | P | | 3 | 48 | P | | 3 | 60.8 | NP | 5 | 3 | 49 | NP | 4 | 2 |
| | Mouse_7 | | | | | | | | | 56.1 | N | 6 | 3 | | | | | | | | |
| | Mouse_8 | | | | | | | | | 67.4 | N | 4 | 3 | | | | | | | | |

Protection: P and NP indicate protected and non-protected mice, respectively
Prepatency: Number of days post infection when parasitemia was detected
Inf. bites: Number of *Pb-PfCSP* infected blood-fed mosquitoes

**B. P values of statistical analysis performed using Mantel-Cox log-rank test**

| **mAbs** | **1210 (n=12)** | **4493 (n=12)** | **CIS43 (n=14)** | **317 (n=12)** |
|---|---|---|---|---|
| **1710 (n=12)** | 0.0421 | <0.0001 | <0.0001 | <0.0001 |
| **317 (n=12)** | 0.0007 | 0.1751 | 0.2171 | |
| **CIS43 (n=14)** | 0.0180 | 0.9525 | | |
| **4493 (n=12)** | 0.0188 | | | |

#### Table S11. *In vivo* potency of non-*IGVH3-33* and *IGHV3*-33-encoded mAbs at 150 µg in protection against mosquito bite challenge infection.

**A. Protection and prepatency status of mice immunized with different antibodies**

| | mAbs | 1210 | | | | 4493 | | | | CIS43 | | | | 317 | | | | 2541 | | | | 1710 | | | |
|---|---|---|---|---|---|---|---|---|---|---|---|---|---|---|---|---|---|---|---|---|---|---|---|---|---|
| Independent experiments | Mouse | Serum conc. (µg/ml) | Protection | Prepatency | Inf. bites | Protection | Protection | Prepatency | Inf. bites | Serum conc. (µg/ml) | Protection | Prepatency | Inf. bites | Serum conc. (µg/ml) | Protection | Prepatency | Inf. bites | Serum conc. (µg/ml) | Protection | Prepatency | Inf. bites | Serum conc. (µg/ml) | Protection | Prepatency | Inf. bites |
| Experiment 1 | | 1210 (n=6) | | | | 4493 (n=6) | | | | CIS43 (n=6) | | | | 317 (n=6) | | | | 2541 (n=6) | | | | 1710 (n=6) | | | |
| | Mouse_1 | 58 | P | | 3 | 26 | P | | 1 | 60.5 | P | | 1 | 51 | P | | 3 | 64.1 | NP | 5 | 4 | 53 | NP | 5 | 1 |
| | Mouse_2 | 71 | P | | 3 | 27 | P | | 3 | 56.3 | NP | 5 | 2 | 50 | P | | 1 | 65.4 | P | | 1 | 66 | NP | 4 | 2 |
| | Mouse_3 | 59 | NP | 6 | 1 | 19 | P | | 1 | 55.1 | P | | 3 | 58 | P | | 4 | 59.6 | P | | 2 | 56 | NP | 4 | 3 |
| | Mouse_4 | 73 | P | | 3 | 25 | P | | 2 | 51.3 | NP | 6 | 3 | 63 | P | | 3 | 68.5 | NP | 5 | 4 | 50 | NP | 4 | 2 |
| | Mouse_5 | 60 | P | | 1 | 22 | NP | 5 | 3 | 74 | P | | 5 | 59 | P | | 5 | 106 | P | | 5 | 52 | P | | 2 |
| | Mouse_6 | 70 | NP | 5 | 2 | 26 | P | | 2 | 59 | P | | 2 | 57.9 | P | | 4 | 57 | P | | 2 | 58 | NP | 4 | 4 |
| Experiment 2 | | 1210 (n=6) | | | | 4493 (n=6) | | | | CIS43 (n=6) | | | | 317 (n=5) | | | | 2541 (n=6) | | | | 1710 (n=5) | | | |
| | Mouse_1 | 63 | NP | 4 | 3 | 37 | P | | 1 | 60.3 | P | | 1 | 59 | P | | 2 | 58.8 | P | | 3 | 71 | P | | 3 |
| | Mouse_2 | 84 | P | | 3 | 32 | P | | 2 | 61.2 | P | | 3 | 75 | P | | 4 | 53.9 | P | | 5 | 75 | NP | 5 | 3 |
| | Mouse_3 | 68 | P | | 2 | 39 | P | | 4 | 56.3 | P | | 1 | 56 | P | | 3 | 59.4 | P | | 1 | 688 | NP | 5 | 3 |
| | Mouse_4 | 71 | P | | 4 | 15 | P | | 3 | 61.6 | P | | 3 | 60 | P | | 2 | 47.9 | P | | 3 | 77 | NP | 4 | 3 |
| | Mouse_5 | 73 | P | | 2 | 37 | P | | 1 | 68 | P | | 4 | 54.5 | P | | 2 | 48 | P | | 1 | 73 | P | | 2 |
| | Mouse_6 | 78 | NP | 5 | 6 | 32 | P | | 2 | 66 | P | | 3 | | | | | 65 | P | | 4 | | | | |
| Experiment 3 | | 1210 (n=6) | | | | 4493 (n=6) | | | | CIS43 (n=6) | | | | 317 (n=6) | | | | 2541 (n=6) | | | | 1710 (n=6) | | | |
| | Mouse_1 | 61 | P | | 3 | 45 | P | | 1 | 68.4 | P | | 2 | 44 | P | | 6 | 45.9 | P | | 5 | 66 | NP | 4 | 4 |
| | Mouse_2 | 63 | P | | 4 | 45 | P | | 4 | 60.7 | P | | 3 | 54 | P | | 4 | 49.6 | P | | 4 | 70 | NP | 4 | 3 |
| | Mouse_3 | 64 | P | | 2 | 39 | P | | 6 | 75.3 | NP | 7 | 5 | 47 | P | | 2 | 47.5 | P | | 3 | 77.7 | NP | 4 | 5 |
| | Mouse_4 | 67 | P | | 2 | 29 | P | | 4 | 57.2 | P | | 3 | 65 | P | | 5 | 49.7 | P | | 4 | 67 | NP | 3 | 3 |
| | Mouse_5 | 57 | NP | 5 | 4 | 34 | P | | 2 | 74 | P | | 3 | 69 | P | | 4 | 55 | NP | 5 | 5 | 77 | NP | 5 | 2 |
| | Mouse_6 | 56 | NP | 4 | 5 | 27 | P | | 5 | 75 | P | | 2 | 66.6 | P | | 1 | 56 | P | | 3 | 57 | NP | 3 | 3 |

Protection: P and NP indicate protected and non-protected mice, respectively
Prepatency: Number of days post infection when parasitemia was detected
Inf. bites: Number of *Pb-PfCSP* infected blood-fed mosquitoes

**B. P values of statistical analysis performed using Mantel-Cox log-rank test**

| mAbs | 1210 (n=18) | 4493 (n=18) | CIS43 (n=18) | 317 (n=17) | 2541 (n=18) |
|---|---|---|---|---|---|
| 1710 (n=17) | 0.0010 | <0.0001 | <0.0001 | <0.0001 | <0.0001 |
| 2541 (n=18) | 0.2263 | 0.2956 | 0.9682 | 0.0827 | |
| 317 (n=17) | 0.0100 | 0.3311 | 0.0830 | | |
| CIS43 (n=18) | 0.2097 | 0.3065 | | | |
| 4493 (n=18) | 0.0359 | | | | |
